# Impacts of Cell Ranger versions on Chromium gene expression data

**DOI:** 10.1101/2024.08.10.607413

**Authors:** Imad Abugessaisa, Akira Hasegawa, Scott Walker, Shintaro Katayama, Juha Kere, Takeya Kasukawa

## Abstract

In droplet-based single cell gene expression data, cell barcode processing by Cell Ranger (CR) is a standard pipeline. But no systematic evaluation of the impact of CR version on single cell gene expression data has been conducted. To comprehensively evaluate the impact of CR version, we considered six molecular quality criteria, quantified gene expression, and performed downstream analysis for 12 single-cell datasets. Each dataset was processed by 15 versions of CR. We demonstrated that different versions of CR yield different numbers of cell barcodes with significant variation in detected UMIs, features, molecular qualities and average gene expression of protein-coding and lncRNA for the same dataset. Our analysis finds distinction between two diverse categories of cell barcodes: common barcodes unmasked by all versions of CR, and specific barcodes only unmasked/masked by some versions. Surprisingly, we observed variation in molecular read-out between common cell barcodes when called by different versions of CR. The specific barcodes yield skewed gene body coverage and form distinct clusters. The choice of CR version affects scores for quality, average gene expression, clustering results, and top cluster marker genes of the dataset.

## Introduction

The introduction of high-throughput RNA barcoding technologies, inDrop^1^ and Dropseq^2^ revolutionized the scRNA-seq field and enabled tracking of the cell-of-origin of mRNA to quantify gene expression levels of the entire set of genes in tens of thousands of cells^3^. The technology uses the cell barcode (nucleic acid sequences of about 10–16⍰bp long) as a label to uniquely identify and track each cell *in silico*. The cell barcodes are introduced into all RNA molecules of a cell, and unique barcodes are given to each cell in a given sample; in this way researchers can process large numbers of cells in parallel^3^. Indeed, microfluidic droplet technology enabled multiplexing has allowed for a plethora of large-scale single cell projects to profile mammalian cells and study complex biological systems, such as the <u>Human Cell Atlas, Tabula Sapiens</u>, and <u>Tabula Muris</u>.

Microfluidic droplet technologies have become more readily available and affordable. The computational challenge became how to process and identify ‘real’ cell reads, further complicated by the reliability of identifying doublet free cell-barcodes (singlets) and excluding problematic droplets. Problematic droplets include droplets containing two or more cells (doublets) resulting in excessive read numbers for a barcode, assumed to be a single cell. Another type of problematic droplet is those without cells at all (empty droplets)^4,5^. In addition to the above errors, another quality issue in microfluidic droplet technologies (droplet capture process) is the presence of extracellular mRNA molecules, or ambient RNA, that impacts the level of background RNA counts^6,7^. Poor quality cells are associated with low numbers of UMI counts and detected features, and high numbers of reads originating from mitochondrial genes (mitGenes)^8^. Gene body coverage of the cell has been used as an indication of the cell quality as well^9^. In this study we refer to detected numbers of UMIs, detected number of genes, percentage of mitGenes, ambient RNA contamination, doublets scores, and gene body coverage as metrics of molecular quality of the barcoded mRNA reads.

10x Genomics has developed <u>Cell Ranger</u> (CR), a software pipeline for the processing of Chromium Single Cell Gene Expression, immune and fixed RNA profiling^1^. The software uses Chromium cellular barcodes for cell calling to generate feature-barcode matrices. Several versions of CR have been released with the aim to enhance the molecular qualities, increase the number of identified cell-barcoded reads and support different kits. While CR development continue to release new versions addressing technical issues or recommend technical setting (e.g. inclusion of intronic read for CR v5.0.0 and v6.0.0, recommend specific version of transcriptomic reference), we noticed that recently published scRNA-seq datasets have used an old version of CR without any explanation or justification for such choice^10-13^. Although CR is widely used as a cell barcode calling pipeline in scRNA-seq studies (Chromium single cell gene expression data), two studies benchmarked two versions of CR; You and colleagues investigated the performance of v6.0.0^14^, and Rich and colleagues in a preprint compared CR v7 vs. v6^15^ output when used with Seurat^8^ and SCANPY^16^ (the dominant scRNA-seq analysis pipelines for R and Python programming languages, respectively).

In a technical note 10x Genomics (CR development team), investigated the mechanisms for the detection of intronic and antisense reads in 10x Genomics gene expression dataset and the impact of intronic and antisense reads on analysis interpretation of gene expression data (Interpreting Intronic and Antisense Reads in 10x Genomics Single Cell Gene Expression Data); primarily, Chromium single cell gene expression data include exonic read. The technical note pointed to the increase in number of UMI counts when including the intronic read, therefore, from CR v7.0.0 intronic reads are included by default in Chromium gene expression data, and 10x Genomics recommended the inclusion of intronic read in v5.0.0 and v6.0.0. Furthermore, the technical note from 10x Genomics (Interpreting Single Cell Gene Expression Data With and Without Intronic Reads) evaluated the impact of including intronic reads by analyzing the dataset PBMC 10K by using CR v6.1.2 with intronic reads and without intronic reads.

The above technical note has investigated the impact of inclusion of intronic UMI and impact of intronic read on total UMI counts, however, no independent systematic evaluation has been conducted to evaluate the impact of CR version choice on the resulting Chromium single cell gene expression dataset, in terms of molecular qualities, quantification of gene expression, and downstream analysis of Chromium single cell gene expression data.

## Results

### Analysis dataset and study design

To investigate and determine the impact of CR cell barcode calling pipeline on 10x Genomics scRNA-seq dataset, we used 12 published human and mouse datasets (Supplementary Table 1) and developed the workflow shown in Fig. 1 and (**METHODS**). The datasets studied cover a wide range of biological systems (whole-blood, cell lines, brain cell nuclei and whole neurons, tissues), processed by different methods (fixed and unfixed). The raw sequence reads (FASTQ files) from each dataset were aligned and quantified by fifteen versions of CR, resulting in 180 datasets with a total number of 702,493 cell barcodes (Fig. 1b and **METHODS**). The molecular qualities of the 180 datasets were evaluated by four state-of-the-art scRNA-seq methods (Fig. 1c and **METHODS**). Number of detected UMI, number of detected gene species, and percentage of mitGenes determined by Seurat 4.3.0^8^; doublets scores estimated by Scrublet^17^; ratio of ambient RNA contamination determined by decontX^7^; and gene body coverage of each cell barcode determined by SkewC^9^. The 180 datasets were processed by Seurat 4.3.0 to determine the impact on clustering analysis, dimensionality reduction, marker gene stability, and average gene expression.

**Figure 1.**
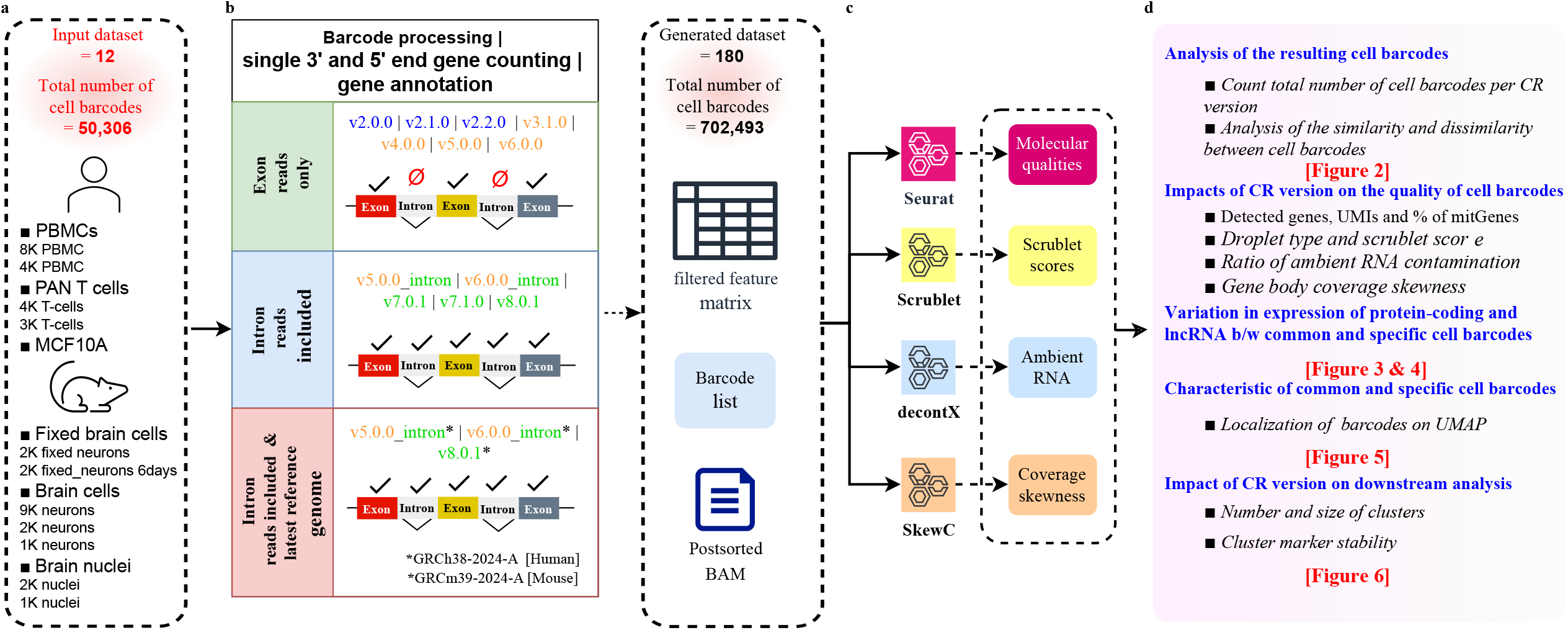
Overview of the computation workflow. **a**. Diverse collection of human and mouse cell lines, primary cell and tissue single-cell gene expression chromium data selected. Human datasets (n=5) were from mammary epithelial cells or peripheral blood cells. The mouse datasets (n=7) were either neurons or nuclei cells. All datasets were generated with Single Cell 3’ v2 protocol. **b**. Cell barcodes for each of the 12 datasets were called 15 times by different versions of Cell Ranger (CR). By default, CR results count exon reads in all versions. We followed recommendations from 10x Genomics and include intron reads and used latest reference genome. Each output of CR version contains post sorted BAM file, feature matrix and metrics summary files among other outputs. The output from each version of CR was used as input for the scRNA-seq QC method. **c**. Four QC methods were used to measure six types of quality criteria. **d**. Summary of the analysis and results.

### Different versions of CR yields different number of cell barcodes

We counted the number of barcodes in the resulting filtered_feature_bc_matrix (output from each version of CR). We observed that processing a dataset with different versions of CR yielded a different number of cell barcodes (Fig. 2a, Supplementary Table 1). There was variation in the number of cell barcodes between each version of CR, but the major changes were observed for v2.1.0 and v6.0.0 where the number of cell barcodes were decreased for all datasets, except for the dataset 9k Brain Cells from an E18 Mouse (neuron_9k), in which the number of cell barcodes were increased (8,965 cell barcodes from v2.0.0 to 9,128 cell barcodes from v2.1.0 and for the dataset 2K brain nuclei from E18 Mouse in v6.0.0). For v3.1.0 we observed a sizable increase in the number of detected cell barcodes for all human and mouse datasets. in the cell barcode calling algorithm in v3.0.0 (V3.1.0 is the oldest available version after the V3.0.0 update). Since v7.1.0 the CR development team restricted the auto-estimated expect-cells upper range in EmptyDrops arguments (as explained in the <u>release note</u> by CR development team). Also, starting from v7.0.0 the intronic reads were included by default. The inclusion of the intronic reads in v5.0.0 and v6.0.0 with the use of the latest reference transcriptome increases the number of detected cell barcodes for all datasets.

**Figure 2.**
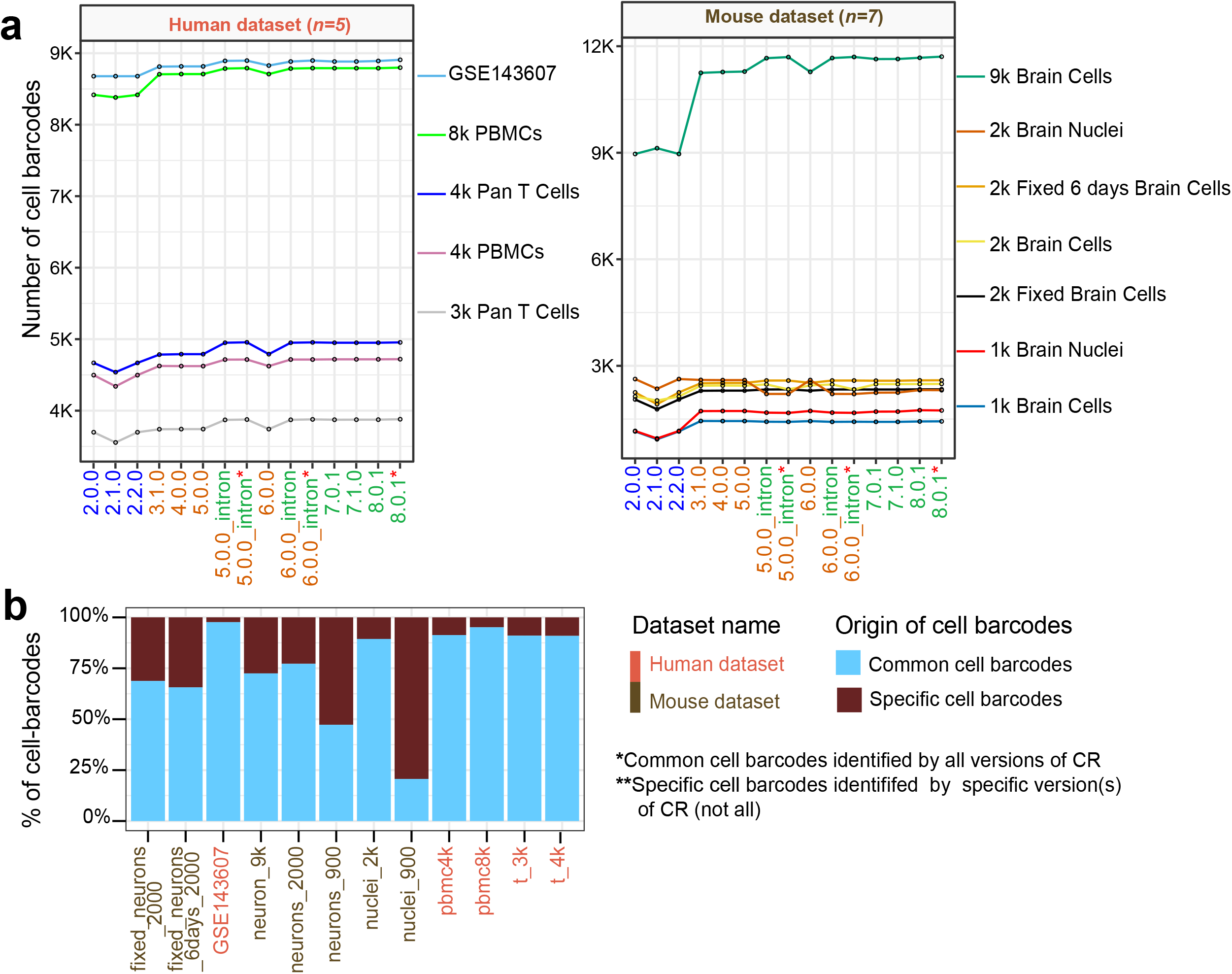
Basic characteristics of cell barcodes generated by different versions of Cell Ranger pipeline. **a**. By basic counting of the resulting cell barcodes by different versions of CR, we found variations in the number of cell barcodes per CR version. Each line represents one dataset. CR version colored by the main feature of the CR version; CR 2.X was an early version of CR before implementing the EmptyDrops algorithm. EmptyDrops was implemented from version 3.x – 7.x. In v7.x CR development team adjusted the parameters of EmptyDrops. The number of cell barcodes changed in version v2.1.0 (reduced) compared to v2.0.0 and v2.2.0. Cell Ranger v3.1.0 and later introduced many cell barcodes, and the number of barcodes kept increasing until the latest version (v8.0.1). **b**. Type and percentage of cell barcodes identified in each dataset. The figure demonstrates the ratio of specific cell barcodes (identified by some version(s) of CR) to the common cell barcodes (identified by all versions of CR) per dataset. The percentage of specific cell barcodes is different between different cell types and sample processing (fixed/unfixed).

As an example, for the dataset of PBMCs from a Healthy Donor (pbmc_8k and pbmc_4k), we retained 12,914 cell barcodes by v2.0.0, 12,721 by v2.1.0 (decreased), and 12,914 by v2.2.0 (increased). The number of cell barcodes increased using v3.1.0 to 13, 330 cell barcodes. A drastic increase in cell barcodes happened when the dataset was processed with v7.0.1 and v7.1.0 (13,508). This resulted in an addition of 787 cell barcodes from v2.1.0. Number of cell barcodes continues to increase when processing the dataset with v8.0.1 and v8.0.1 using the latest transcriptomic reference.

The datasets “nuclei_900”, “neurons_900”, and “fixed_neurons” originating from combined cortex, hippocampus and subventricular zone, retained high numbers of specific cell barcodes when reprocessed with version 3.1.0 and later compared to primary cells (Pan T Cells) and cell lines (MCF10A) (Fig. 1b). This suggested an effect by sample source and preparation in the resulting scRNA-seq dataset processed by different versions of CR. During the analysis of the datasets “nuclei_900” and “neurons_900” we detected some issues (alerts), low fraction reads confidently mapped to transcriptome or low fraction reads in cells. We didn’t apply any solutions to handle this problem.

### Two diverse categories of cell barcodes identified by CR version per dataset

To further characterize the resulting cell barcodes, we compared retained cell barcodes to determine the similarities and differences (in barcode names) between the resulting cell barcodes for each version of CR. We classified two diverse categories of cell barcodes; the first category referred to in this study as ‘**common**’ cell barcodes which were called (or unmasked) by all versions of CR; and the second category, the ‘**specific**’ cell barcodes called (or masked/unmasked) by specific but not all versions of CR. We noticed variation in the ratio of common to specific cell barcodes among datasets (Fig. 2b). This quantitative analysis of cell barcodes indicated that with every release of CR, a specific set of cell barcodes were called, hence the observed increases or decreases seen in the number of cell barcodes (n) per dataset. The choice of CR version therefore will impact the reported number of cell barcodes in a study. The inclusion of a large number of specific cell barcodes took place after v3.0.0, the specific cell barcodes are unmasked at step two of the cell barcodes calling (<u>Calling Cell Barcodes</u>). But CR has no support to annotate those cell barcodes.

After analyzing the number and origin of cell barcodes, we next investigate two questions, first test whether the common cell barcodes generated by different versions of CR have similar molecular profile (UMI, Genes, mitogens, Scrublet scores, and ambient RNA). The second was to find the variations between common and specific cell barcodes in each dataset.

### Significant variations in molecular qualities between cell barcodes called by different versions of CR

We observed that the molecular profile of the common cell barcodes is changed by different versions of CR (Fig. 2a and Supplementary Fig. 2a). The lowest number of detected UMIs and genes were yielded by CR v2.0.0, v2.1.0, v2.2.0, v3.1.0, v4.0.0, v5.0.0 and v6.0.0. These versions only included exonic reads. The number of detected UMIs and genes improved by including the intronic reads and by using the latest reference transcriptome.

Analyzing the molecular quality by Seurat, we identified significant differences in the number of detected UMIs, and genes and percentage of mitGenes between common and specific cell barcodes generated by different versions of CR (Fig. 3a and Supplementary fig. 2a). In the dataset “8k PBMCs from a Healthy Donor” (Fig. 3a), the common cell barcodes showed high values of detected UMIs compared to the specific cell barcodes (median 4,221 vs 962), detected genes (median 1,378 vs. 514), and lower percentage of mitGenes (median 2.6% vs. 6.4%). Likewise, we observed variations in the above qualities among the common cell barcodes in other datasets (upper panel of Fig. 3a and Supplementary fig. 2a). We also quantified the ratio of ambient RNA and observed a lower ratio of ambient RNA contamination among common barcodes compared to specific barcodes (median 0.04 vs. 0.06) (Fig. 3a and Supplementary fig. 2a). Furthermore, we observed that common cell barcodes have higher Scrublet scores compared to the specific cell barcodes (median 0.04 vs. 0.01) (Fig. 3a and Supplementary fig. 2a). In general, the best quality values were obtained by CR 6.0.0 (including intron reads & latest transcriptome reference) and v8.0.1 with latest transcriptome reference as well. The poorest quality was seen in datasets processed with by v3.1.0, v4.0.0, as well as v5.0.0 and v6.0.0 without intronic reads (Fig. 3a and Supplementary fig. 2a). The increased number of UMI counts was explained in 10x Genomics technical note and attributed to the inclusion of intronic reads (see above). However, for CR versions with only exonic read (v3.0.1, v4.0.0, v5.0.0 and v6.0.0) we still observed increases in UMI count. We speculate that the gene model version could have an impact on the number of detected genes.

**Figure 3.**
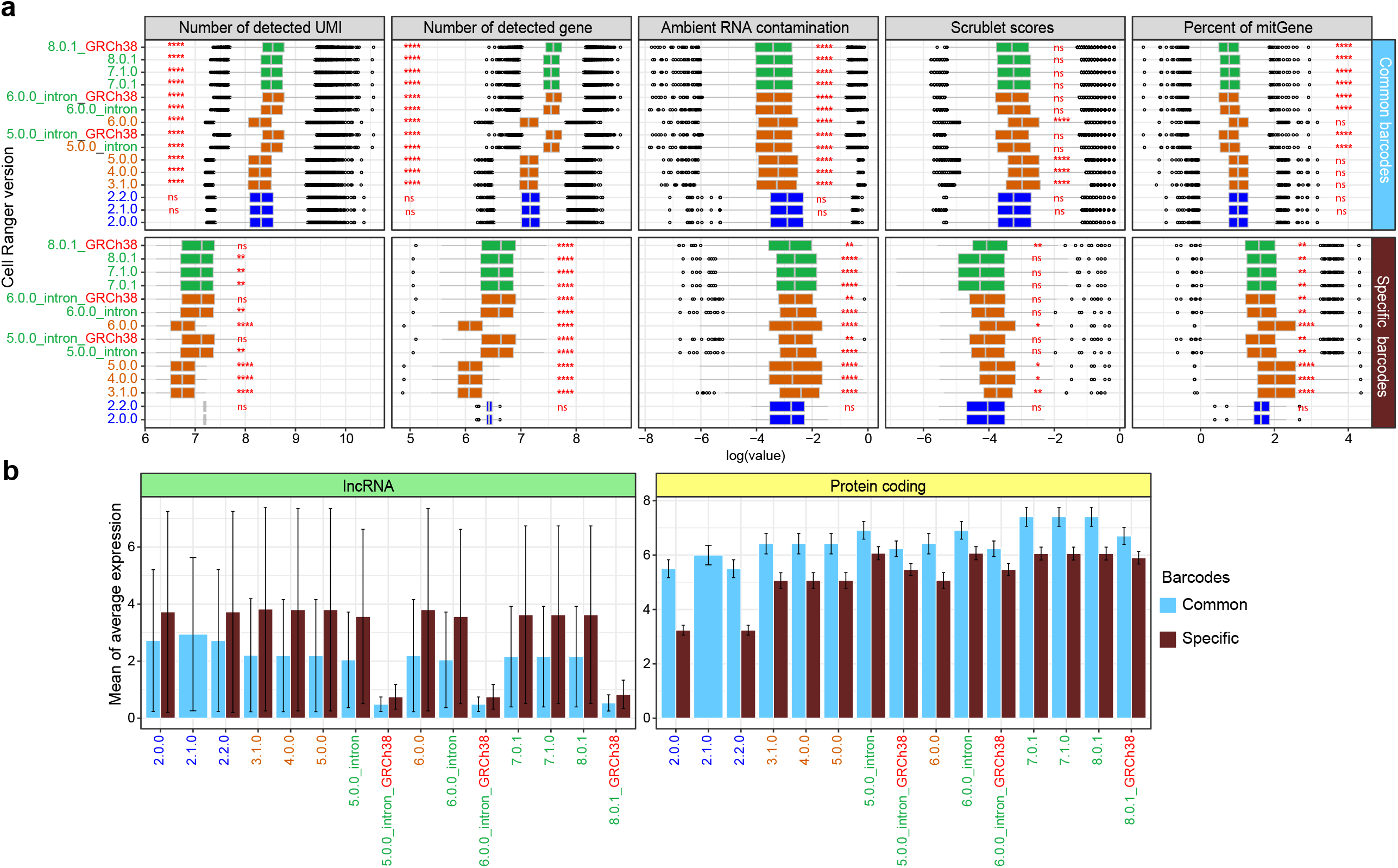
Significant differences in molecular qualities, ambient RNA contamination and Scrublet scores between common and specific cell barcodes. **a**. Significant differences between common and specific barcodes in molecular qualities determined by number of detected UMIs, genes, ratio of ambient RNA contamination, Scrublet scores, and percent of mitGenes. The upper panel shows common barcodes, and the lower panel shows specific barcodes. In the boxplot, the Y-axis represents the CR version, and the X-axis represents the quality value. The mean comparison p-values are added to the boxplot to indicate the statistical significance. The reference version for the t-test was CRv2.0.0. We used asterisks to show the p-values in the plots ns P > 0.05, * P ≤ 0.05, ** P ≤ 0.01, *** P ≤ 0.001, **** P ≤ 0.0001. **b**. Bar plot showing the average gene expression of lncRNA (left) and protein coding genes (right) for human 8k PBMCs from a Healthy Donor. Each bar in the plot represents a Cell Ranger version and the error bar represents the SEM. The bars are colored by the type of cell barcodes.

### Average gene expression of protein-coding and lncRNA impacted by CR version

Detection of protein coding and non-coding (lncRNA) genes is an important quality of scRNA-seq protocols in general. To determine the impact of CR version choice on the average gene expression of protein-coding and lncRNA, we used the GENCODE annotation (gencode.v44.annotation.gtf for human and gencode.vM33.annotation.gtf for mouse) for annotation **(METHODS)**. We used Seurat’s AverageExpression function to obtain the average feature expression by identity class. We found significant variation between common and specific barcodes, in terms of average gene expression of protein-coding and lncRNA, for each dataset when generated by different versions of CR. In contrast to specific cell barcodes, common cell barcodes yielded a higher average gene expression of protein-coding genes (right panel of Fig. 3b and Supplementary fig. 3b). Also, CR v7.0.1 and v7.1.0 gave a higher average gene expression of protein-coding genes for common cell barcodes compared to the rest of the CR versions analyzed. We observed the opposite for the average gene expression of lncRNA, the average gene expression was higher for specific cell barcodes than for common barcodes in all datasets, but not in the “2k Brain Nuclei from an Adult Mouse (>8 weeks)” dataset (Supplementary fig. 3b). CR v5.0.0, v6.0.0, v8.0.1 (including intron reads & latest transcriptome reference) gave lower levels of average gene expression of lncRNA compared to the other CR versions. In the technical note (<u>Interpreting Intronic and Antisense Reads in 10x Genomics Single Cell Gene Expression Data</u>), 10x Genomics has recommended to include the intronic read to increase the sensitivity, i.e. mean genes-per-cell detected. In contrast to the technical note (<u>Interpreting Single Cell Gene Expression Data With and Without Intronic Reads</u>) our observation shows negative impacts on sensitivity when including intronic reads with the recommended transcriptome reference (Fig. 3b and Supplementary fig. 2b) especially on the average gene expression of lncRNA.

The variation in the average gene expression of protein-coding and lncRNA genes will impact differential gene expression analyses, gene set enrichment analyses and marker gene stability, and thus also the biological interpretation of the data.

### Specific cell barcodes exhibits skewed gene body coverage

We investigated the patterns of the gene body coverage for cell barcodes called by different versions of CR. Gene body coverage of the common and specific cell barcodes were computed and visualized by SkewC^9,18^ (Fig. 4 and Supplementary fig. 3). We found that common cell barcodes had a typical gene body coverage expected for 3’ scRNA-seq (peak of the coverage at the 3’ end of the gene region) while the specific cell barcodes had a more skewed gene body coverage (peak at the 5’ end or at the middle of the gene region). We observed variations in the size of the skewed cell populations among datasets. Nuclei from cells from “combined cortex”, “hippocampus”, and “subventricular zone of an E18 mouse”, had a high percentage of skewed cells compared to the other datasets. This could be due to technical failure or biological condition of the sample cells (e.g., apoptosis, among other explanations).

**Figure 4.**
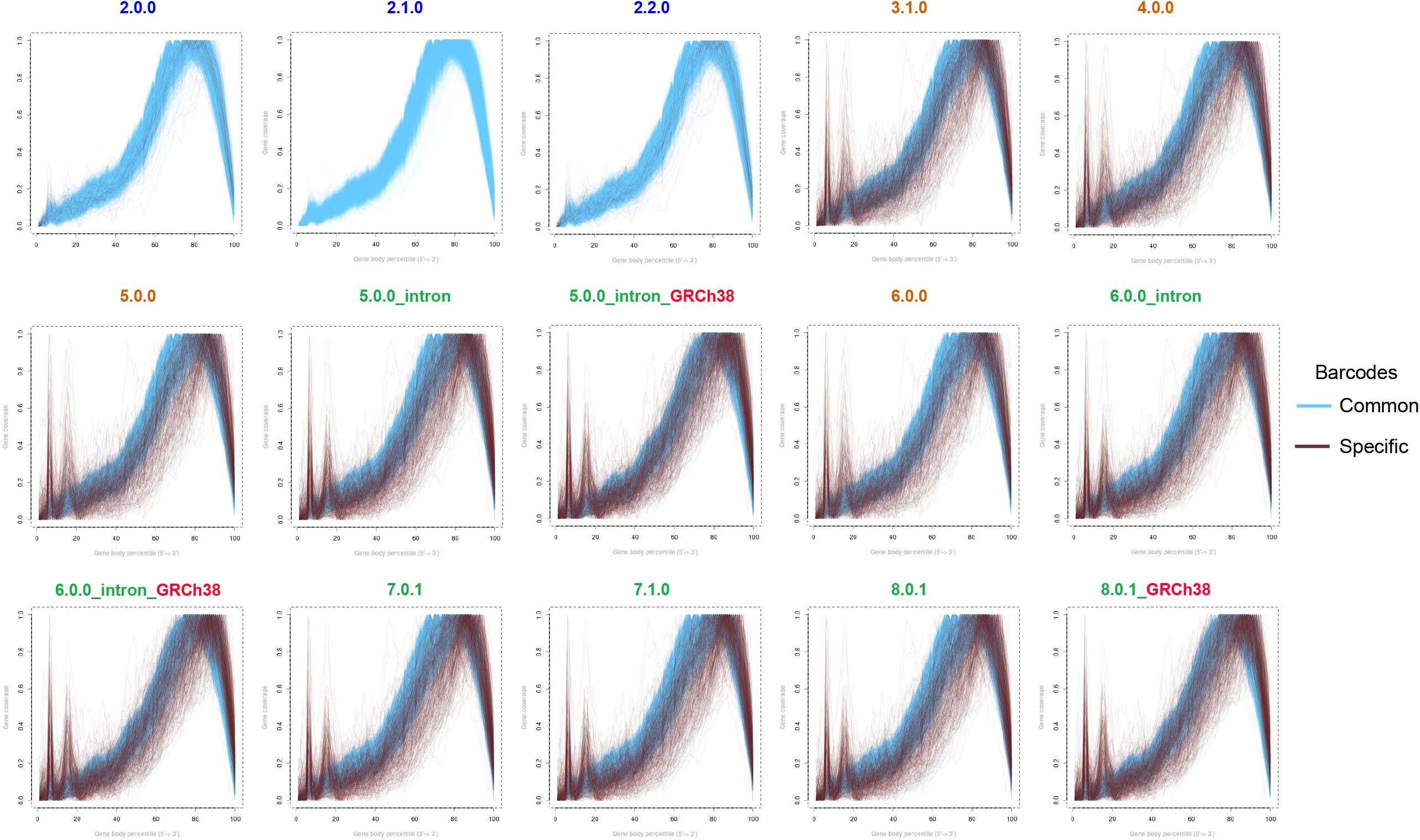
Gene body coverage skewness of common and specific cell barcodes. Gene body coverage of the dataset 8k PBMCs from a Healthy Donor as produced by all versions of CR. The gene body coverage was computed and visualized by SkewC. Each line represents a cell barcode. X-axis shows the gene body coverage percentile 5`-end to 3`-end and the Y-axis is the gene body coverage value. Gene body coverage plots are colored by the type of cell barcodes. Note that the specific cell barcodes have peaks at the 5` end of the gene region and are therefore skewed.

### Specific cell barcodes formed a distinct population of cells at the edge of the embedded clusters

The significant variation in the molecular qualities of the datasets generated by different versions of CR and the variation between the common and specific cell barcodes led us to investigate the expression patterns and biological characteristics of common and specific cell barcodes. We investigated the distribution/localization of common and specific cell barcodes in dimensionality reduction plots. We used Uniform Manifold Approximation and Projection (UMAP)^19^ using a common ‘random state’ setting to avoid changes in the plot due to UMAP’s inherent stochasticity. All embeddings were made following normalization of expression with the LogNormalize method in Seurat (Methods). We observed that specific cell barcodes formed a distinct population of cells and clustered together at the edge of the embedded clusters (Fig. 5 and Supplementary fig. 4). This suggested that the specific cell barcodes have some inherently similar characteristics with each other, and revealed different biological characteristics compared to the common cell barcodes.

**Figure 5.**
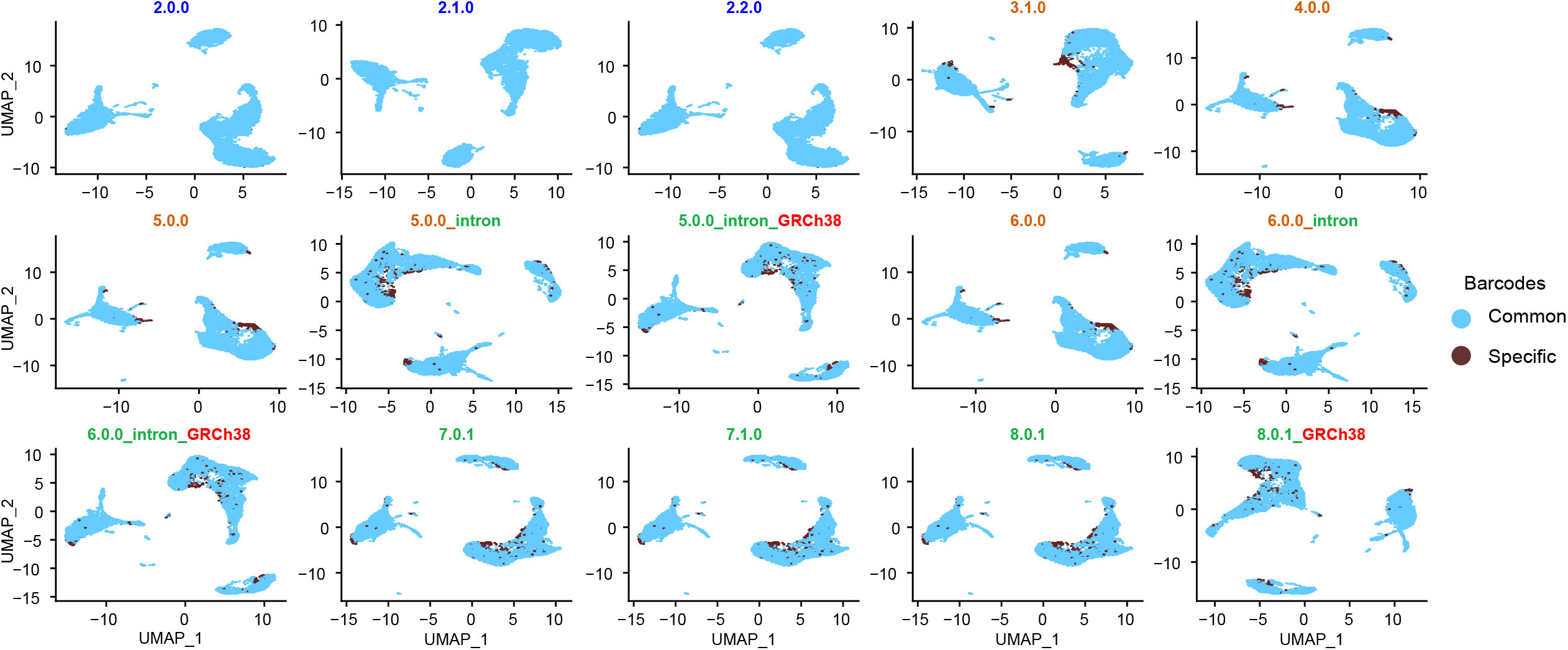
Characteristics of common and specific barcodes. UMAP embedding of PBMC_8k cells generated by different versions of Cell Ranger colored by the source of the cell barcodes. Cell barcodes in light blue represent the cell barcodes called by all versions of Cell Ranger (common barcodes). Cell barcodes colored by brown are the cell barcodes called by the specific Cell Ranger version (specific barcodes).

### Number of clusters and cluster sizes changed with each CR version

We moved on to investigate the impact of CR versions on downstream scRNA-seq analyses. We considered two types of downstream analyses applied in almost all studies: unsupervised clustering and differential gene expression (DGE) analysis. For each dataset generated by each CR version, we identified clusters of cell barcodes using Seurat’s FindClusters() function that supports shared nearest neighbor modularity optimization based graph creation and the Leiden clustering algorithm. The analysis enabled us to determine the number of clusters and cluster sizes produced from data analyzed with each CR version. Then, we annotated cell barcodes in each cluster as either common or specific cell barcodes (Fig. 6a and Supplementary fig. 5a). We demonstrated that the number of clusters and cluster sizes obtained by unsupervised clustering changed with each CR version. The variation in the inferred number of clusters was evident in all datasets. The number of clusters changed from 11 clusters in v2.0.0 to 15 clusters in v7.0.1, v7.1.0 and v8.0.1 (Fig. 5a). Moreover, we observed changes in cluster size in all datasets. For example, cluster 0 size changed drastically from 1,750 cell barcodes in v2.0.0 to 2,000 cell barcodes in v3.1.0, v4.0.0, v5.0.0, and v6.0.0 (Fig. 6a). This then dropped to 1,500 cell barcodes in v7.0.1 and v7.1.0. These results clearly indicated that different versions of CR can result in deviations in both the number of clusters and the barcodes attributed to each cluster.

**Figure 6.**
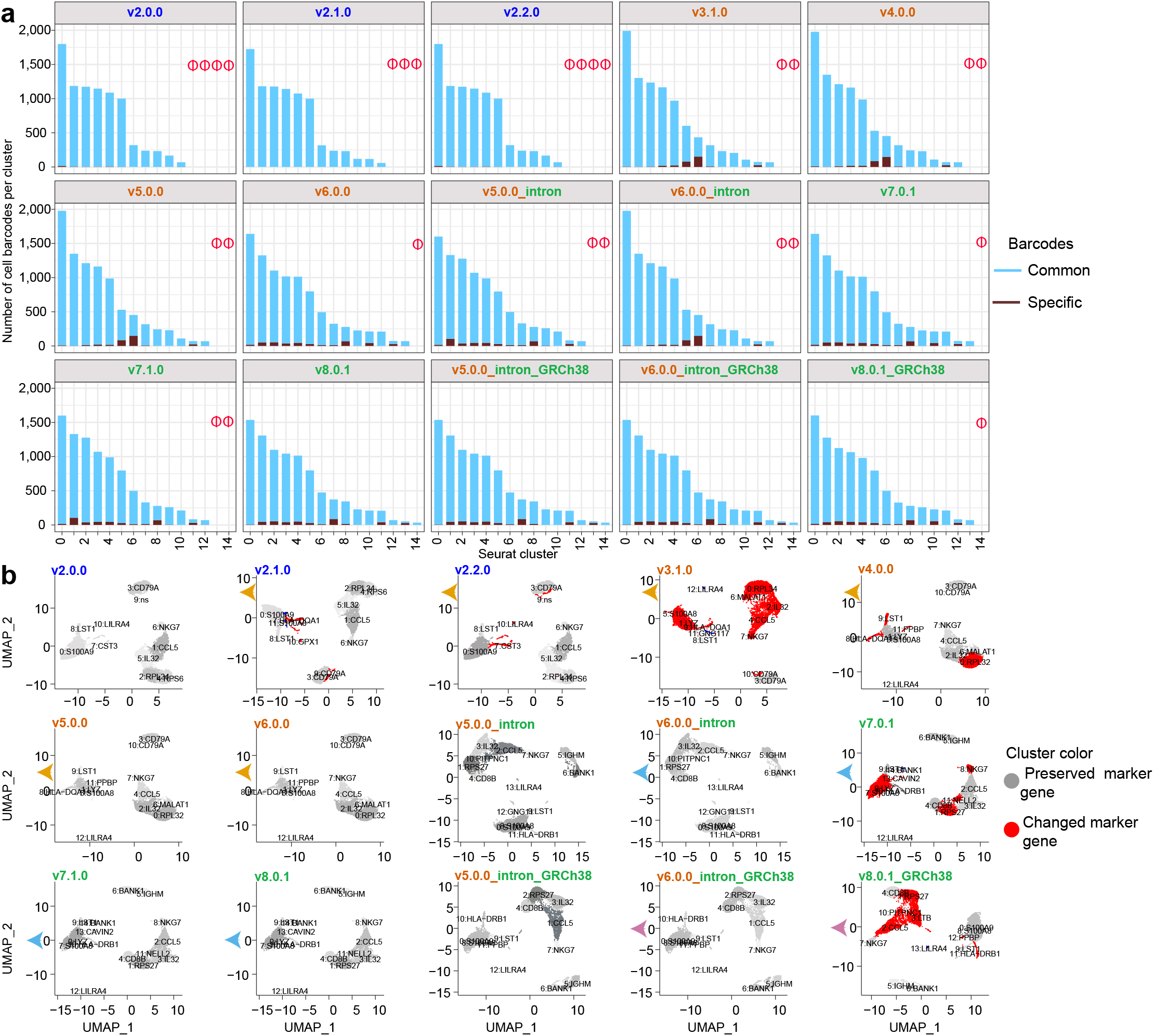
Impact of cell calling algorithm on number of clusters, cluster size, and cluster marker genes. **a**. Number of clusters and cluster size changed per Cell Ranger version. Each bar plot represents one dataset generated by a version of CR. One bar per Seurat cluster, and each bar shows the number of identified cell barcodes in that cluster. The bars are colored by the type of cell barcode. The phi (1) symbol indicates that no cluster was identified. **b**. UMAP plots of the “common” cell barcodes, each represents a UMAP embedding produced for each Cell Ranger version. Each cluster in the UMAP is labeled with the cluster number and top marker gene. Unsupervised clusters in each UMAP are colored by the type of cluster markers. Red colored clusters indicate that the marker genes are changed from a previous CR version while the gray colored clusters indicate that the marker gene was conserved (not changed) from the previous version of CR. The arrows at the y-axis of each UMAP, indicate the comparison direction (e.g. UMAP produced by CR v2.1.0 compared against the UMAP produced by v2.0.0, another example, UMAP generated by v8.0.1_GRCh38 was compared against the UMAP generated by v6.0.0_intron_GRCh38). Drastic changes in the cluster marker genes observed in UMAP of v3.1.0, v7.0.1 and v8.0.1_GRCh38. For compatibility, we didn’t highlight the comparison between v5.0.0 vs v5.0.0_intron and v6.0.0 vs v6.0.0_intron etc. but differences in the marker genes were seen.

### Cluster marker genes are changed by change of CR version

Next, we performed DGE to identify cluster markers by fetching conserved top marker gene(s) for each cluster in each dataset. To prevent the contribution of the specific cell barcode data to the DGE analysis we only used the common cell barcodes. We observed that cluster marker genes changed within the same dataset analyzed by different versions of CR (Fig. 6b and Supplementary fig. 5b). We observed that cluster marker genes were either preserved between versions or were changed either by introducing new markers or new clusters. The lncRNA MALAT1 (Metastasis Associated Lung Adenocarcinoma Transcript 1) was identified as a cluster marker gene for cluster number 6 in a dataset analyzed by v3.1.0, v4.0.0, v5.0.0 and v6.0.0 (Fig. 6b). MALAT1 is conserved in mammals and is associated with several diseases^20^. In the MCF10A dataset (Supplementary fig. 5b), Nuclear Enriched Abundant Transcript 1 (NEAT1), a nuclear lncRNA, was the top marker gene for cluster 2 when processed by all versions of CR save for v7.0.1 and v7.1.0; in these versions two more clusters were added and MALAT1 replaced NEAT1. This same observation about the change in lncRNAs as cluster markers could be observed for the rest of the datasets (Supplementary fig. 5b). This analysis shows that cluster marker genes are impacted by CR version regardless of the inclusion / exclusion of the intron read.

## Discussion

Our study demonstrates that the number and molecular quality of cell barcodes is strongly influenced by CR version choice and setup (inclusion of intronic read, version of the gene model). The latest version of CR introduced a large set of specific cell barcodes when compared to the oldest CR version analyzed here for a variety of mammalian datasets. We categorized the resulting cell barcodes to two categories, common and specific. Overall, the specific cell barcodes have poorer molecular qualities and skewed gene body coverage when compared to common cell barcodes. The ratio of the specific cell barcodes varies per sample type and processing method.

According to the release note of CR 3.X, the development team has changed the cell-calling method to include EmptyDrops^5^, but it is not obvious whether any other changes in the cell-calling algorithm were introduced in version 3.X and later. Further investigation (Code walkthroughs) might help to understand the interaction and implementation of EmptyDrops in CR and the cell calling stage FILTER_BARCODES.

The molecular qualities of the common cell barcodes improved by the introduction of CR v7.0.1 due to the adjustment of EmptyDrops parameters by the CR development team. EmptyDrops approach was based on detecting statistically significant deviations from the expression profile of the ambient solution. Our analysis demonstrated that post-EmptyDrops CR versions (3.X – 8.X) unmasked cell barcodes that were masked in pre-EmptyDrops CR versions (2.X).

UMAP is a stochastic method and running the same embedding multiple times yields slightly different results. The population of specific cell barcodes at the edge of UMAP cluster(s) impacted by some versions of CR could form artificial clusters and lead to misinterpretation of the clusters when the cluster markers are changed. The diversity of cell/sample types used in this study enabled us to investigate the impact of sample type origin and biological condition of the cell on the resulting cell barcodes when the dataset was processed by different versions of CR. The analysis of the common cell barcodes indicated that different CR versions altered molecular characteristics of individual cell barcodes. The change in quantification of the protein-coding genes and lncRNA between CR versions might have a significant impact on the biological interpretation of the dataset and influence studies investigating lncRNAs.

To address the quality issues due to the cell barcode calling algorithm, we anticipate that the CR development team will provide an option for the user to separate common cell barcodes from specific cell barcodes. Providing this feature, the user will be able to work with common cell barcodes and QC them further.

Our study demonstrates the limitation of the 10x Genomics evaluation study (<u>Interpreting Single Cell Gene Expression Data With and Without Intronic Reads</u>) on the impact of the inclusion of the intron reads on Chromium gene expression data analysis. The study was limited to one version of CR v6.1.2 and the study conclusion was questionable.

For the re-use of published 10x Genomics scRNA-seq dataset (e.g., HCA datasets, Tabula Sapiens^21^, etc.) generated by any of the post-EmptyDrops CR versions we recommend careful quality checking of the data before re-using the dataset for downstream analyses.

In conclusion, our comprehensive evaluation and analysis of the impact of CR version, indicates that, the choice of CR version affects scores for quality, gene expression, clustering, and top cluster marker genes of the dataset.

## Methods

### Study workflow

To investigate the impact of the cell-calling algorithm on the quality of scRNA-seq data, we designed the workflow in Fig. 1a. The workflow covers single cell (SC) gene expression chromium dataset acquisition, cell barcodes calling, and CR output (Fig. 1a). In addition to the analysis workflow used to determine different types of quality metrics and investigate CR version impact on downstream analysis (bottom panel Fig. 1a).

### SC gene expression chromium dataset acquisition

We used 12 datasets in this study. The raw sequence read files (FASTQ) were downloaded either from the 10x Genomics data portal (<u>10xgenomics.com/datasets)(n=11</u>) or NCBI GEO (n=1). The datasets studied cover human (n=5) and mouse (n=7) primary cells and cell lines (Supplementary Table 1, and the downloaded datafiles in **Data availability**). The associated metadata for each dataset is available at SCPortalen2^22^. All datasets were generated by Single Cell Chromium 3’ GEX v2 which supports all versions of CR.

### Cell barcode calling and basic analysis

The downloaded FASTQ files were processed by 10 versions of the CR pipeline (v2.0.0, v2.1.0, v2.2.0, v3.1.0, v4.0.0, v5.0.0, v6.0.0, v7.0.1, v7.1.0 and v8.0.1) using the *cellranger count* command. The default parameters were used at first. For v5.0.0 and v6.0.0 we reprocessed the data by following the 10x Genomics (<u>Recommendation on Including Introns for Gene Expression Analysis</u>) (--include-introns) flag. As for transcriptome reference, in CR v2.X we used refdata-cellranger-GRCh38-1.2.0 for human and refdata-cellranger-mm10-1.2.0 for mouse. For the rest of the versions of CR we used refdata-cellranger-GRCh38-3.0.0 for human dataset and refdata-cellranger-mm10-3.0.0 for mouse dataset.

Furthermore we ran CR v5.0.0 (including introns), v6.0.0 (including introns) and 8.0.1 using refdata-gex-GRCh38-2024-A for human dataset and refdata-gex-GRCm39-2024-A for mouse dataset. The run command is stored in a text file (_cmdline) in the output folder of CR.

CR generates several files, the output from each run provided in the Data Availability section. In our analysis we used the files in the folder filtered_feature_bc_matrix which contains TAB delimited text files with the barcodes list, another file of the same type with the list of genes, and the expression matrix in Matrix Market format (.mtx). In addition, we used the ‘possorted_genome_bam’ BAM file and the metrics summary text file.

To count the resulting cell barcodes from each CR version, we used the file text barcodes list which is used to plot Fig. 1b. The same file was used to investigate the similarities and differences between cell -barcodes which lead to categorization of cell barcodes as common or specific cell barcodes (Fig. 1c.).

### Determining molecular quality with Seurat

Seurat QC^8^ metrics of scRNA-seq cover three molecular characteristics of cell barcodes: detected number of UMI (nCount_RNA), genes (nFeature_RNA), and the percentage of mitochondrial genes (percent.mt). To perform the Seurat QC, Seurat R objects were created for each dataset. We used the R implementation of Seurat version 4.3.0. The R Seurat function *CreateSeuratObject()* enables the creation of a Seurat object for each dataset. The output of *CreateSeuratObject()* was an R object with basic metadata information for each cell barcode. The basic metadata includes two QC variables:

nFeature_RNA and nCount_RNA. To obtain the percent.mt for each cell barcode, we used the function *PercentageFeatureSet()*. This function takes as input two parameters: the first was the R object created itself and second was the pattern; we used the pattern = “^MT-” to get the number of mitochondrial genes.

We added two metadata columns named CR and barcode using *AddMetaData()*. The CR column distinguishes the CR version of the Seurat R object, and the barcode column indicates the category of the cell barcode as described above. Quality values from Seurat were used to plot the box plot in Fig. 2a and Supplementary Fig. 2a.

### Compute doublet scores with Scrublet

Scrublet ^17^ detect doublets in scRNA-seq data (transcription-based). We used the wrapper script provided by Scrublet to get Scrublet scores. We used the following command to run Scrublet in a Singularity container:

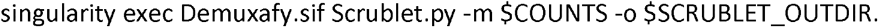

Where the $COUNTS was the filtered_feature_bc_matrix for each dataset. For each dataset Scrublet gives two text files and two image files. The text file Scrublet_results.tsv contains three columns, the cell barcode, the Scrublet_droplet type and Scrublet scores. The Scrublet_summary.tsv gives an overall summary of the total number of singlets and doublets. Finally, we used the function *AddMetaData()* to add Scrublet scores to the Seurat R object of each dataset.

### Computing the ratio of Ambient RNA contamination

The R Bioconductor package decontX was used to compute the ratio of ambient RNA contamination. DecontX accepts the output from CR by using the function *importCellRanger()*. The result of running the function *decontX()* on the exported Cell Ranger object will result in a text file with the ratio as a floating point number between 0-1 of ambient RNA per cell barcodes. The ratio of ambient RNA per cell was added as a metadata column as described above.

### Computing gene body coverage with SkewC

SkewC^9,18^ takes the possorted_genome_bam BAM file and the barcodes text file as inputs. SkewC splits the pooled BAM files into individual BAM files (one BAM file per cell barcode), then computes the gene body coverage. For each dataset generated by CR versions, we computed and visualized the gene body coverage from the 5`-end to the 3`-end of the gene region. The gene body coverage was used to plot Fig. 3 and Supplementary Fig. 3.

### Dimensionality reduction and UMAP plotting

The Seurat R objects were used to create the UMAP unsupervised clusters for each dataset. Several functions are used prior to plotting the UMAP (Seurat tutorial) (Code Availability). In short, each dataset was normalized using the *LogNormalize* method with a scale factor of 10,000. To identify the top 2,000 variable genes, *FindVariableFeatures* method was used using the ‘vst’ method. The top variable genes were used by the *RunPCA* function to perform PCA dimensionality reduction. Top ten Principle components were used by the *FindNeighbors()* function to compute and construct a Shared Nearest Neighbor Graph. The function *RunUMAP()* was used to embed the UMAP dimensionality reduction for each dataset; a random seed was set to remove the impact of stochasticity in our results. Finally, the *DimPlot()* function is used to plot the UMAP (See code availability).

### Cluster marker identification

To identify the differentially expressed markers for each cluster, we used the annotated Seurat R object created previously and called the Seurat *FindAllMarkers()* function. We subset the R object and limited it to the common cell-barcodes.

## Data availability

We provide the raw and processed data for this study’s datasets via the following website:

https://single-cell.riken.jp/suppl/skewc/Cell_Ranger_Study_Data_Files/. We created a folder for each dataset that contains the following subfolders and files:

CR2.0.0_dataset_name/ [CR output files].

CR7.1.0_dataset_name/ Seurat_Objects_RDS/

dataset_name_2.0.0.rds

…

dataset_name_7.1.0.rds SkewC_Coverage/

dataset_name_2.0.0/ [For each cell barcode, the folder contains .R and .text file with the full gene body coverage]

possorted_genome_bam.AAACCTGGTCTCGTTC-1.geneBodyCoverage.r possorted_genome_bam.AAACCTGGTCTCGTTC-1.geneBodyCoverage.txt

…

dataset_name_7.1.0/

possorted_genome_bam.AAACGGGTCCGCGGTA-1.geneBodyCoverage.r possorted_genome_bam.AAACGGGTCCGCGGTA-1.geneBodyCoverage.txt

fastq/ [Source Fastq files from 10x Genomics support or NCBI GEO]

XXXX. fastq.gz

## Code availability

- R Code to Create Seurat object and plot UMAP
https://single-cell.riken.jp/suppl/skewc/Cell_Ranger_Study_Data_Files/R_Code_for_Creating_Seurat_Object_plotting_UMAP.R
- Code to run SkewC

Latest version of SkewC is available in GitHub https://github.com/LSBDT/SkewC

## Acknowledgement

This work was supported by research grants for the RIKEN Center for Life Science Technologies, RIKEN Center for Integrative Medical Sciences, NBDC grant Number JPMJND2202 from Japan Science and Technology Agency (JST), and RIKEN Open Life Science Platform project from MEXT, Japan. JK was supported by Swedish Research Council, Swedish Brain Foundation, Jane and Aatos Erkko Foundation (Finland), and Sigrid Jusélius Foundation (Finland). This work was initiated when JK was a Japan Society for the Promotion of Science Fellow (Japan) at RIKEN Center for Integrative Medical Sciences.

## Supplementary Table legends

**Supplementary Table 1: Single-cell gene expression chromium dataset: List of the study datasets and the change in the number of cell barcodes called by different versions of Cell Ranger**. The table shows the list of the datasets used in the study, type of cells, species (human and mouse) and the cell barcode count for each of the Cell Ranger versions.

## Supplementary figure legends

**Supplementary figure 2a Molecular quality, ratio of ambient RNA contamination, and Scrublet scores Supplementary figure 2b Average expression of protein-coding and lncRNA genes**

**Supplementary figure 3. Gene body coverage of single cell 10x 3` datasets (Human and Mouse)**.

**Supplementary figure 4 UMAP plots**.

**Supplementary Figure 5a. Number of clusters and cluster size**

**Supplementary Figure 5a. Analysis of variable genes and feature expression per cluster**.

## Footnotes

1 https://www.10xgenomics.com/jp/support/software/cell-ranger/latest/resources/supported-libraries

## References

1 Klein, A. M. et al. Droplet barcoding for single-cell transcriptomics applied to embryonic stem cells. Cell 161, 1187–1201 (2015). 10.1016/j.cell.2015.04.044

2 Macosko, E. Z. et al. Highly Parallel Genome-wide Expression Profiling of Individual Cells Using Nanoliter Droplets. Cell 161, 1202–1214 (2015). 10.1016/j.cell.2015.05.002

3 Vallot, C. RNA barcoding: the catalyst for the single-cell revolution. Nat Rev Genet 24, 491 (2023). 10.1038/s41576-023-00624-7

4 Muskovic, W. & Powell, J. E. DropletQC: improved identification of empty droplets and damaged cells in single-cell RNA-seq data. Genome Biol 22, 329 (2021). 10.1186/s13059-021-02547-0

5 Lun, A. T. L. et al. EmptyDrops: distinguishing cells from empty droplets in droplet-based single-cell RNA sequencing data. Genome Biol 20, 63 (2019). 10.1186/s13059-019-1662-y

6 Young, M. D. & Behjati, S. SoupX removes ambient RNA contamination from droplet-based single-cell RNA sequencing data. Gigascience 9 (2020). 10.1093/gigascience/giaa151

7 Yang, S. et al. Decontamination of ambient RNA in single-cell RNA-seq with DecontX. Genome Biol 21, 57 (2020). 10.1186/s13059-020-1950-6

8 Butler, A., Hoffman, P., Smibert, P., Papalexi, E. & Satija, R. Integrating single-cell transcriptomic data across different conditions, technologies, and species. Nat Biotechnol 36, 411–420 (2018). 10.1038/nbt.4096

9 Abugessaisa, I. et al. SkewC: Identifying cells with skewed gene body coverage in single-cell RNA sequencing data. iScience 25, 103777 (2022). 10.1016/j.isci.2022.103777

10 Kedlian, V. R. et al. Human skeletal muscle aging atlas. Nat Aging (2024). 10.1038/s43587-024-00613-3

11 Reed, A. D. et al. A single-cell atlas enables mapping of homeostatic cellular shifts in the adult human breast. Nat Genet 56, 652–662 (2024). 10.1038/s41588-024-01688-9

12 Siletti, K. et al. Transcriptomic diversity of cell types across the adult human brain. Science 382, eadd7046 (2023). 10.1126/science.add7046

13 Lindeboom, R. G. H. et al. Human SARS-CoV-2 challenge uncovers local and systemic response dynamics. Nature 631, 189–198 (2024). 10.1038/s41586-024-07575-x

14 You, Y. et al. Benchmarking UMI-based single-cell RNA-seq preprocessing workflows. Genome Biol 22, 339 (2021). 10.1186/s13059-021-02552-3

15 Rich, J. M. et al. The impact of package selection and versioning on single-cell RNA-seq analysis. bioRxiv (2024). 10.1101/2024.04.04.588111

16 Wolf, F. A., Angerer, P. & Theis, F. J. SCANPY: large-scale single-cell gene expression data analysis. Genome Biol 19, 15 (2018). 10.1186/s13059-017-1382-0

17 Wolock, S. L., Lopez, R. & Klein, A. M. Scrublet: Computational Identification of Cell Doublets in Single-Cell Transcriptomic Data. Cell Syst 8, 281–291.e289 (2019). 10.1016/j.cels.2018.11.005

18 Abugessaisa, I., Hasegawa, A., Katayama, S., Kere, J. & Kasukawa, T. Computational approach to evaluate scRNA-seq data quality and gene body coverage with SkewC. STAR Protoc 4, 102038 (2023). 10.1016/j.xpro.2022.102038

19 Leland, M., John, H., Nathaniel, S. & Lukas, G. Vol. 3 (Journal of Open Source Software, 2018).

20 Amodio, N. et al. MALAT1: a druggable long non-coding RNA for targeted anti-cancer approaches. J Hematol Oncol 11, 63 (2018). 10.1186/s13045-018-0606-4

21 Jones, R. C. et al. The Tabula Sapiens: A multiple-organ, single-cell transcriptomic atlas of humans. Science 376, eabl4896 (2022). 10.1126/science.abl4896

22 Abugessaisa, I. et al. SCPortalen: human and mouse single-cell centric database. Nucleic Acids Res 46, D781–D787 (2018). 10.1093/nar/gkx949

