## Supplementary figures and images for "Impacts of Cell Ranger versions on Chromium gene expression data"

### Supplementary_Figure_2A_fixed_neurons_6days_2000.pdf

fixed_neurons_6days_2000

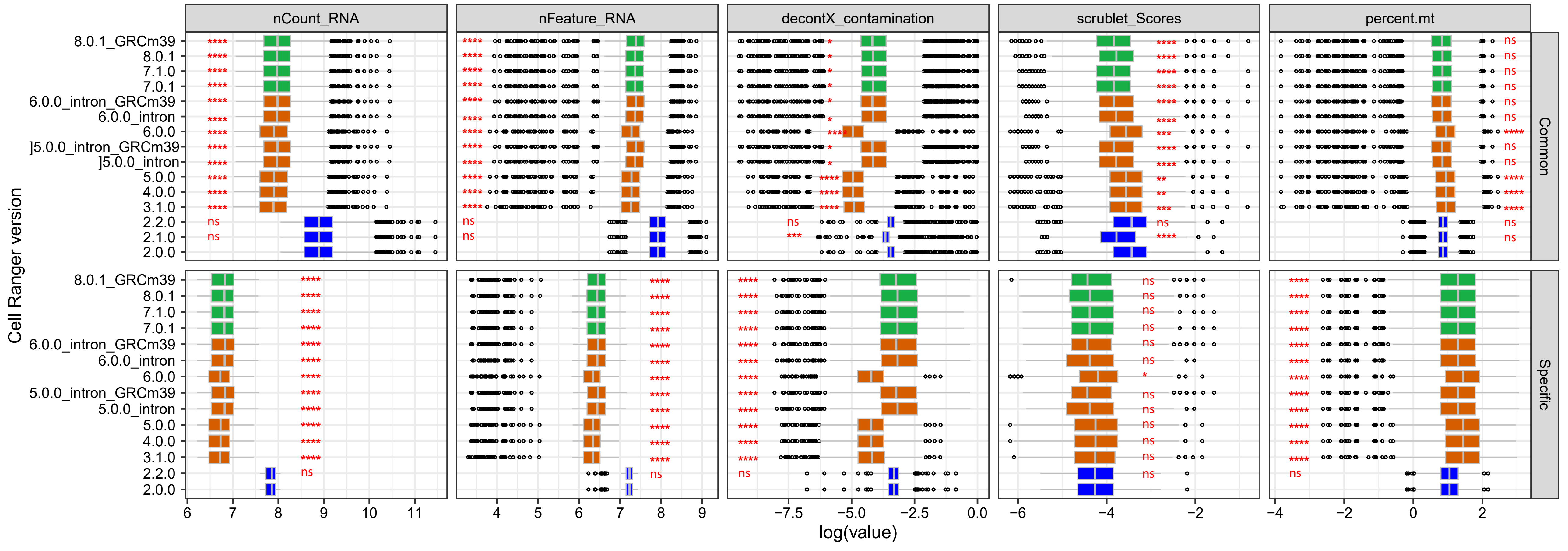

### Supplementary_Figure_2A_fixed_neurons_2000.pdf

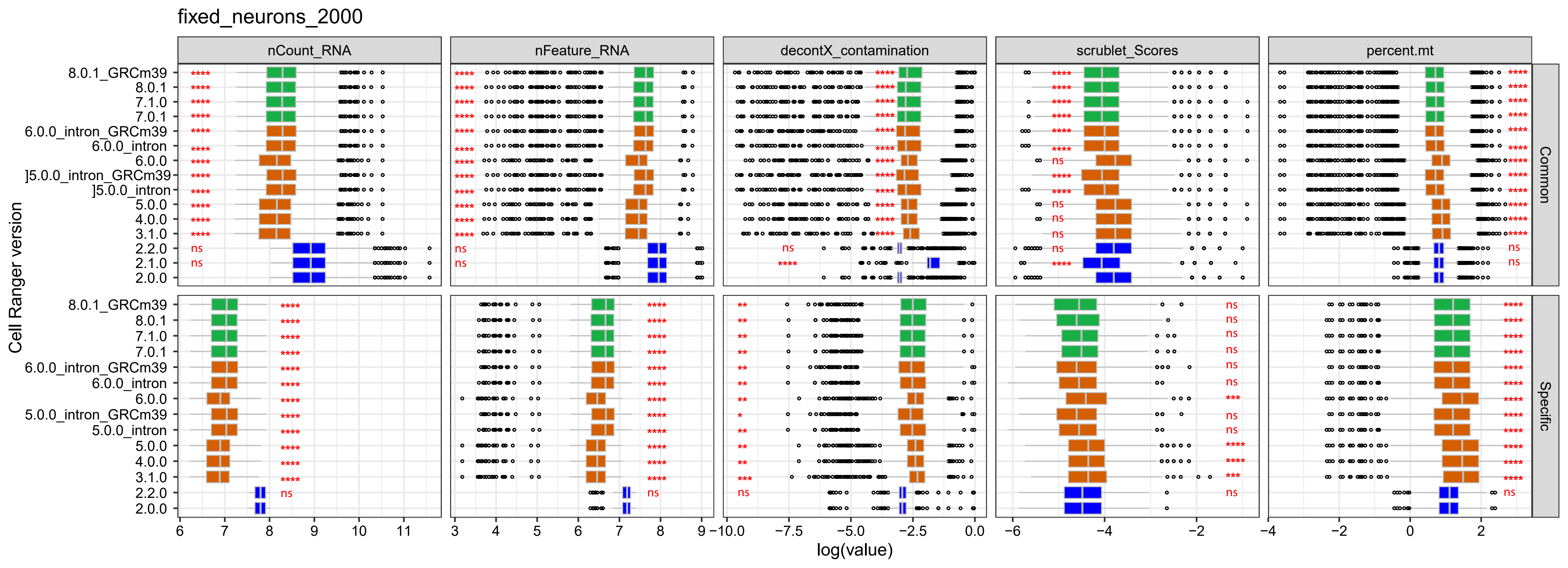

### Supplementary_Figure_2A_GSE143607.pdf

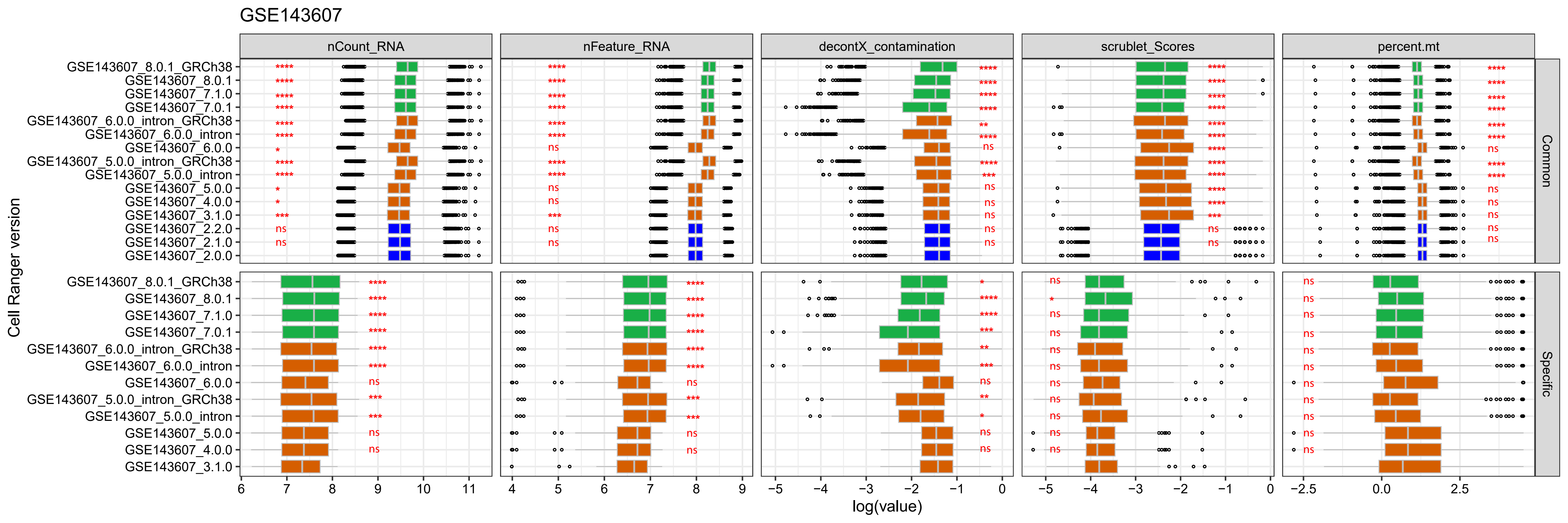

### Supplementary_Figure_2A_neuron_9k.pdf

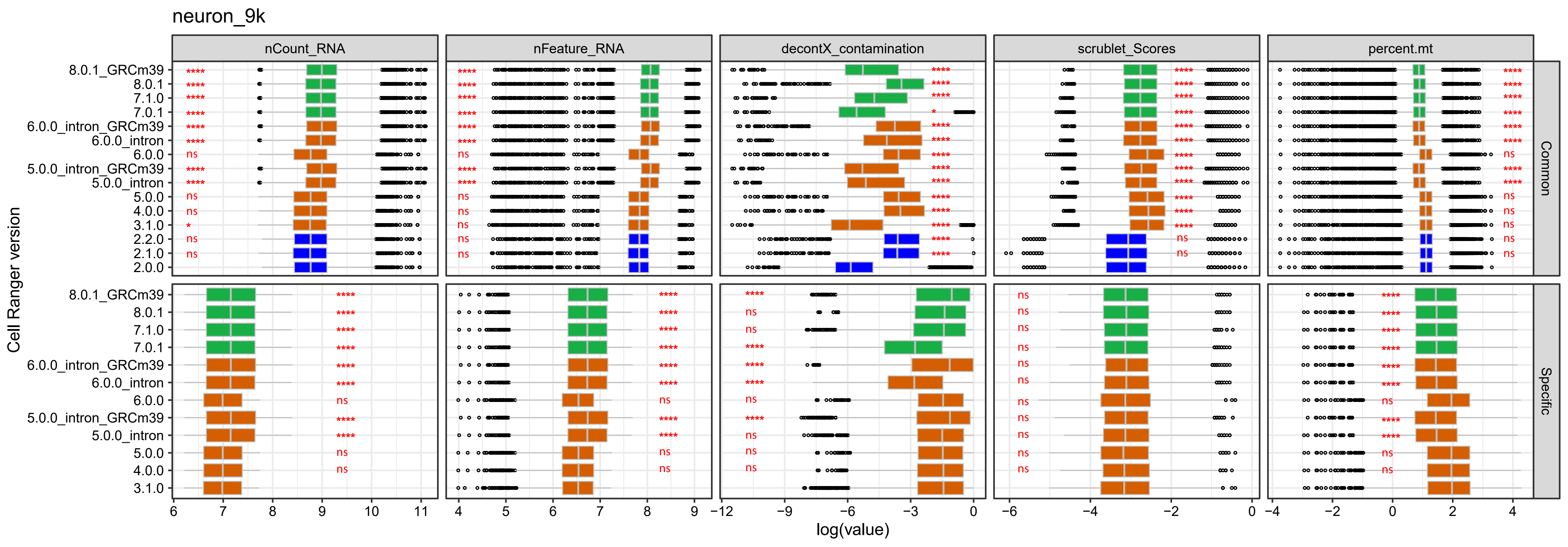

### Supplementary_Figure_2A_neurons_900.pdf

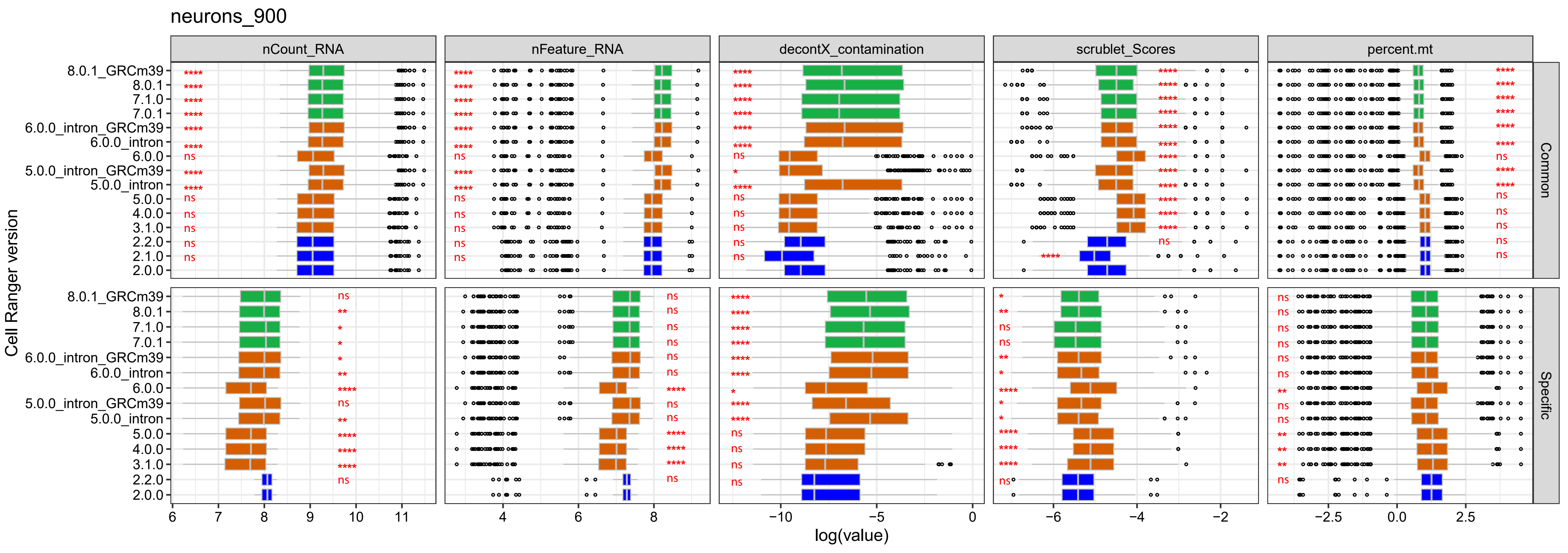

### Supplementary_Figure_2A_neurons_2000.pdf

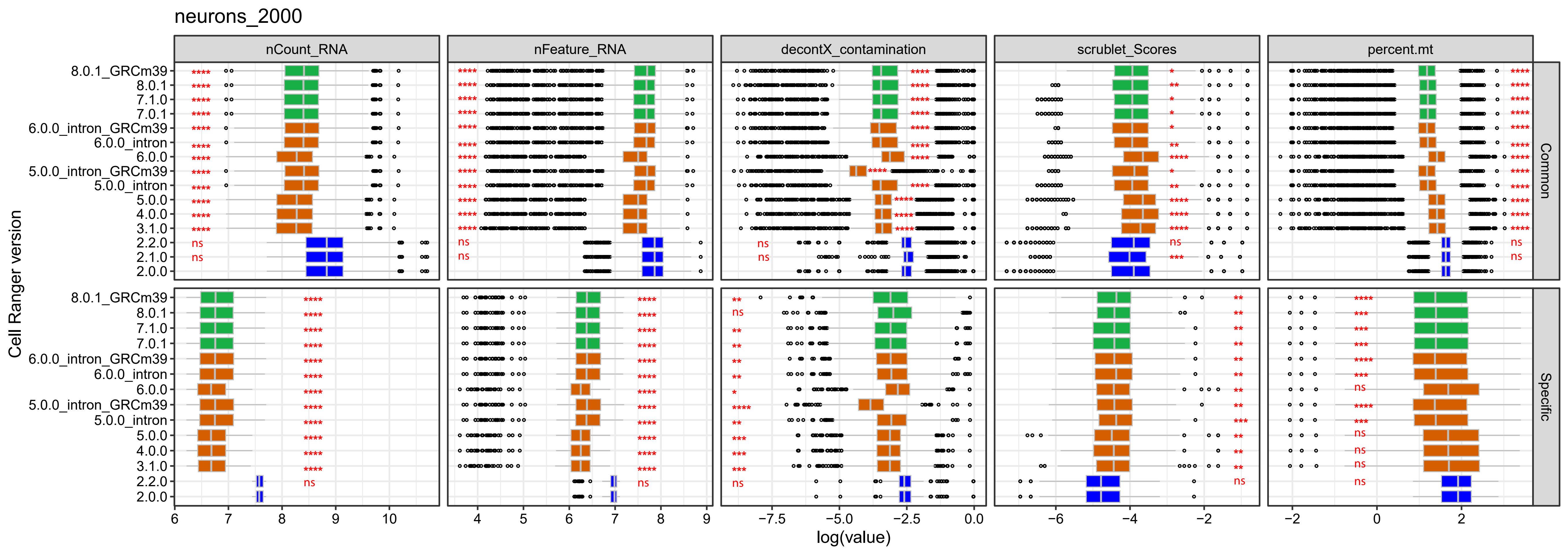

### Supplementary_Figure_2A_nuclei_2k.pdf

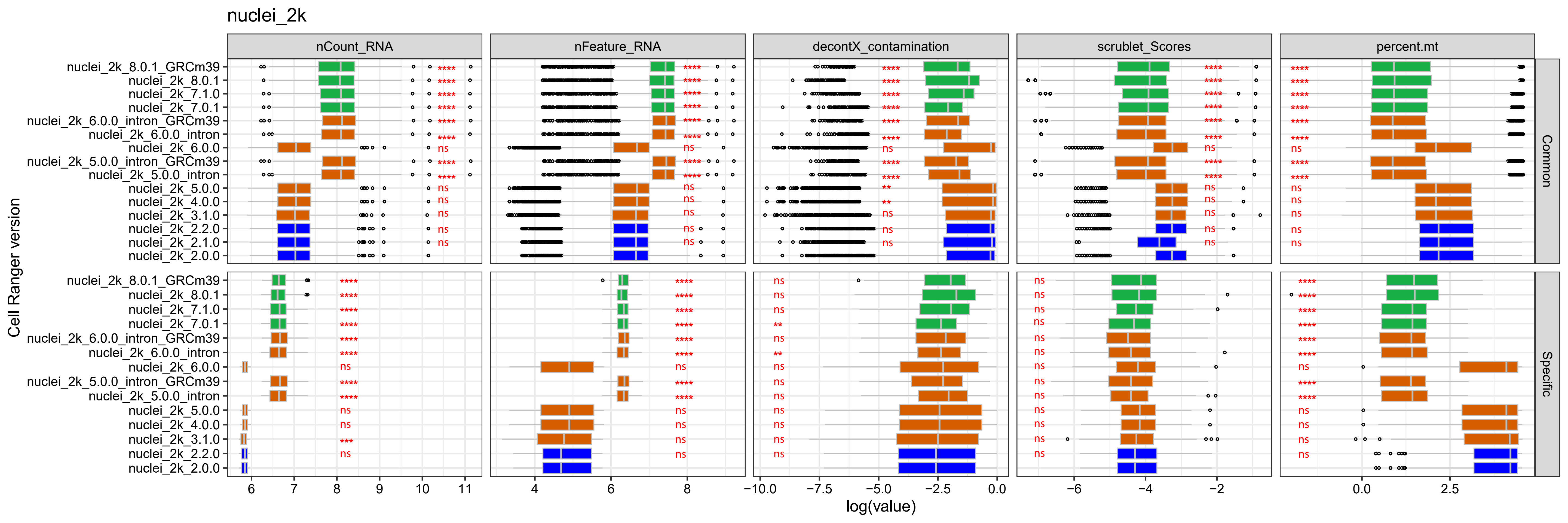

### Supplementary_Figure_2A_nuclei_900.pdf

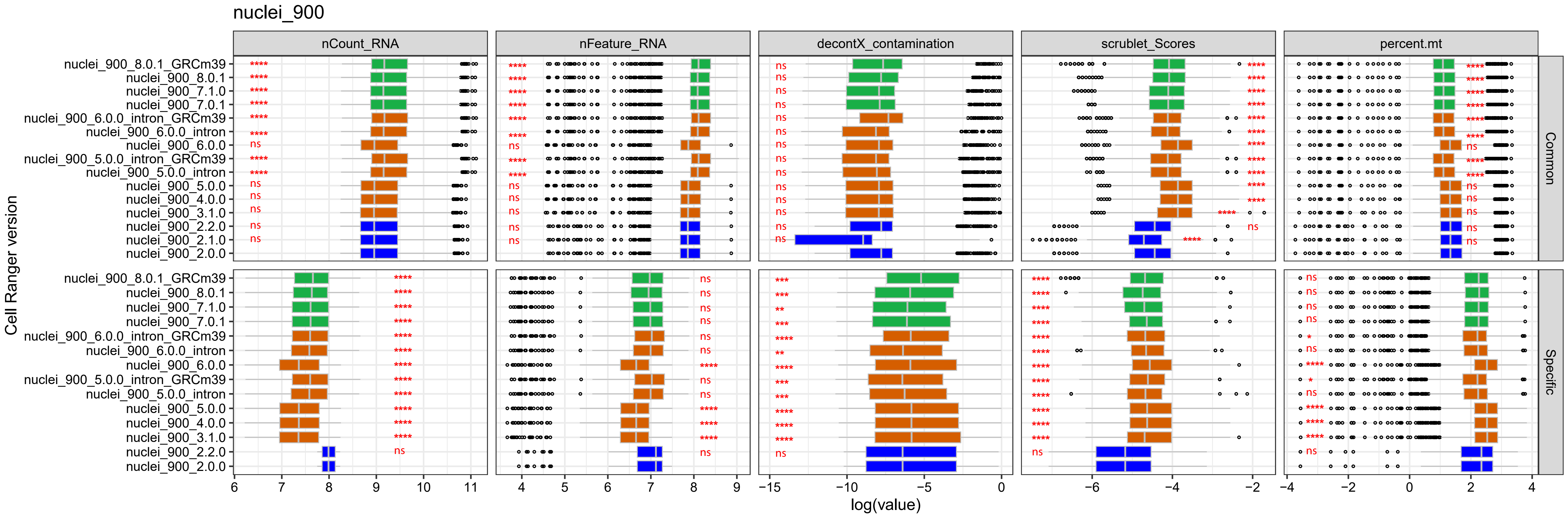

### Supplementary_Figure_2A_pbmc4k.pdf

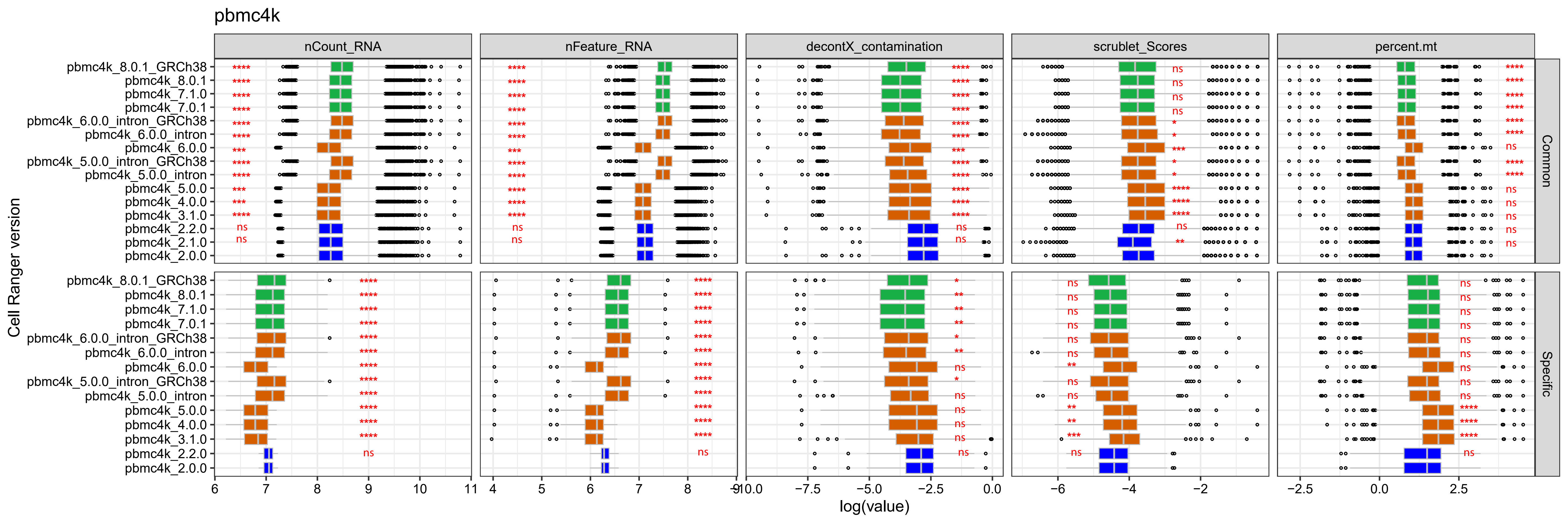

### Supplementary_Figure_2A_t_3k.pdf

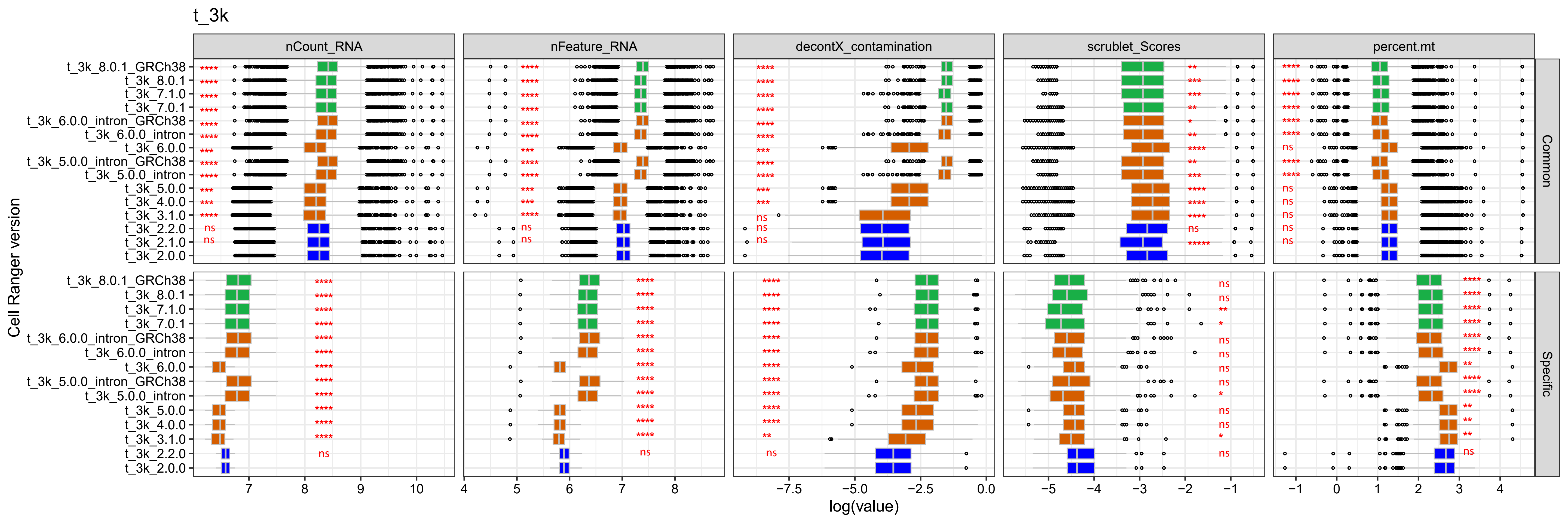

### Supplementary_Figure_2A_t_4k.pdf

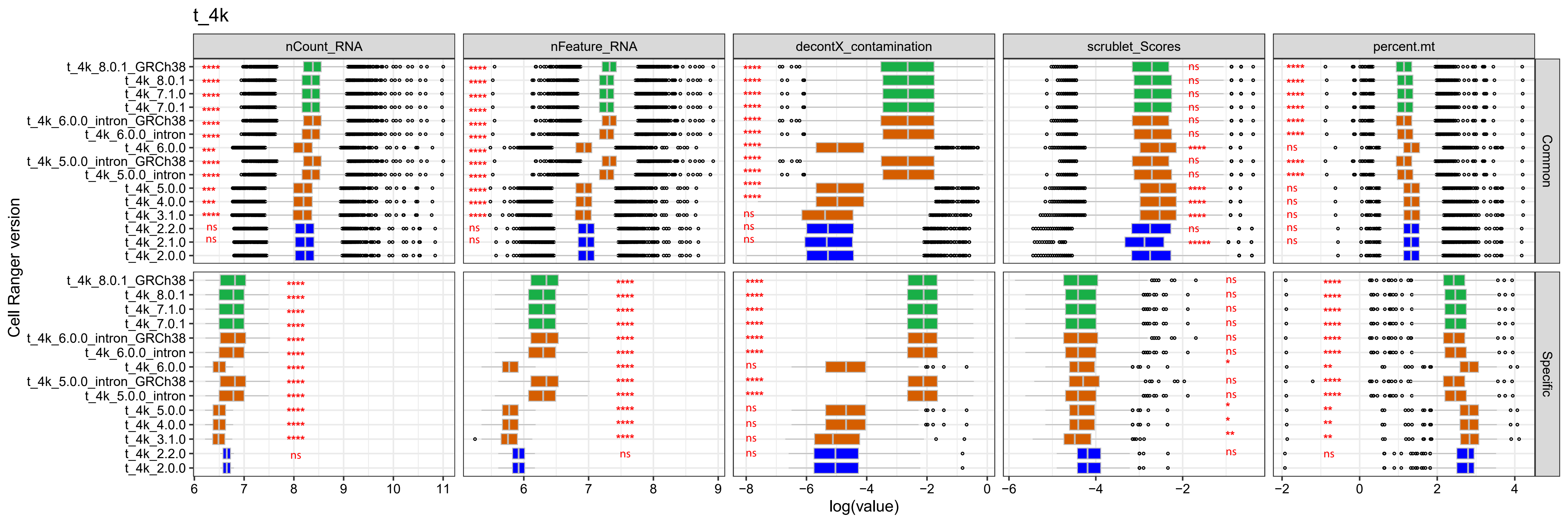

### Supplementary_Figure_2B_fixed_neurons_6days_2000.pdf

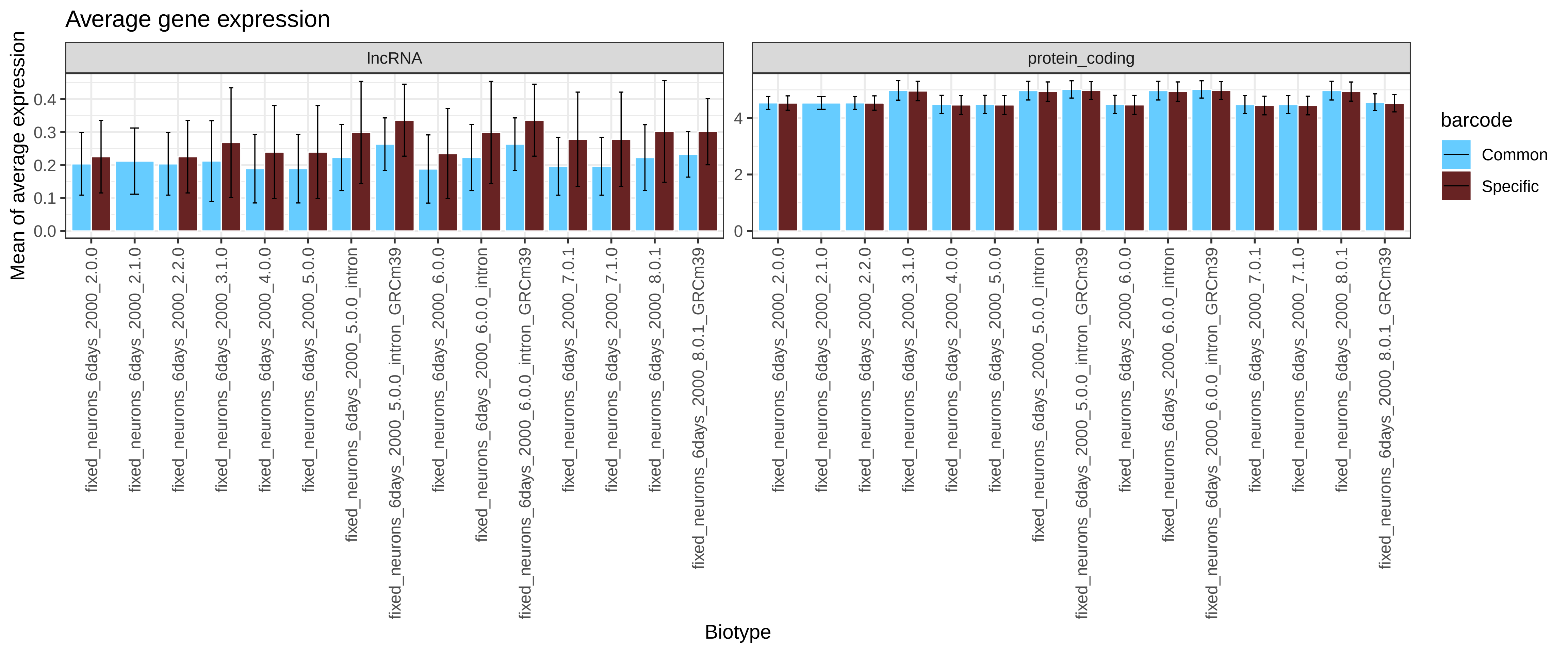

### Supplementary_Figure_2B_fixed_neurons_2000.pdf

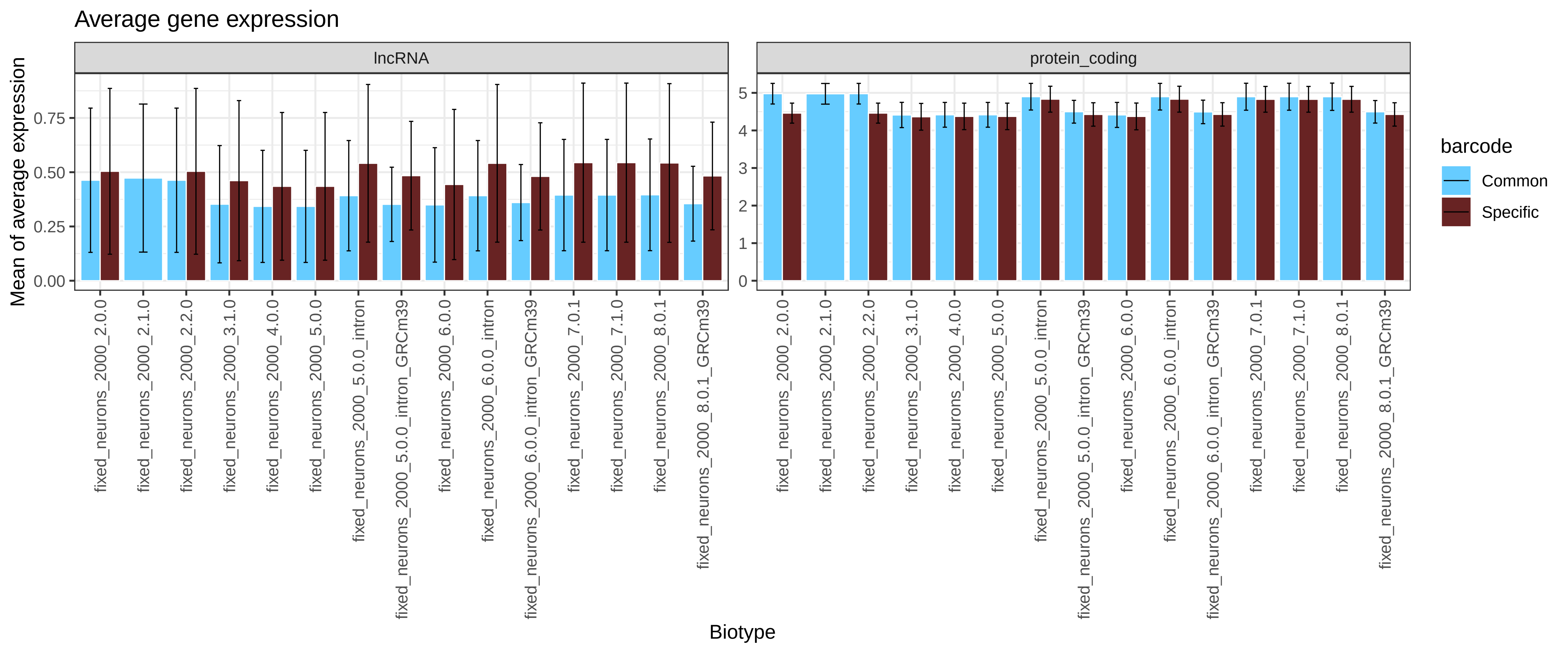

### Supplementary_Figure_2B_GSE143607.pdf

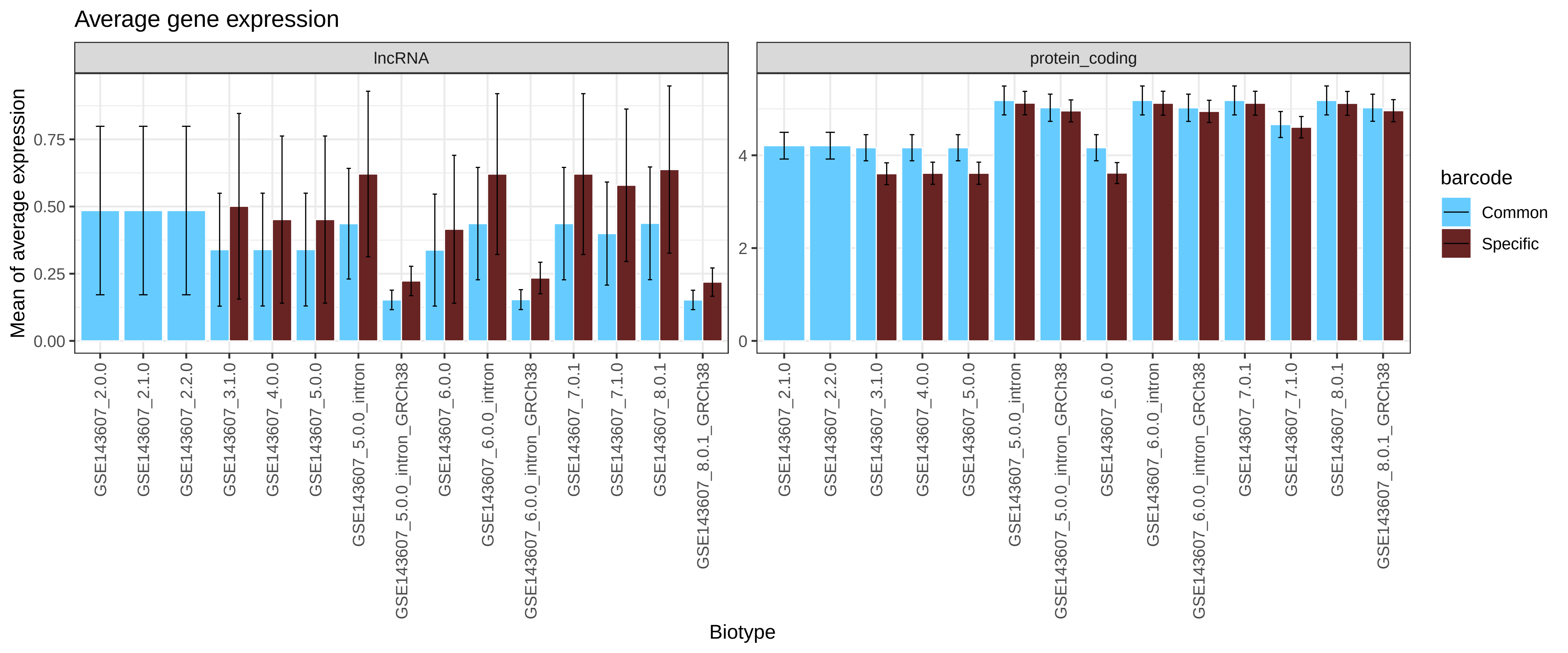

### Supplementary_Figure_2B_neuron_9k.pdf

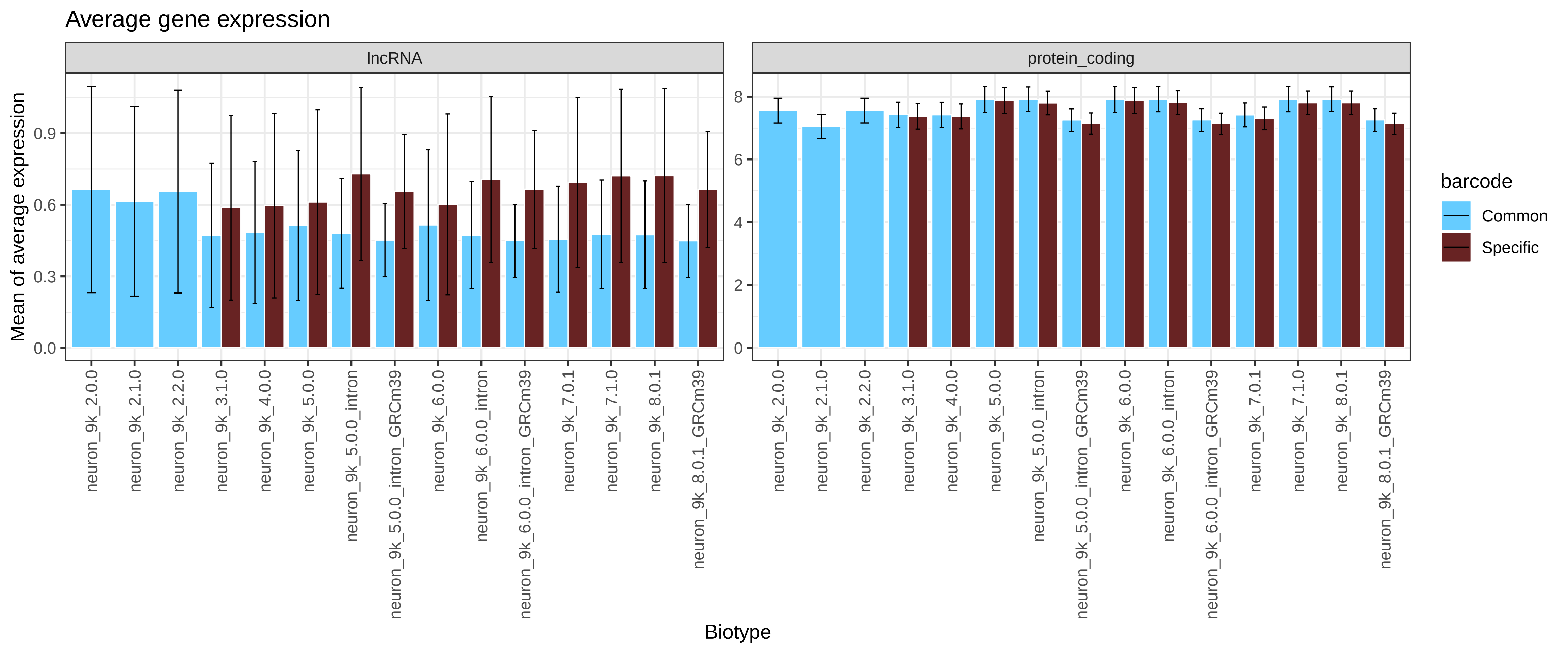

### Supplementary_Figure_2B_neurons_900.pdf

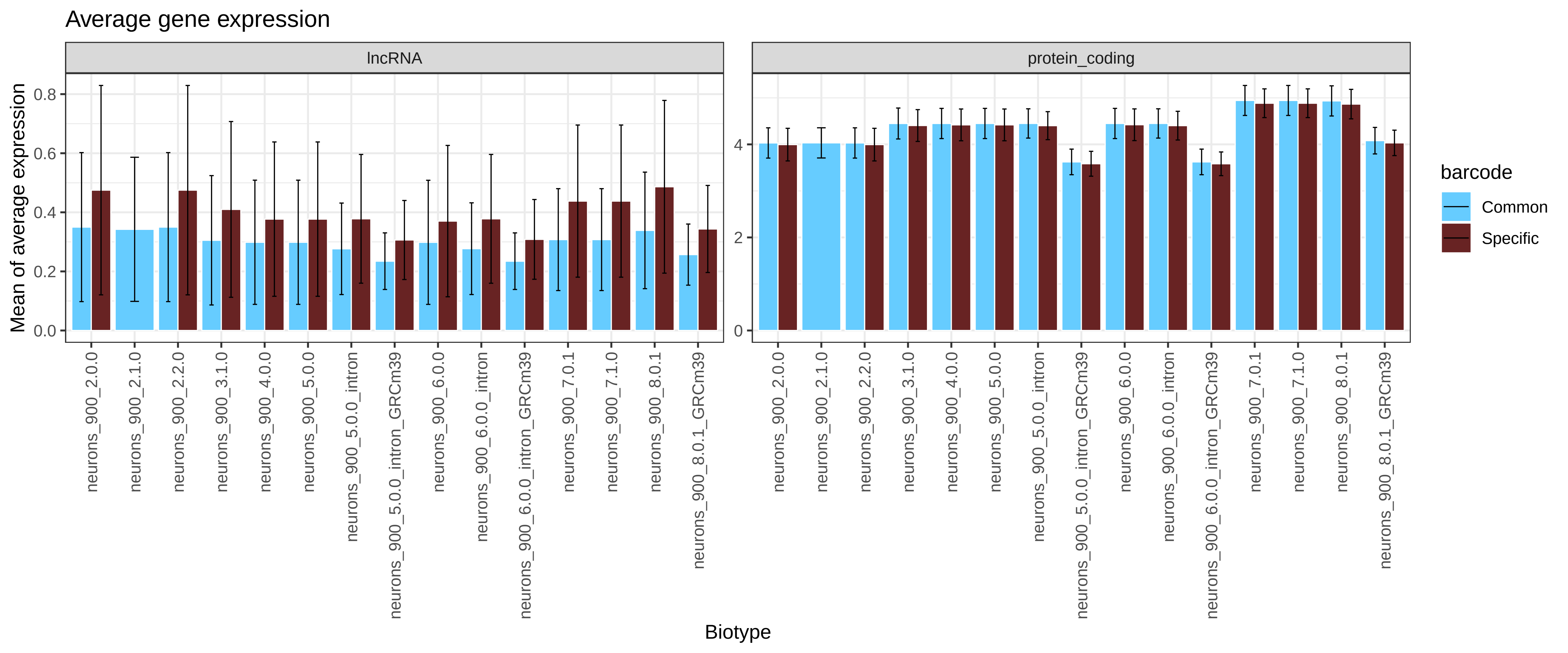

### Supplementary_Figure_2B_neurons_2000.pdf

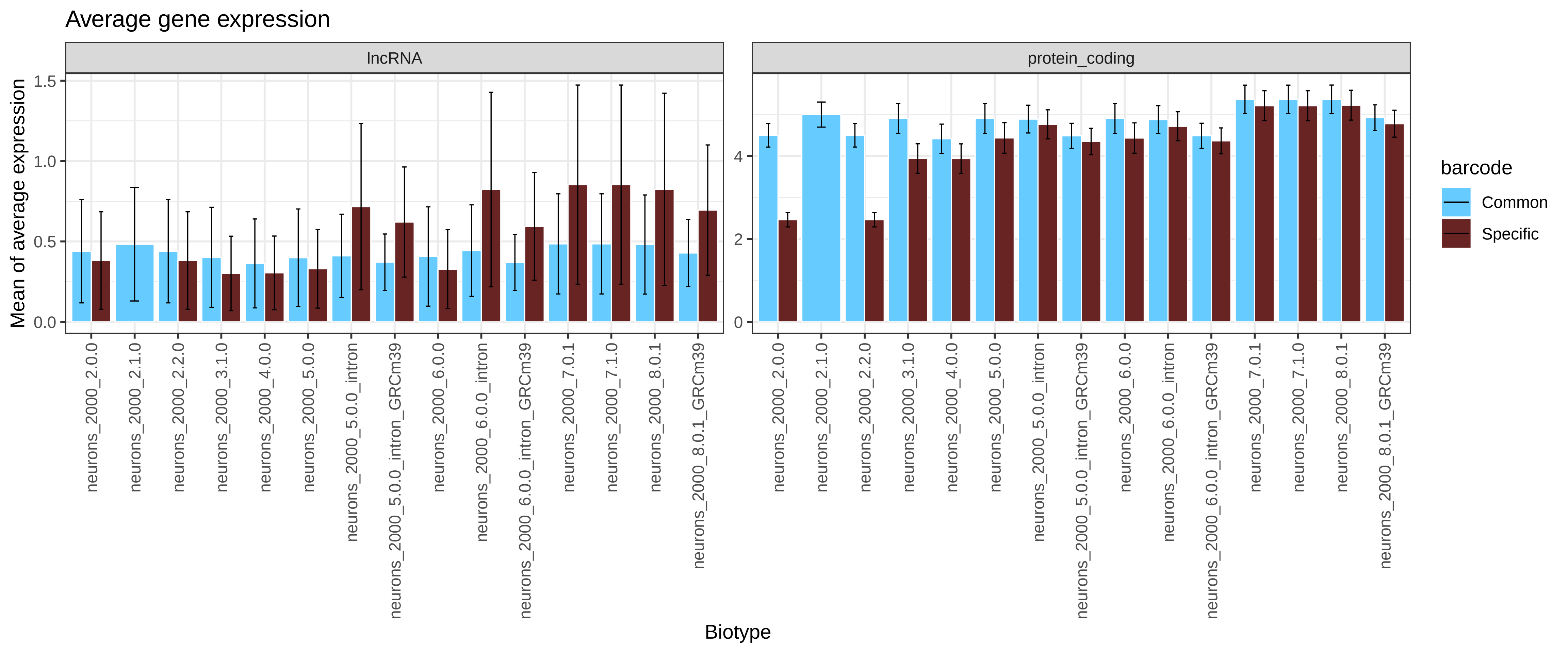

### Supplementary_Figure_2B_nuclei_2k.pdf

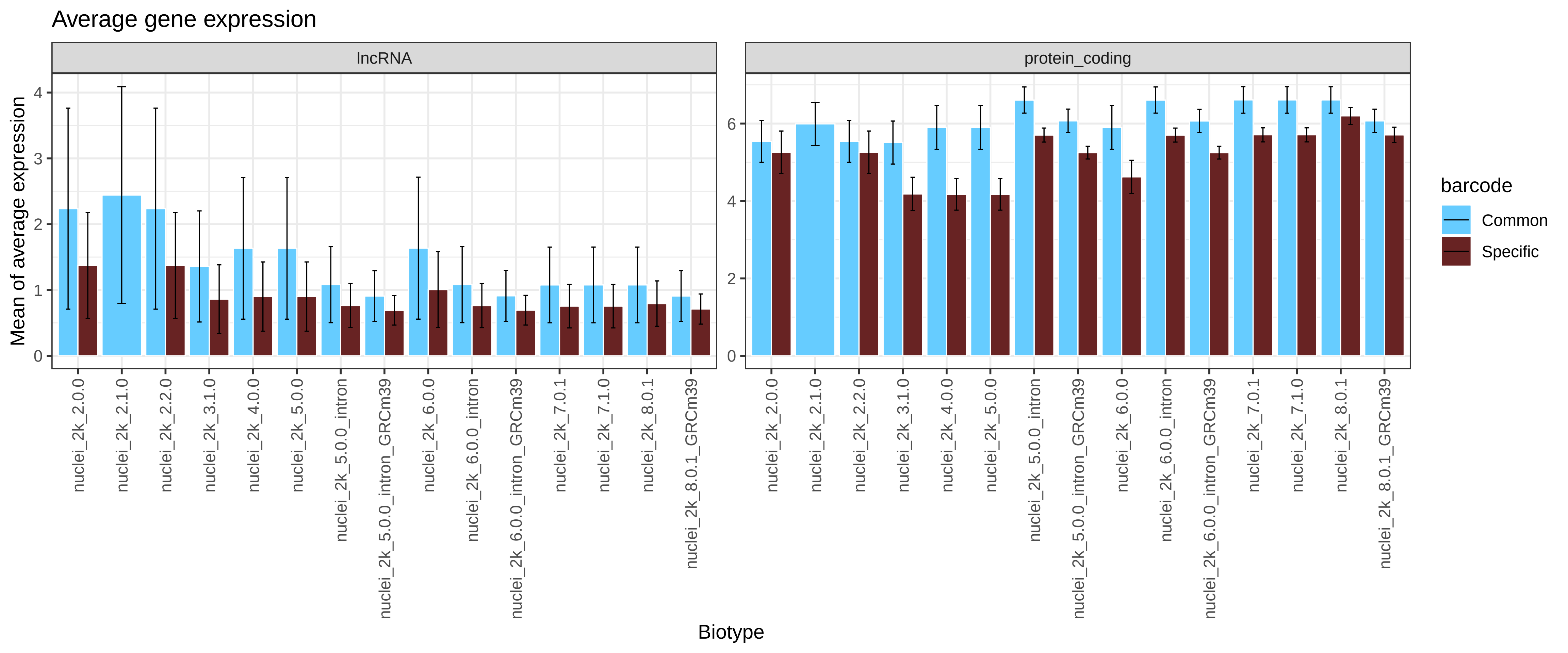

### Supplementary_Figure_2B_nuclei_900.pdf

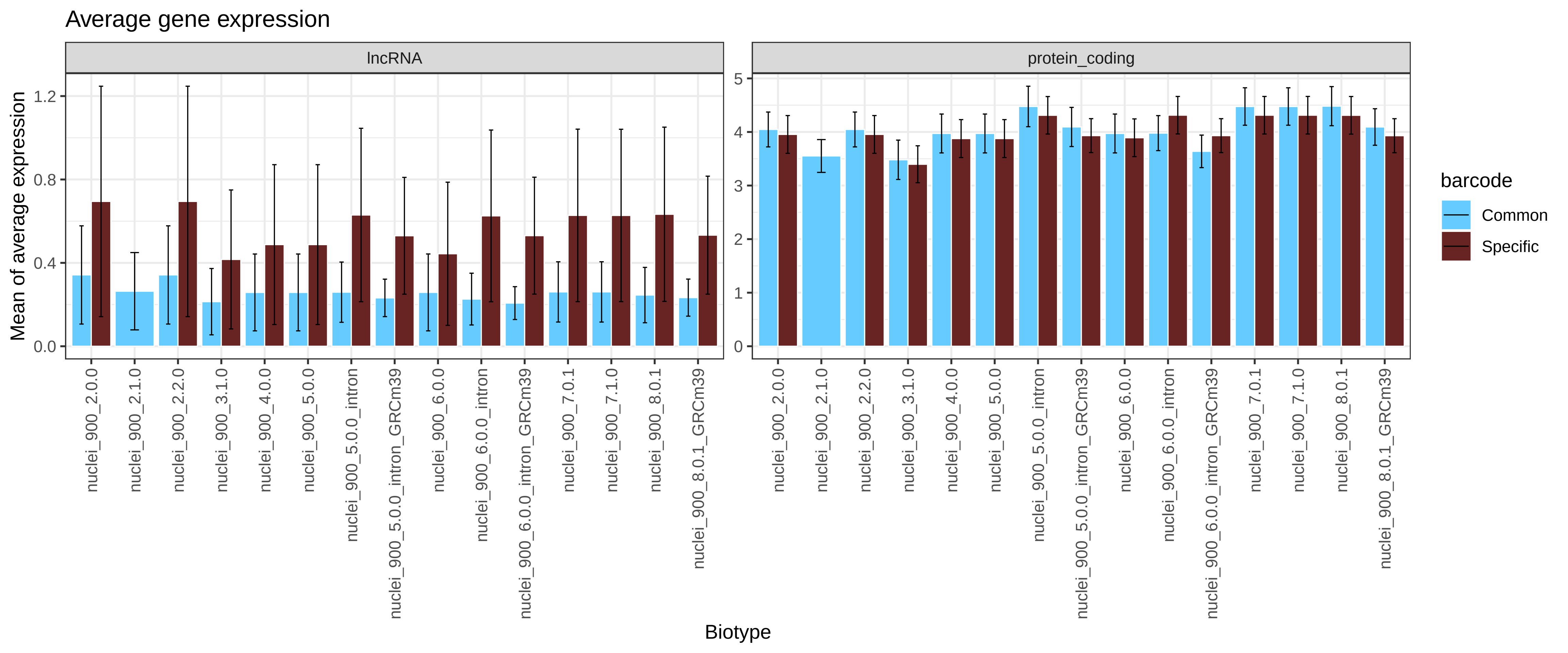

### Supplementary_Figure_2B_pbmc4k.pdf

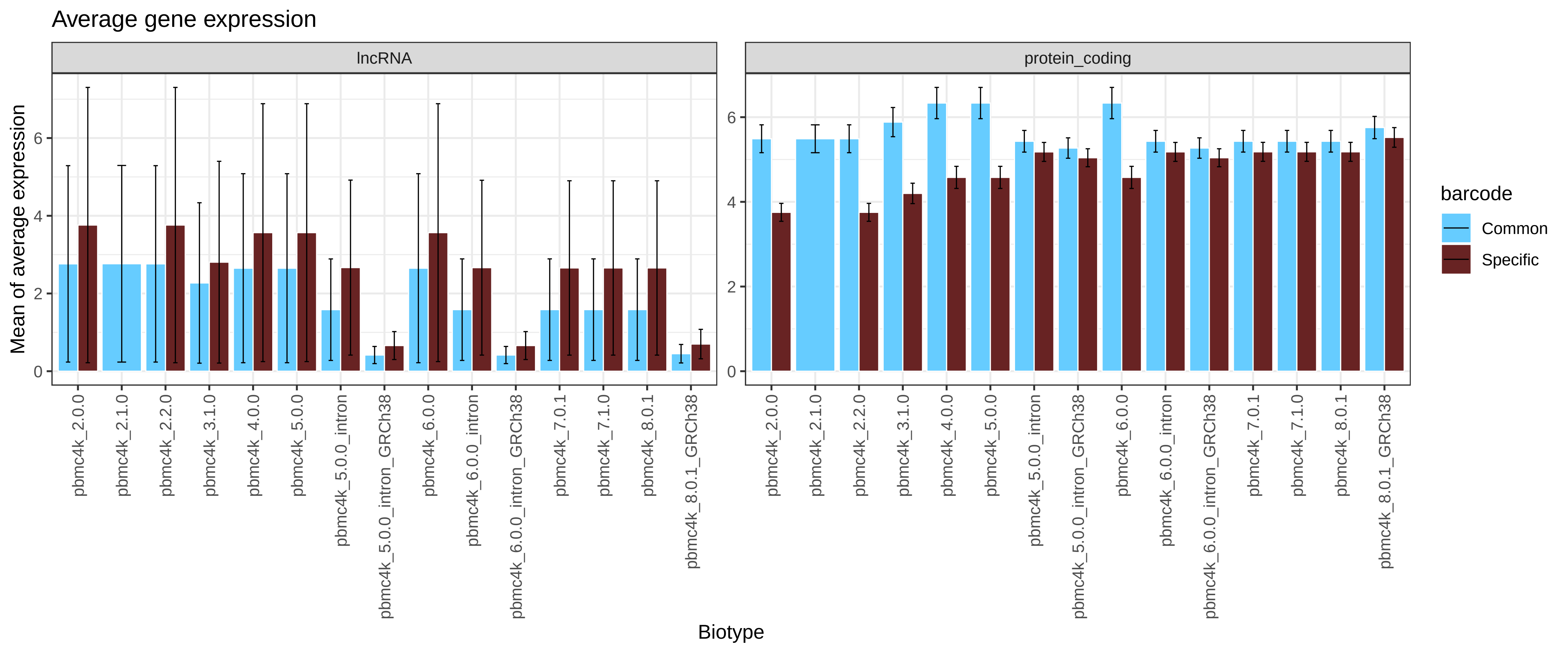

### Supplementary_Figure_2B_t_3k.pdf

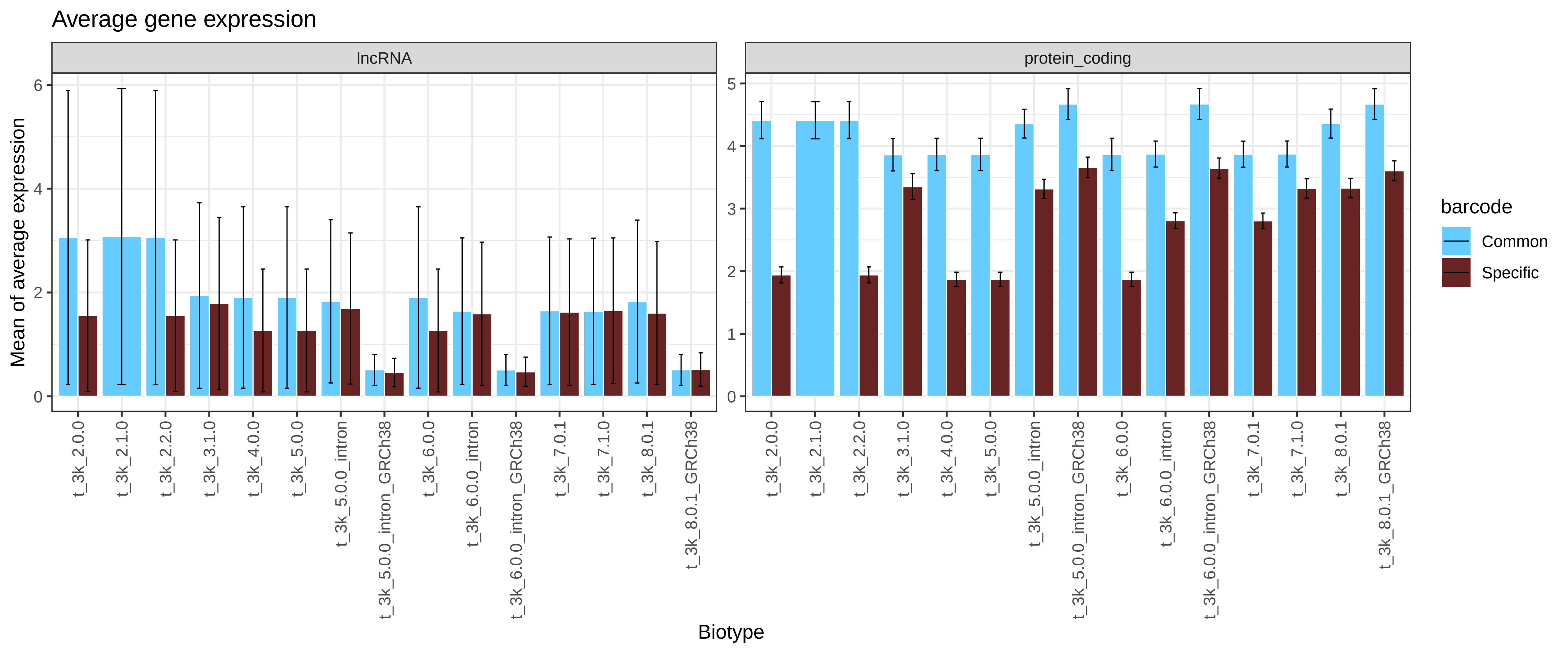

### Supplementary_Figure_2B_t_4k.pdf

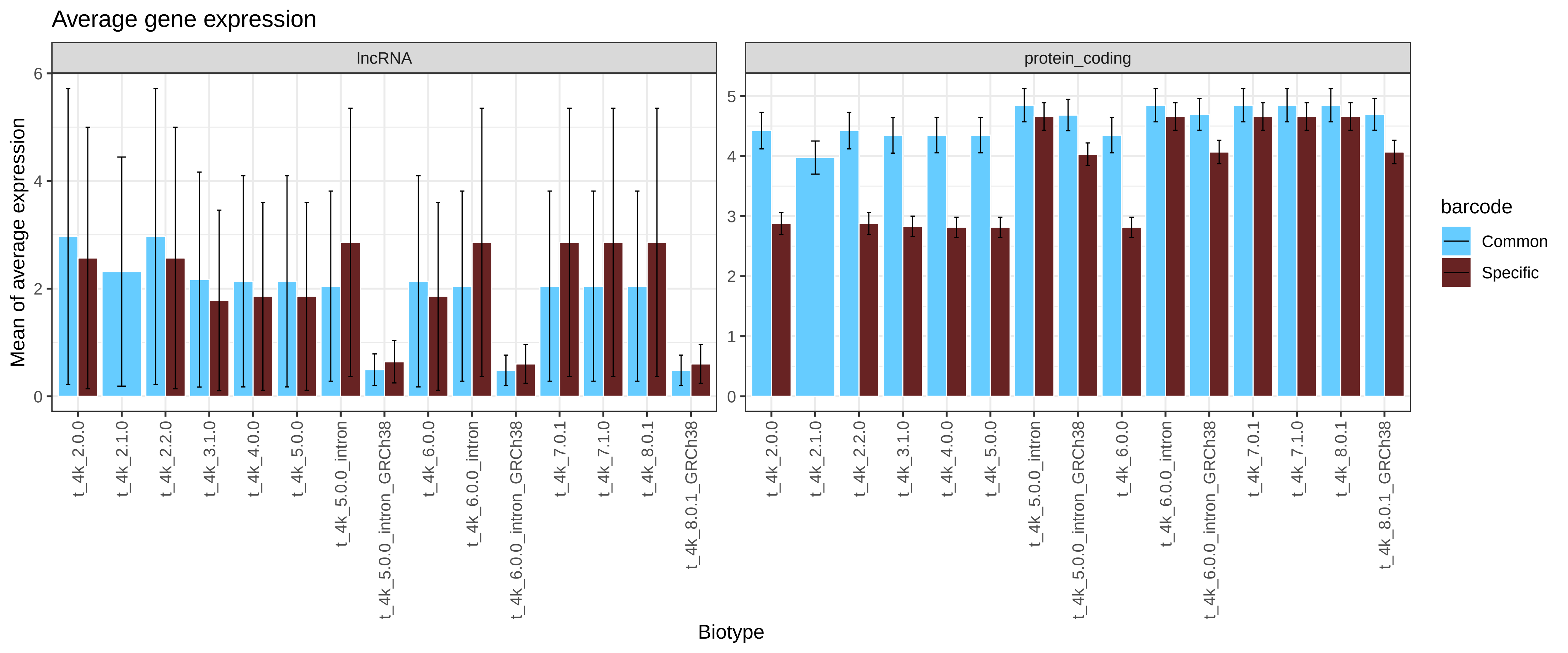

### Supplementary_Figure_3_fixed_neurons_6days_2000.pdf

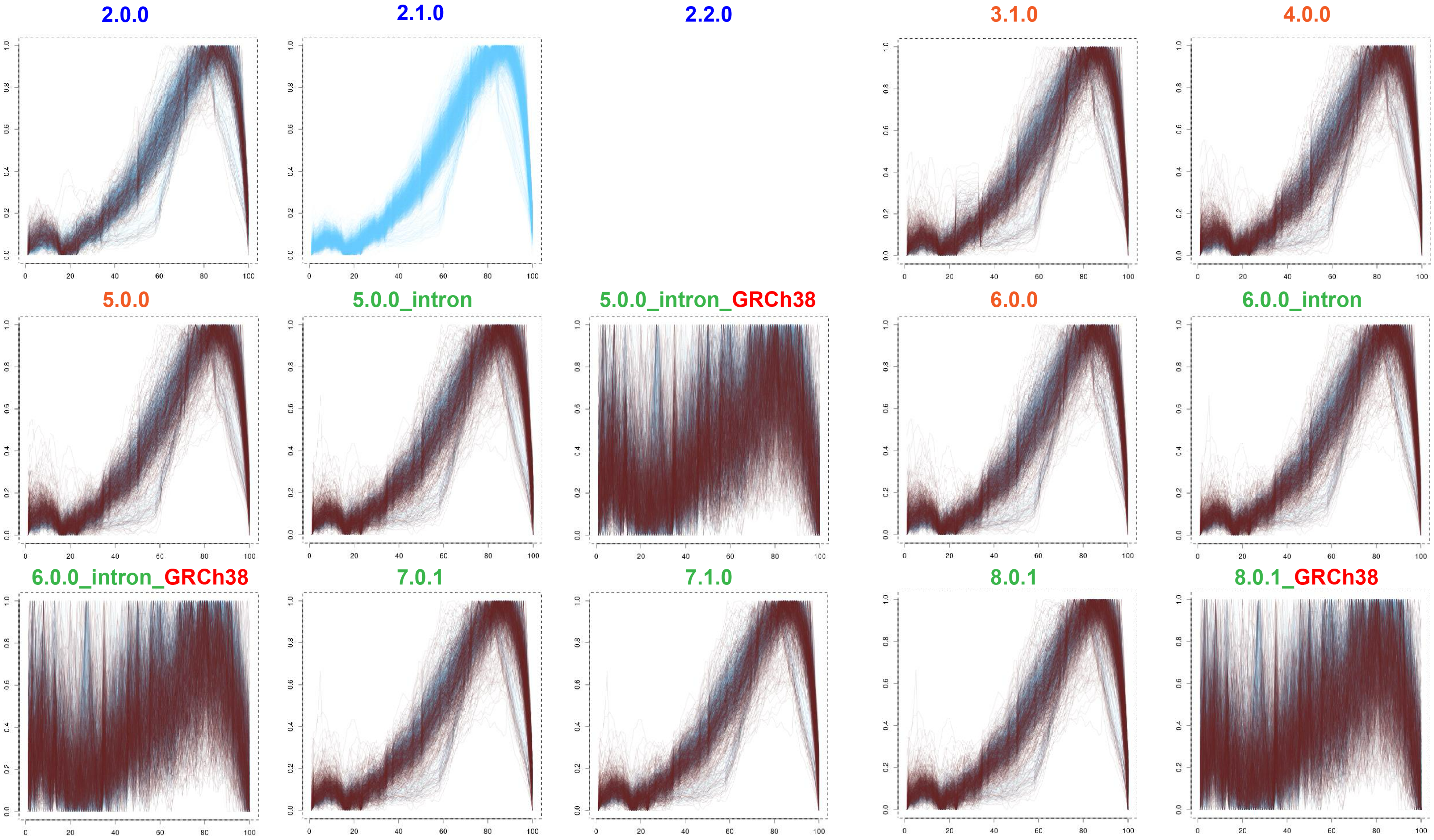

### Supplementary_Figure_3_fixed_neurons_2000.pdf

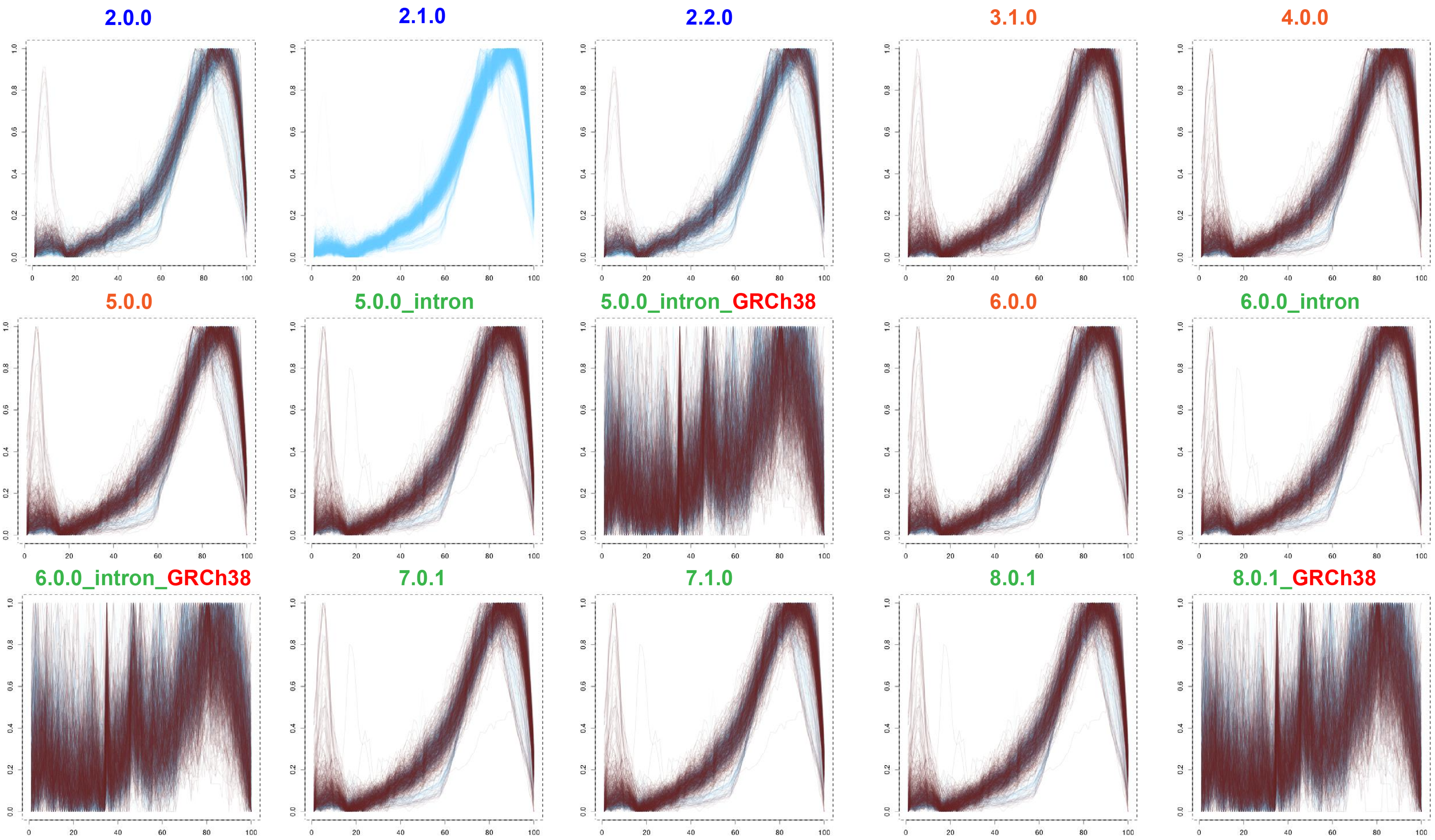

### Supplementary_Figure_3_neuron_9k.pdf

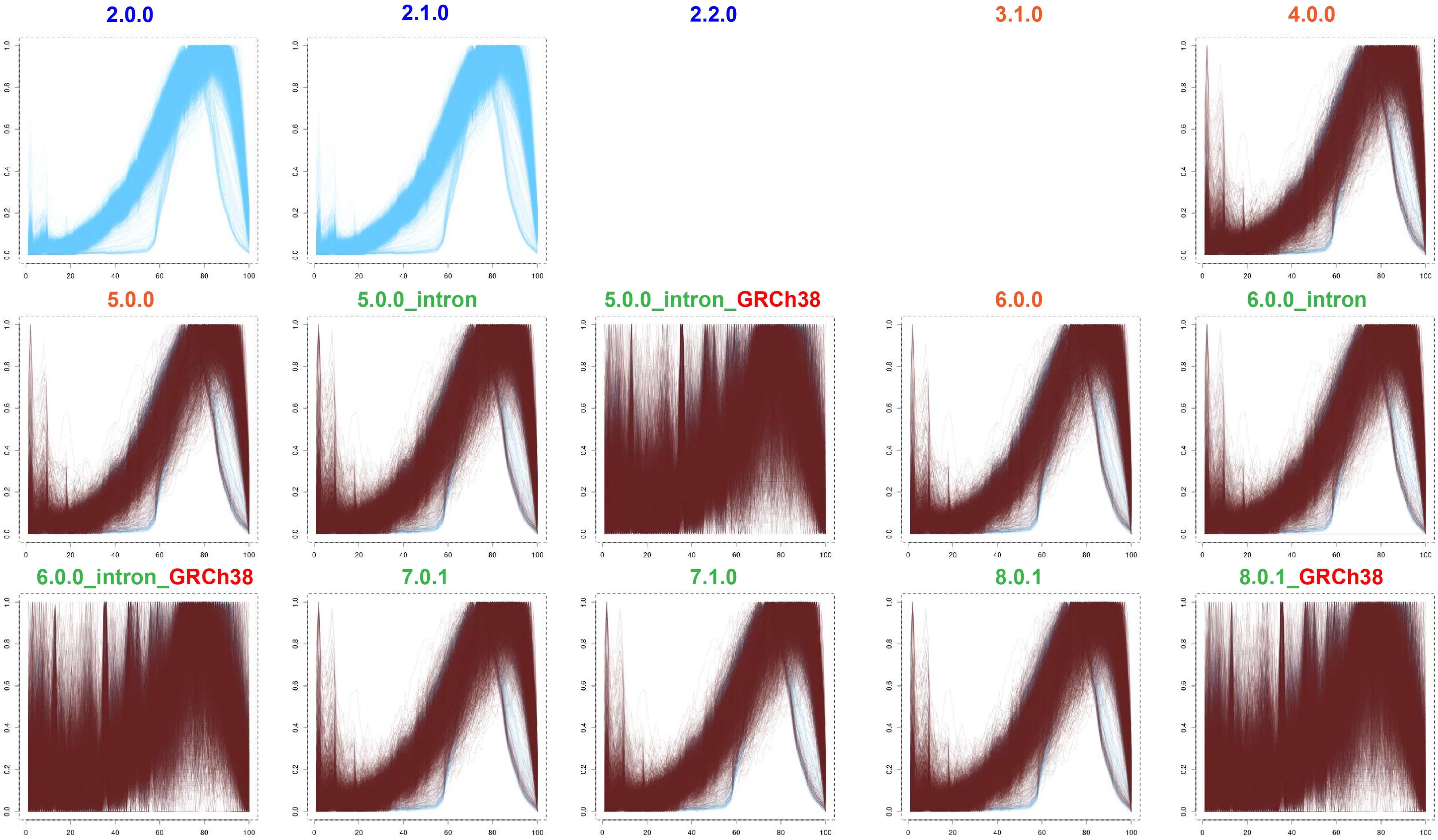

### Supplementary_Figure_3_neurons_900.pdf

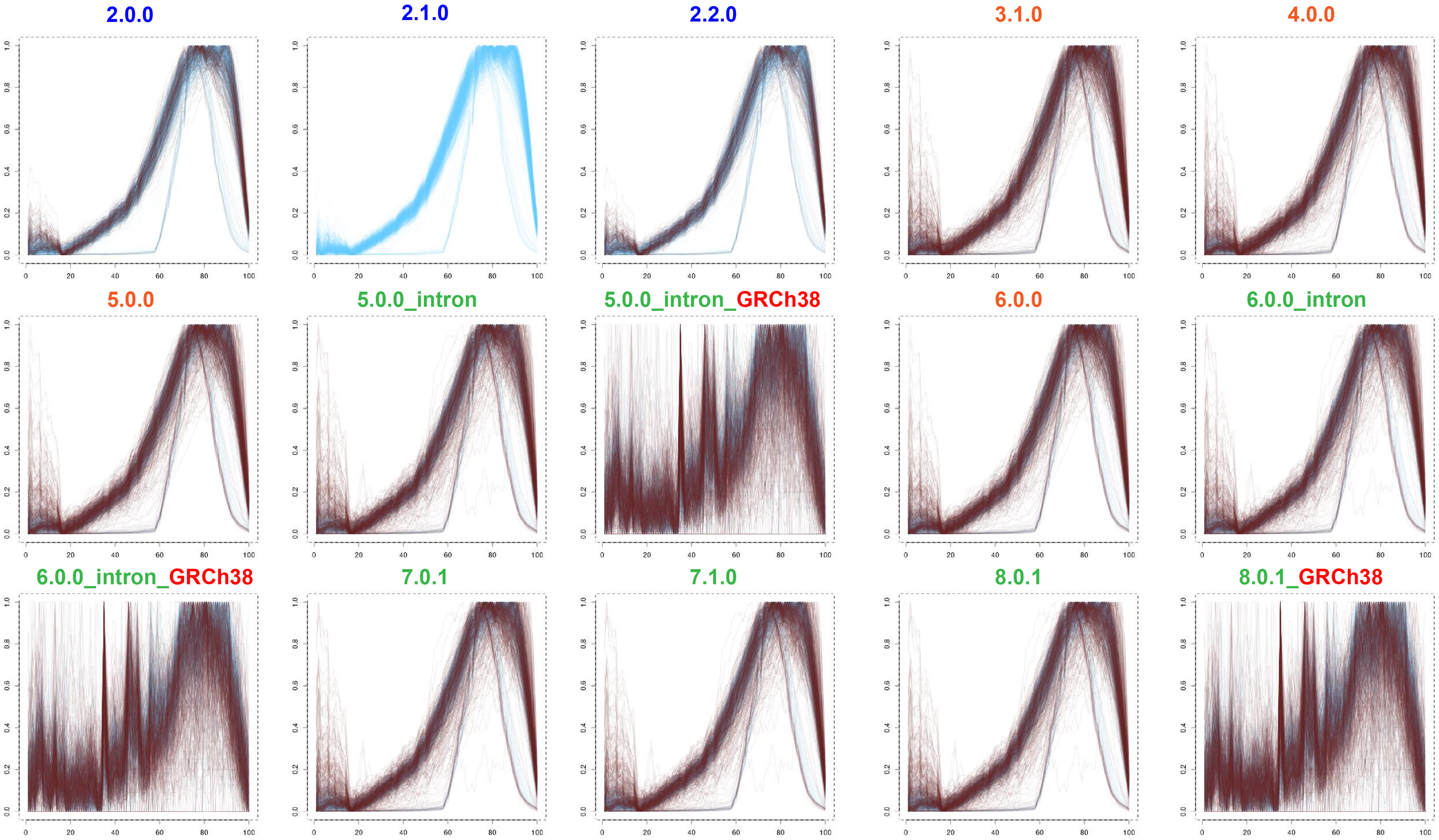

### Supplementary_Figure_3_neurons_2000.pdf

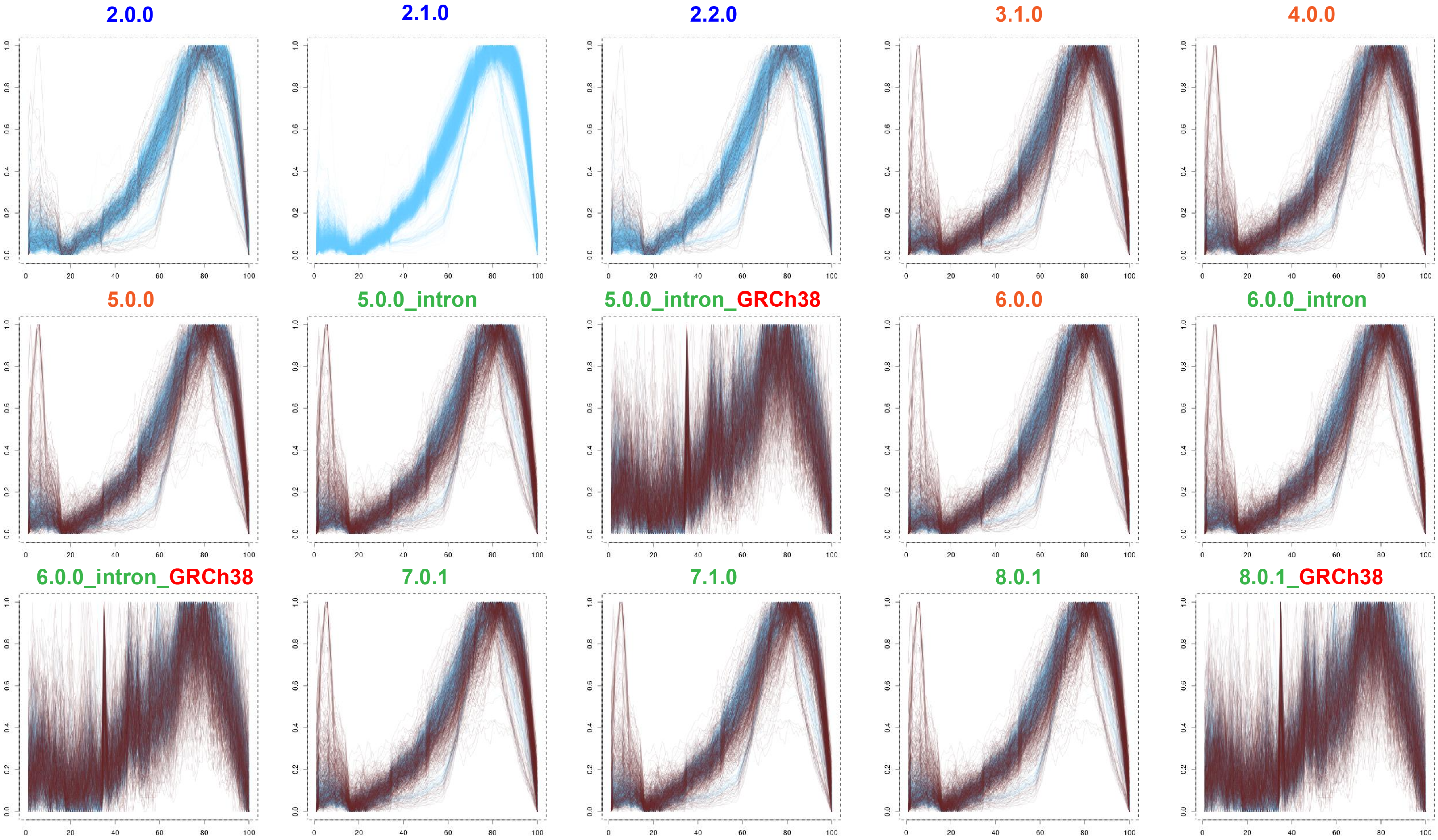

### Supplementary_Figure_3_nuclei_2k.pdf

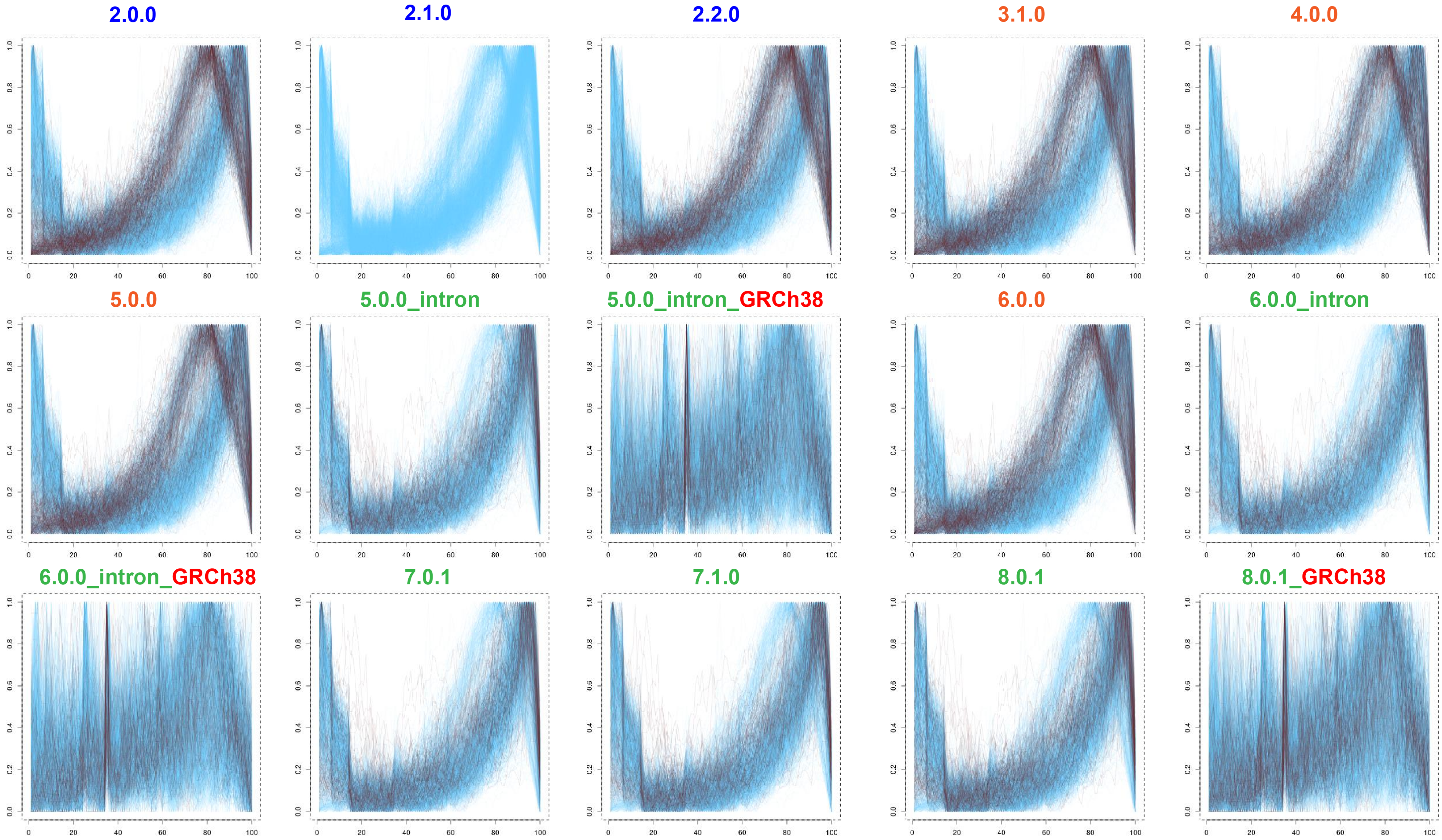

### Supplementary_Figure_3_nuclei_900.pdf

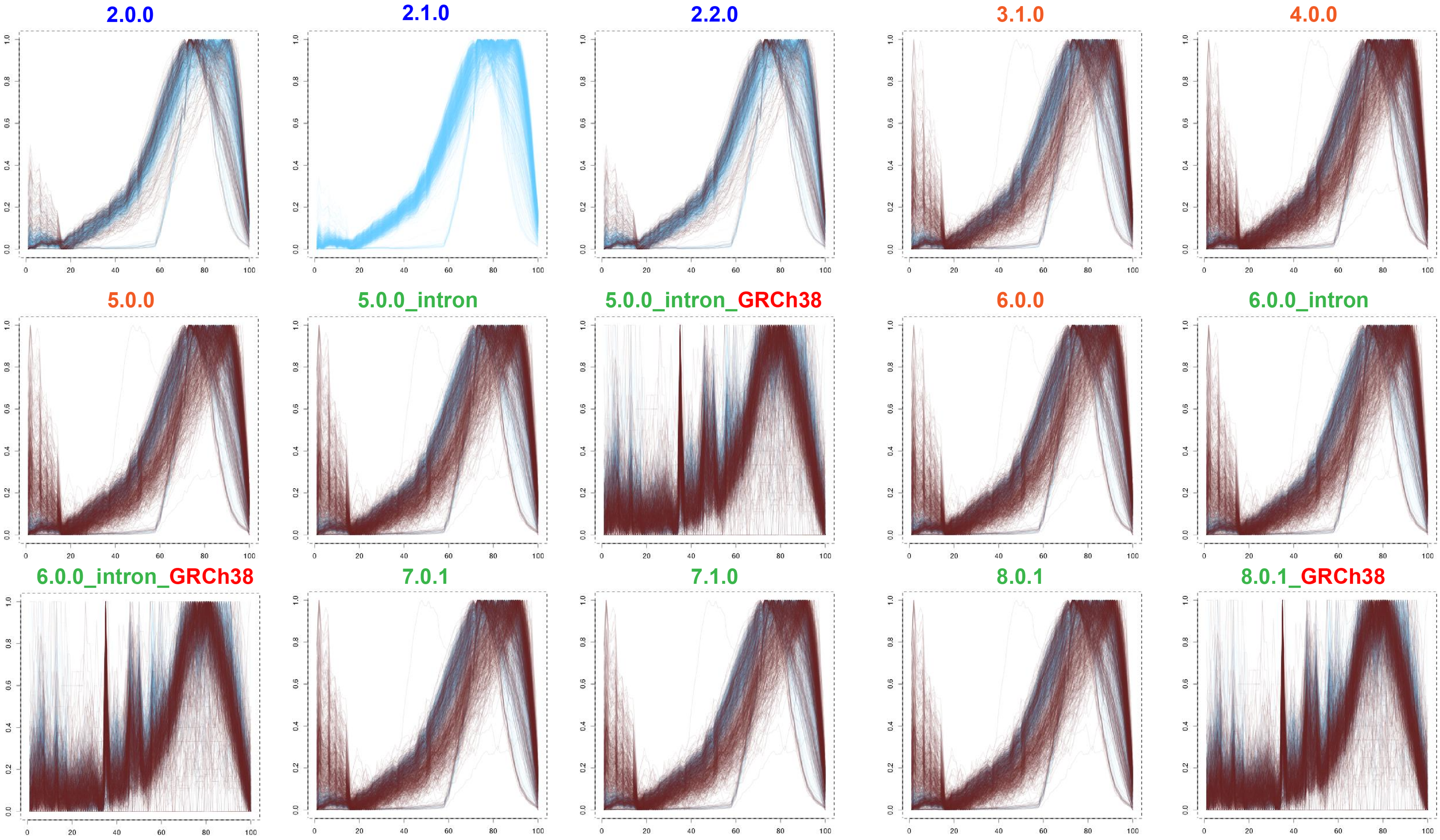

### Supplementary_Figure_3_pbmc4k.pdf

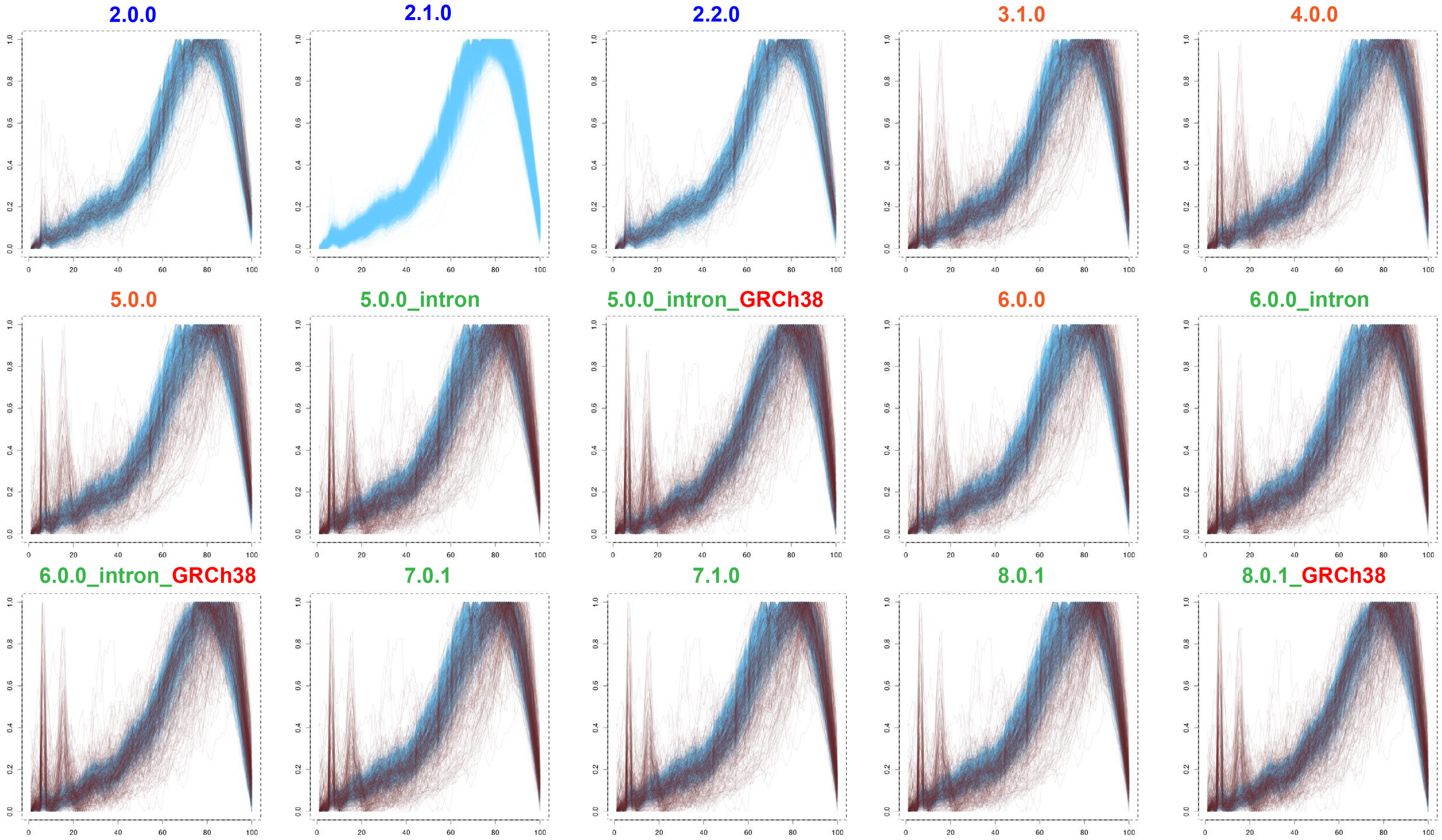

### Supplementary_Figure_3_pbmc8k.pdf

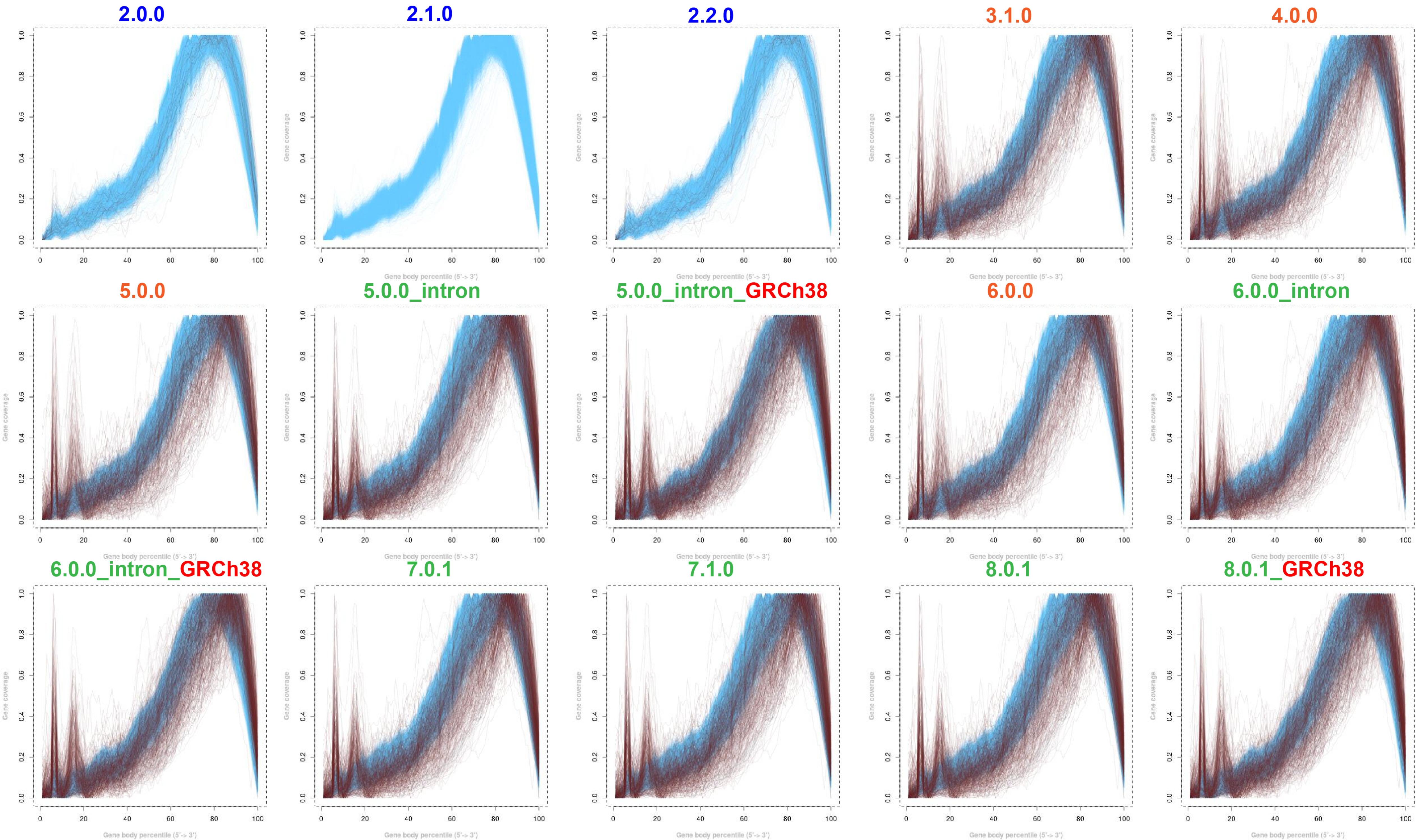

### Supplementary_Figure_3_t_3k.pdf

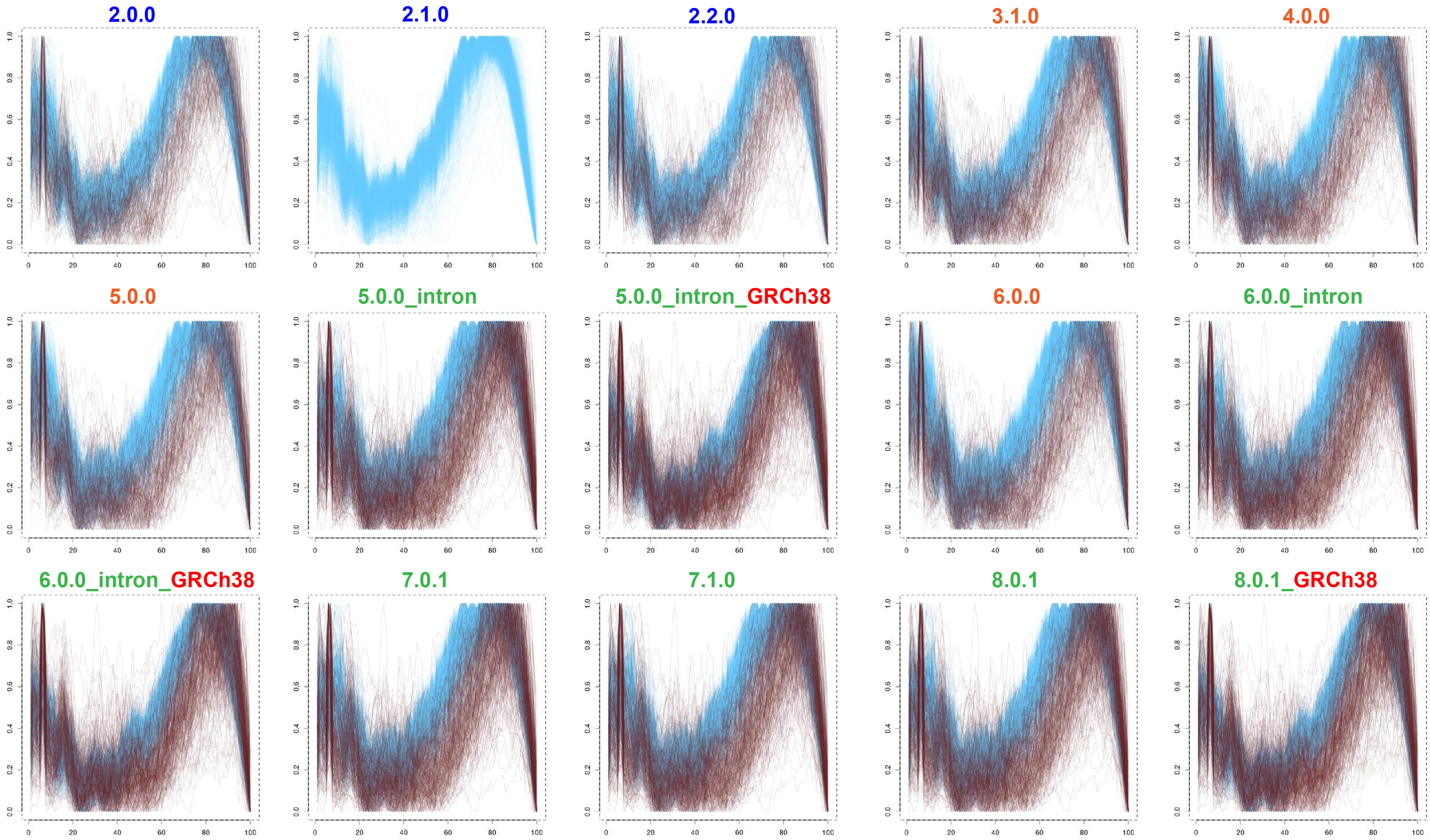

### Supplementary_Figure_3_t_4k.pdf

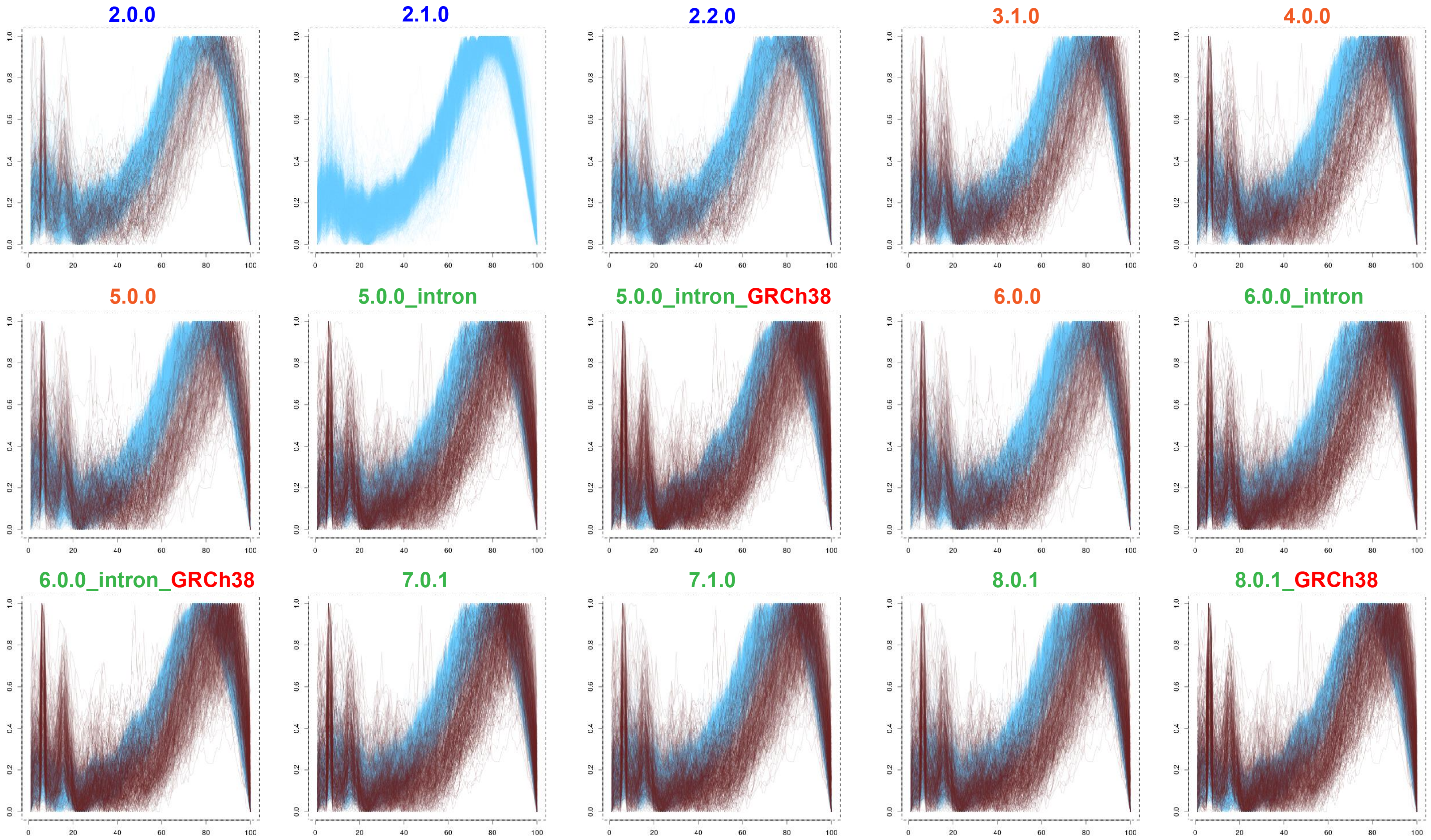

### Supplementary_Figure_4_fixed_neurons_6days_2000.pdf

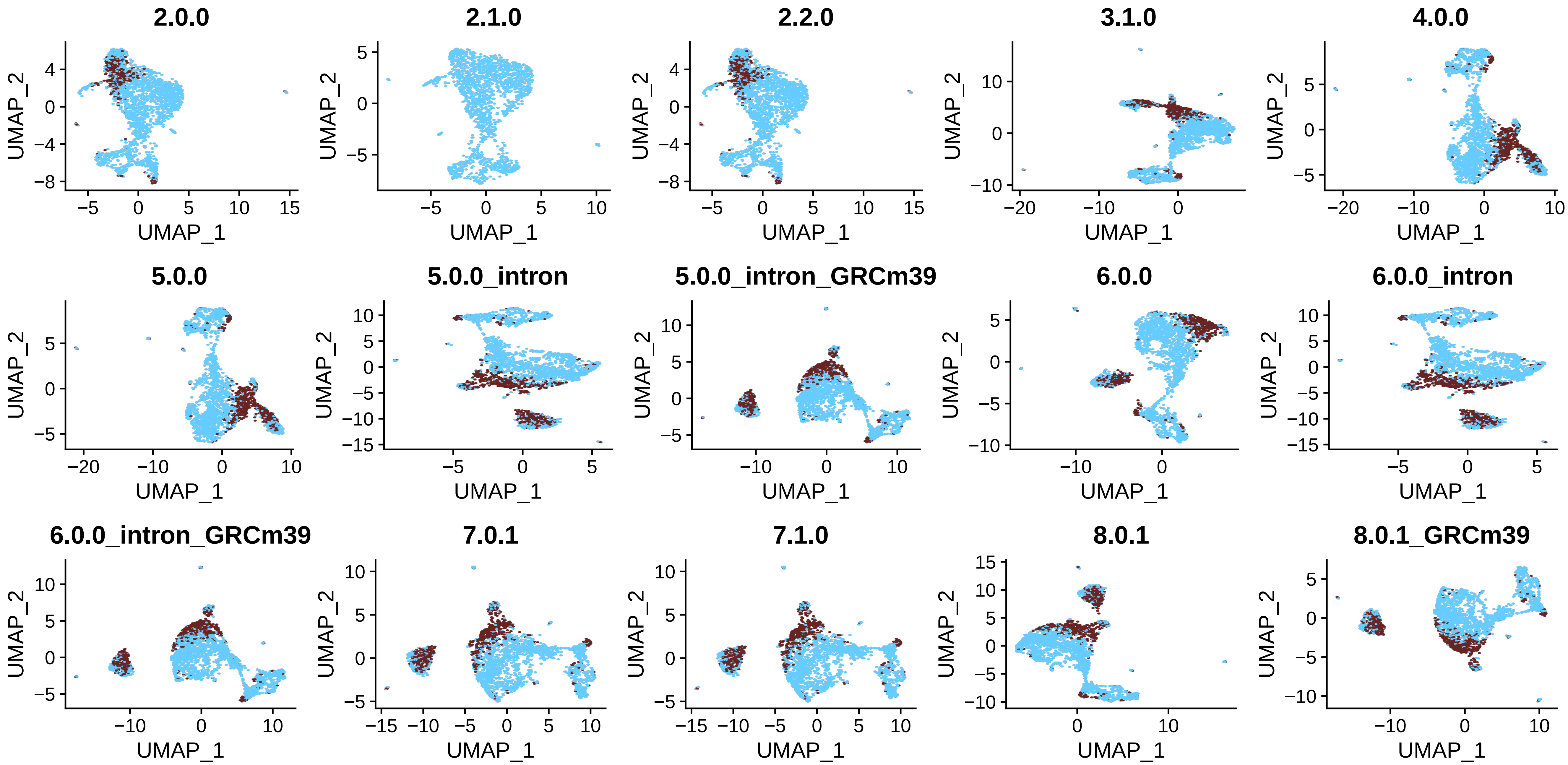

### Supplementary_Figure_4_fixed_neurons_2000.pdf

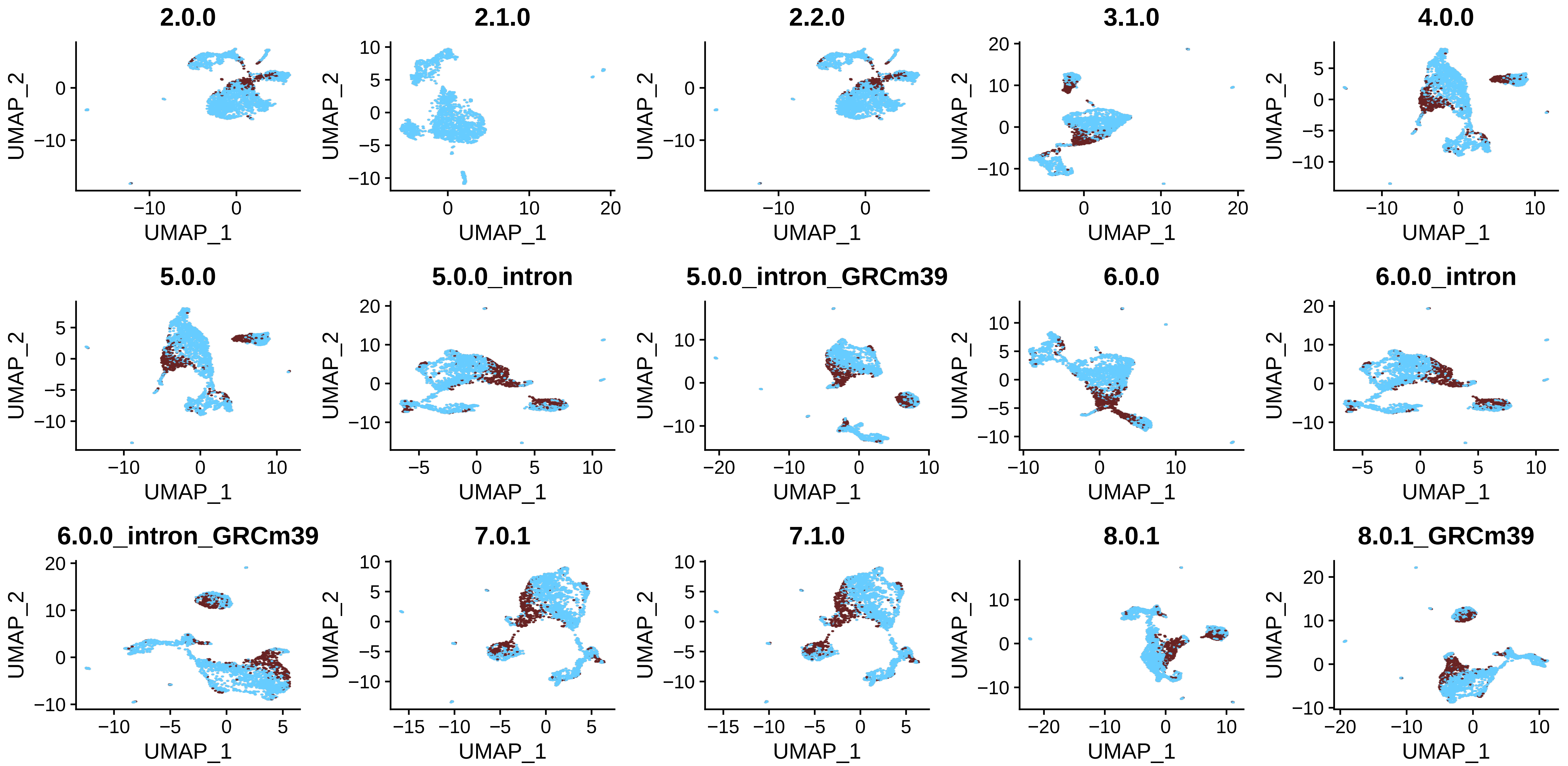

### Supplementary_Figure_4_GSE143607.pdf

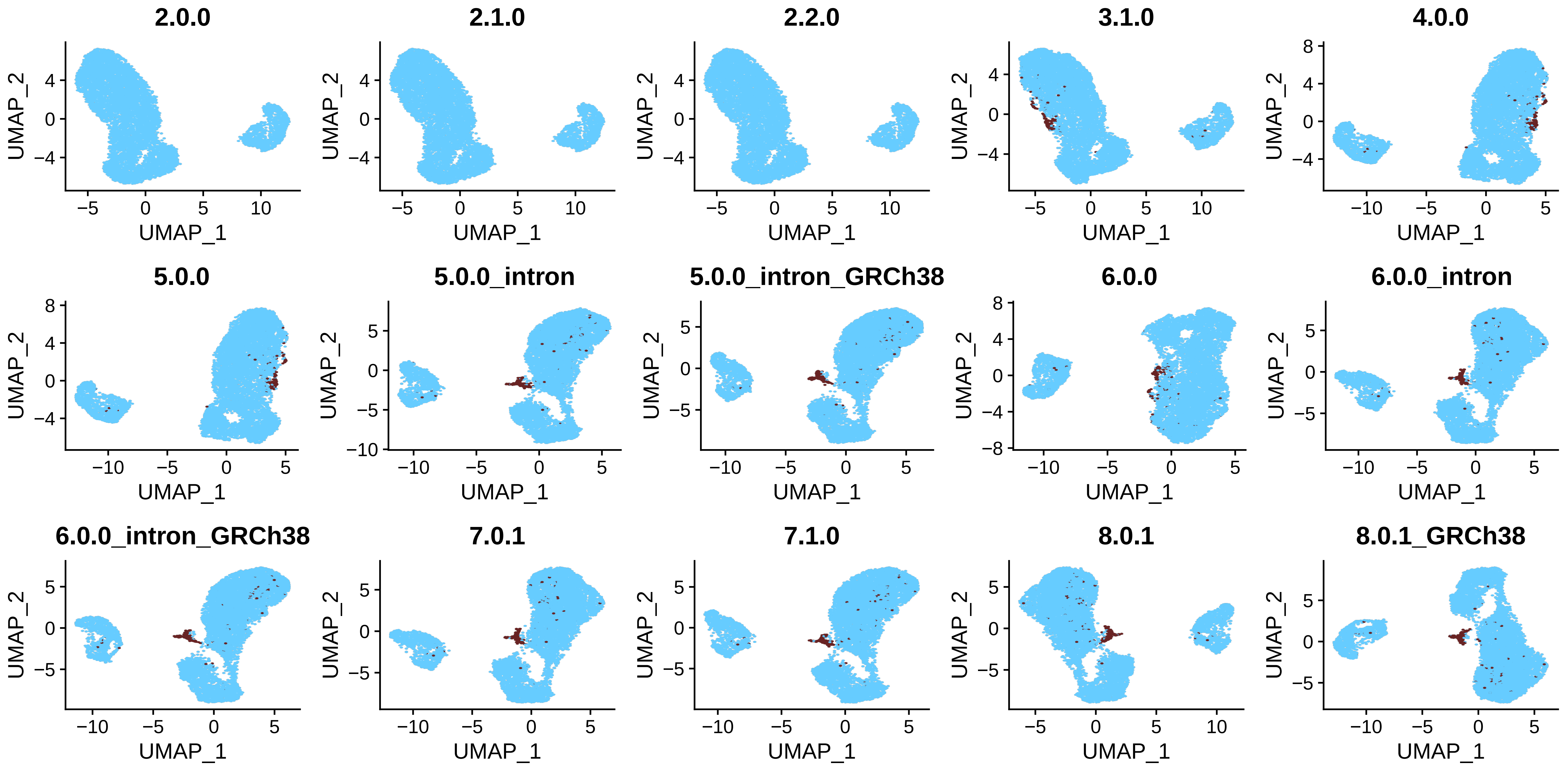

### Supplementary_Figure_4_neuron_9k.pdf

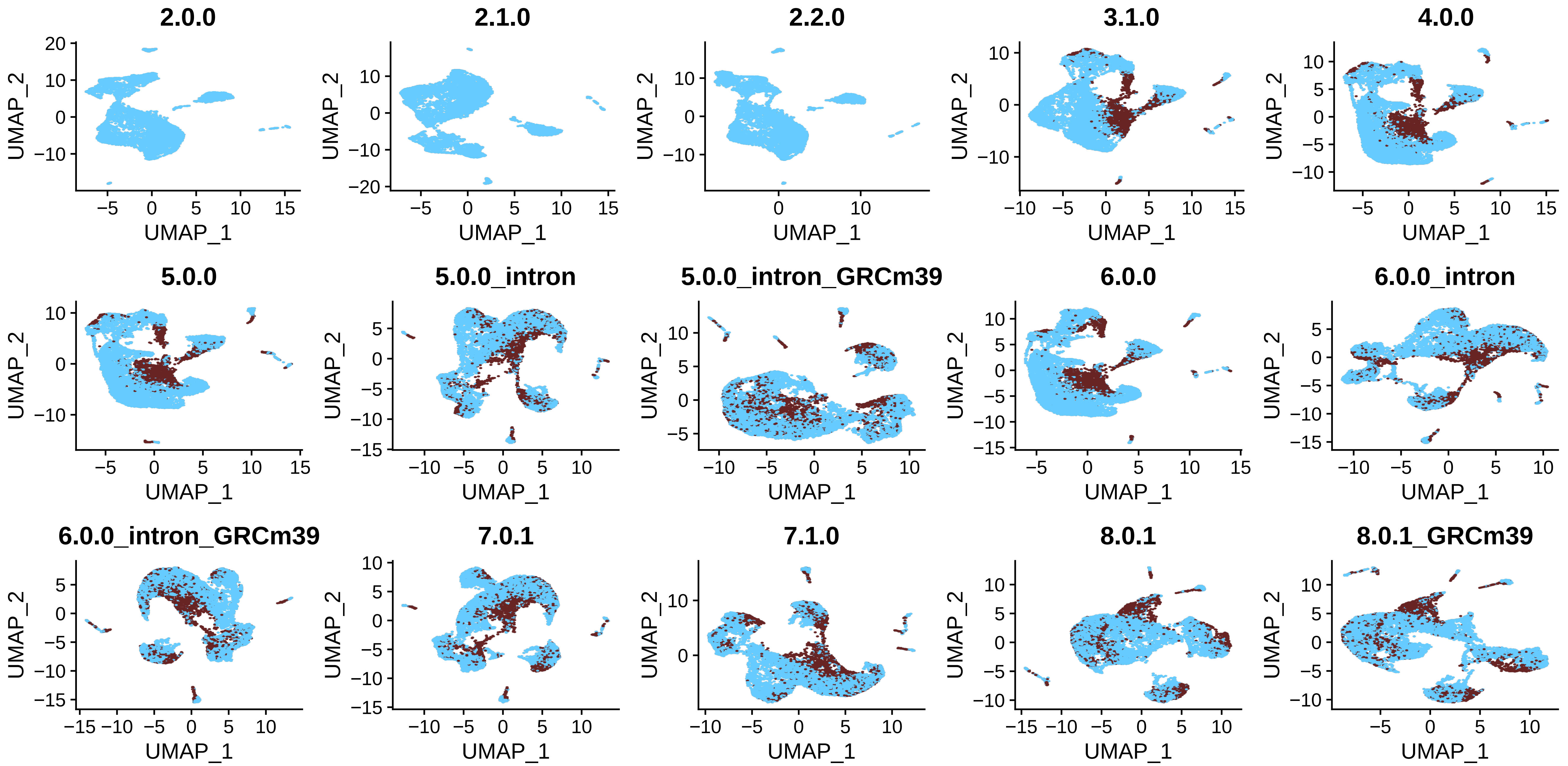

### Supplementary_Figure_4_neurons_900.pdf

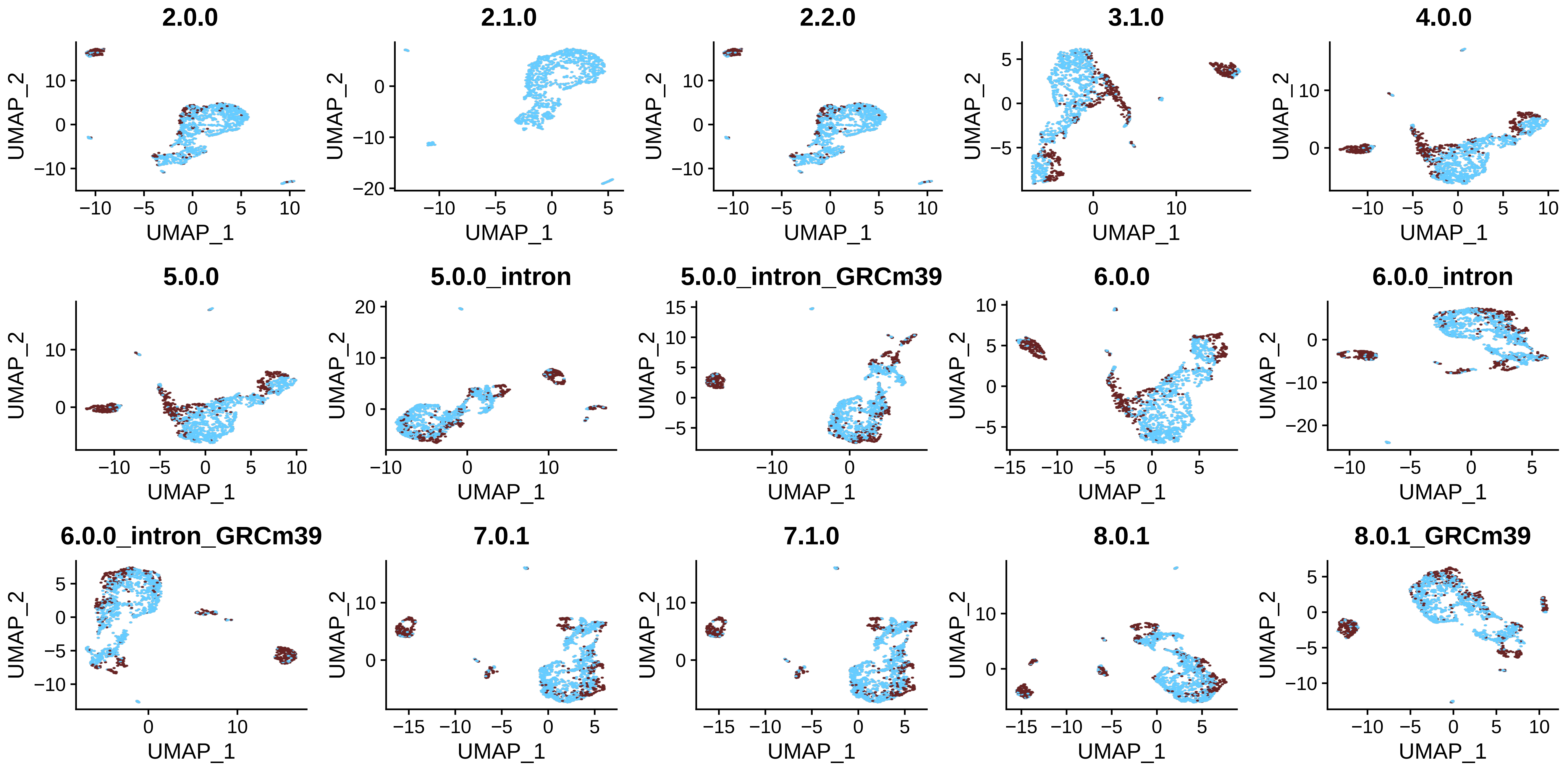

### Supplementary_Figure_4_neurons_2000.pdf

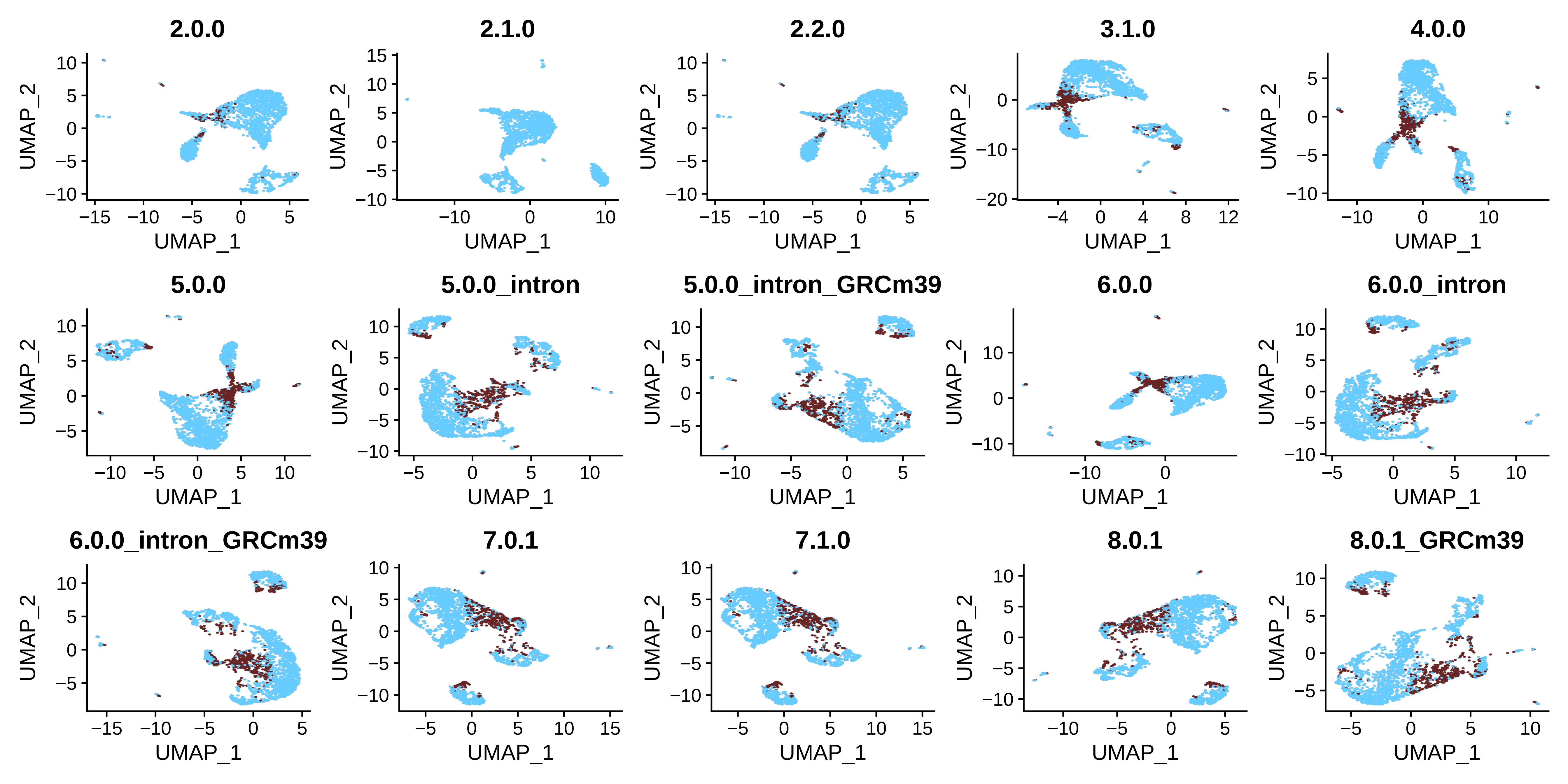

### Supplementary_Figure_4_nuclei_2k.pdf

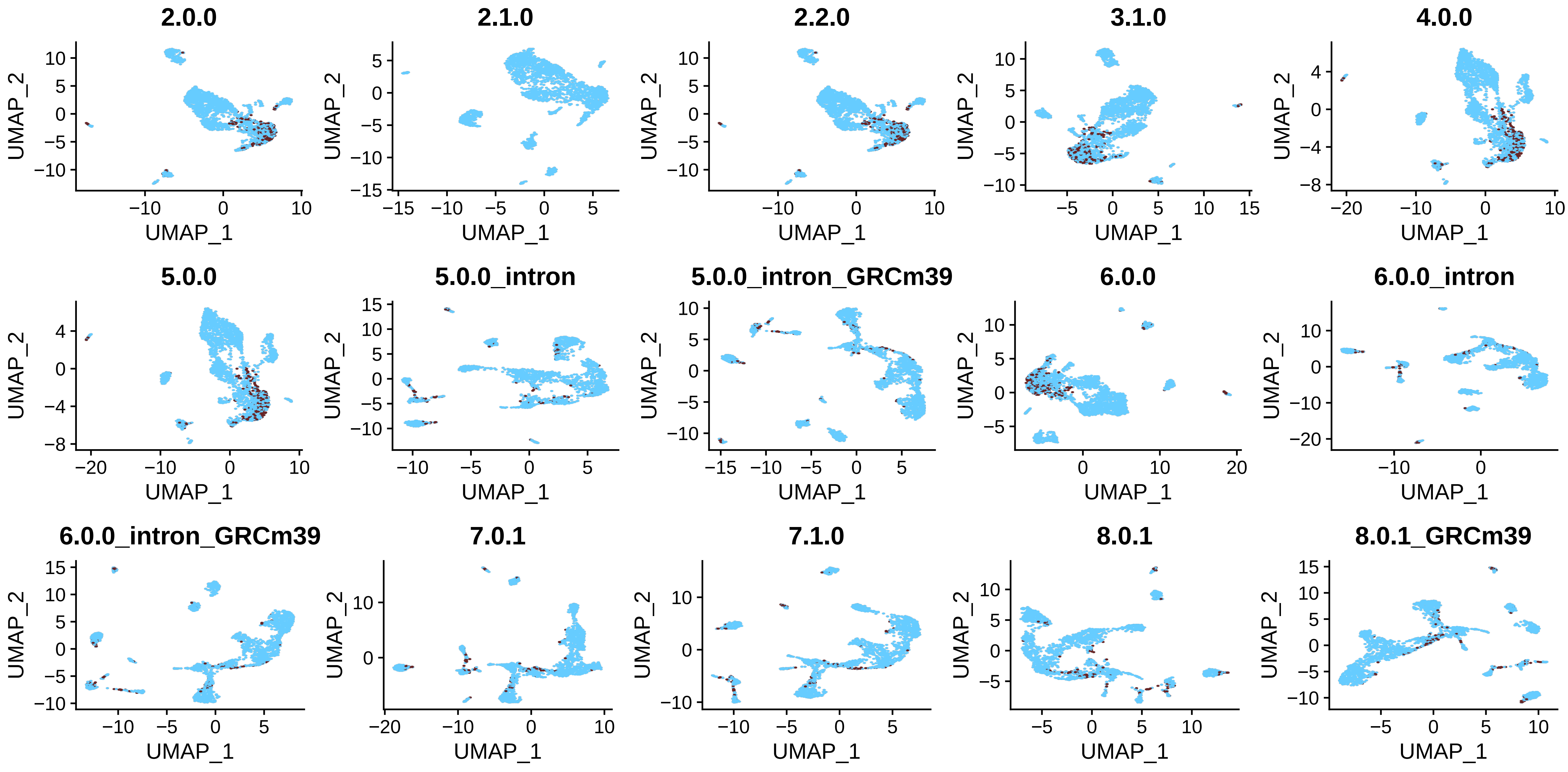

### Supplementary_Figure_4_nuclei_900.pdf

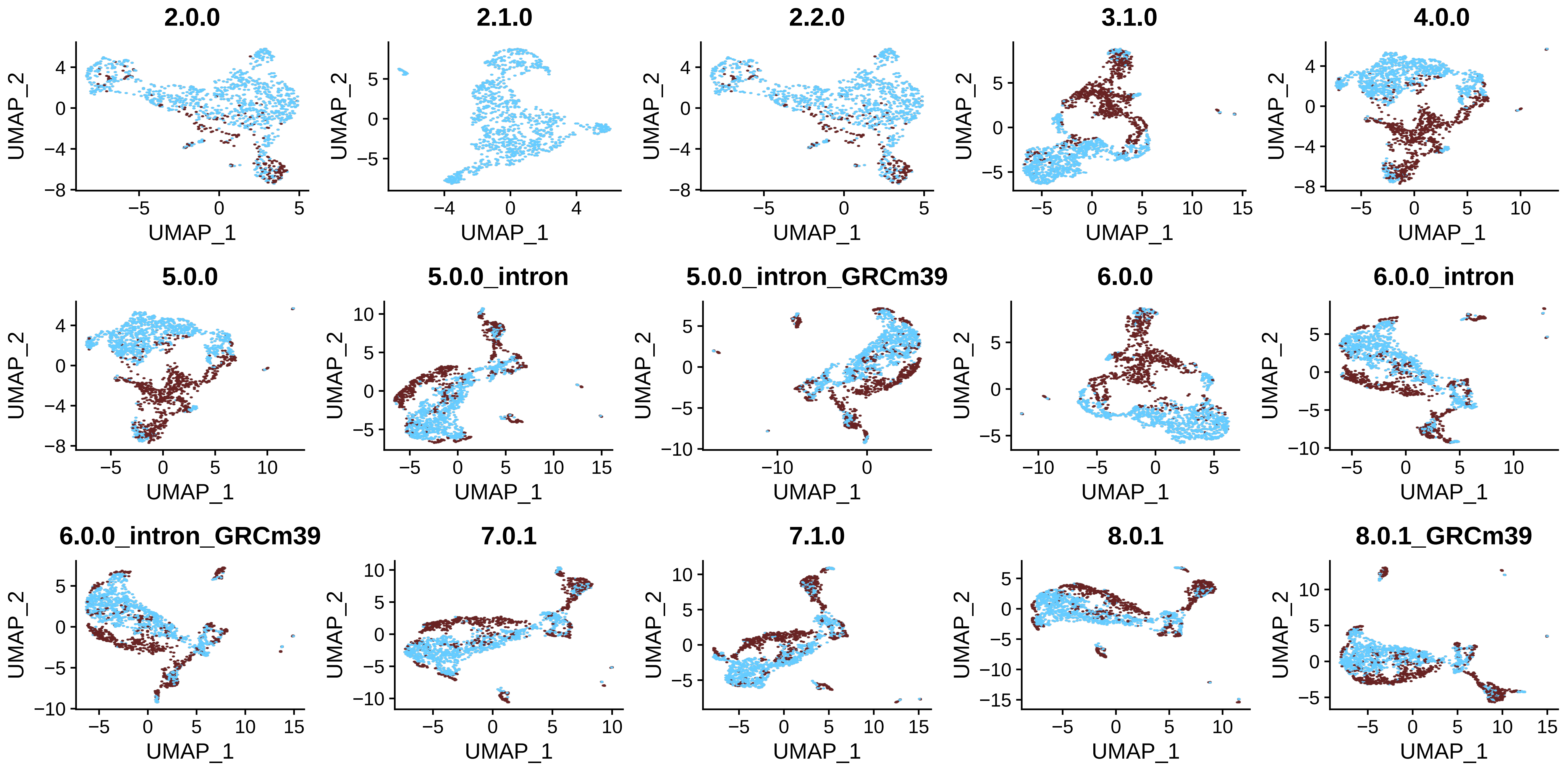

### Supplementary_Figure_4_pbmc4k.pdf

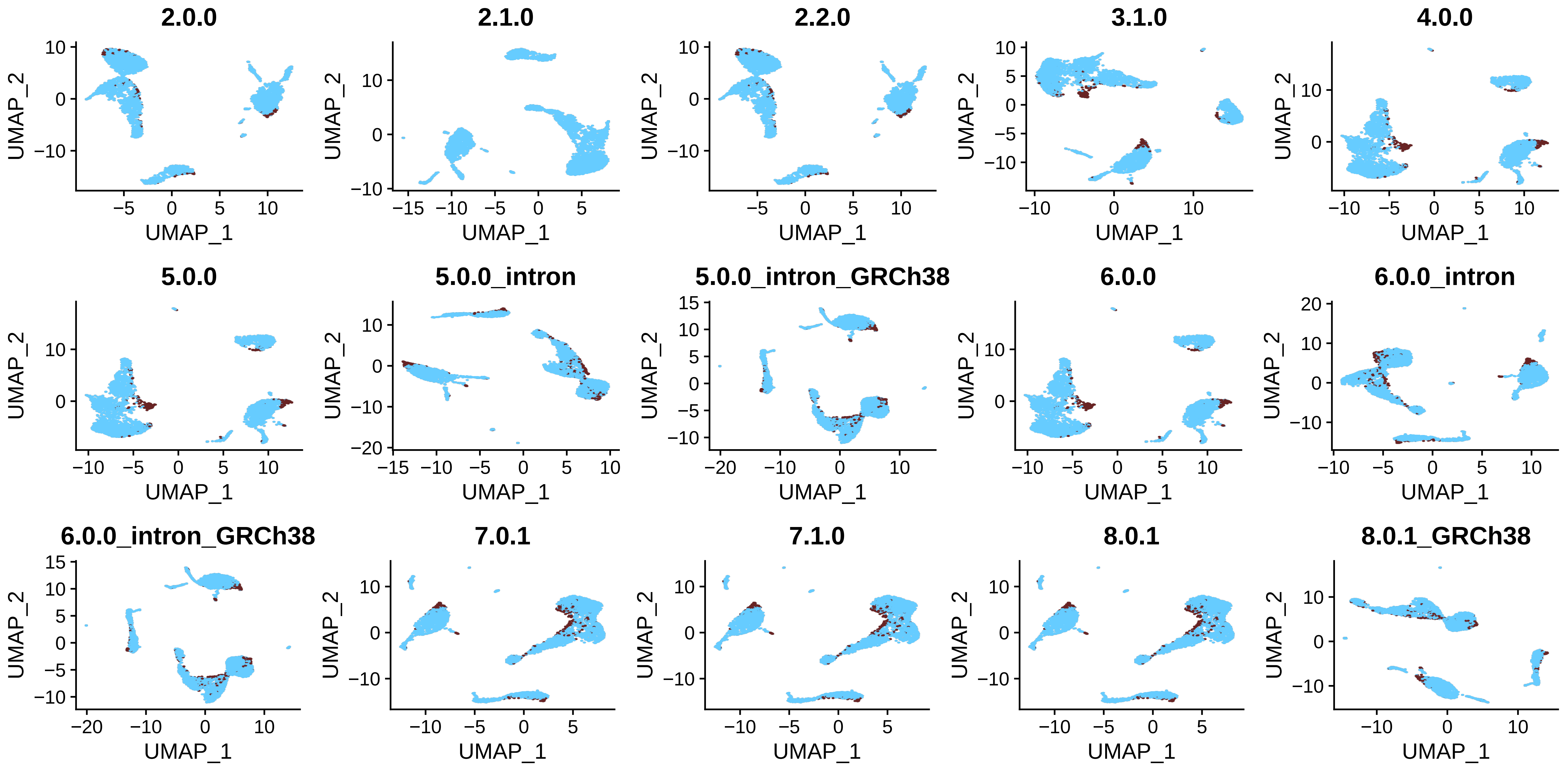

### Supplementary_Figure_4_t_3k.pdf

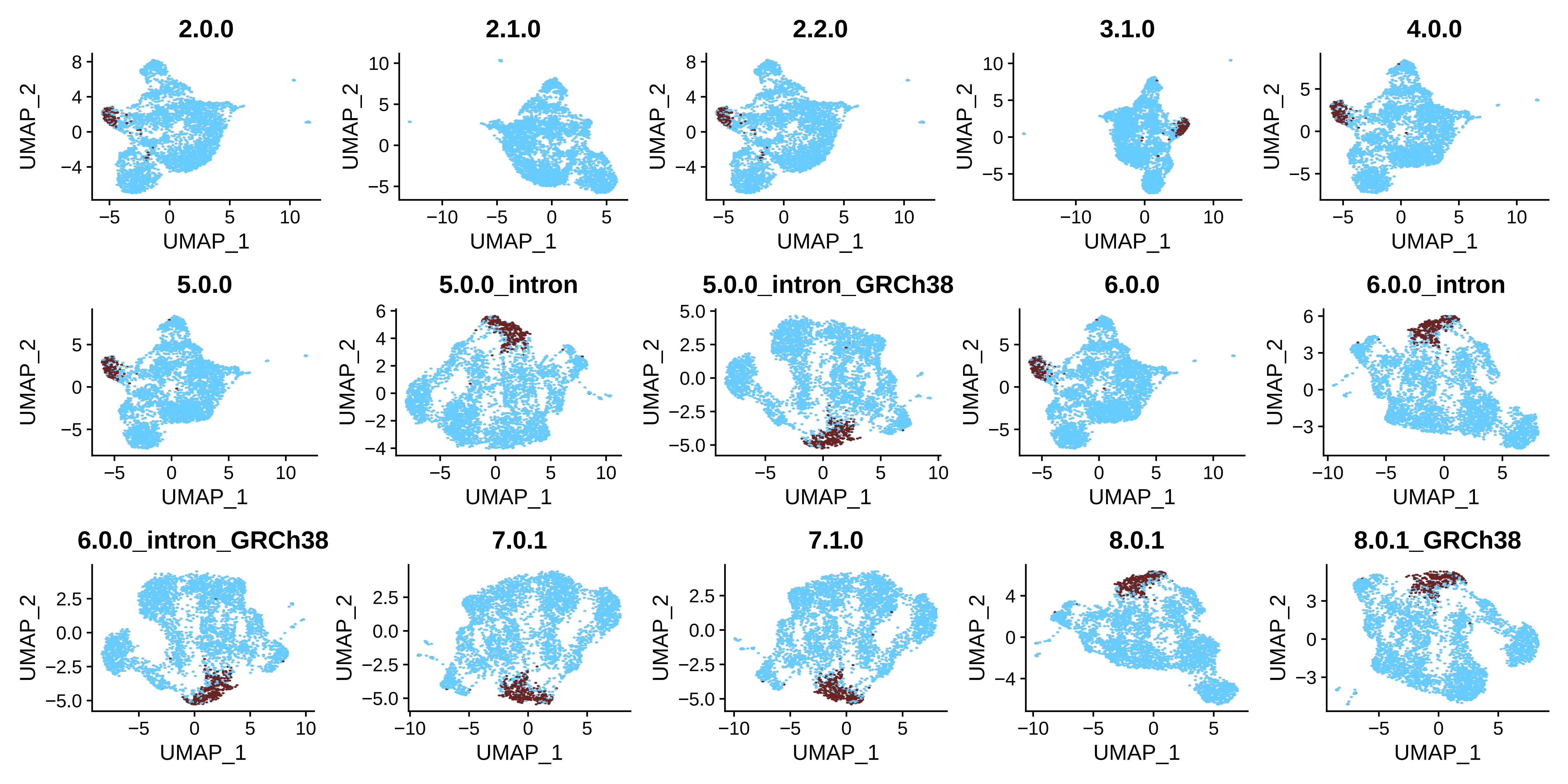

### Supplementary_Figure_4_t_4k.pdf

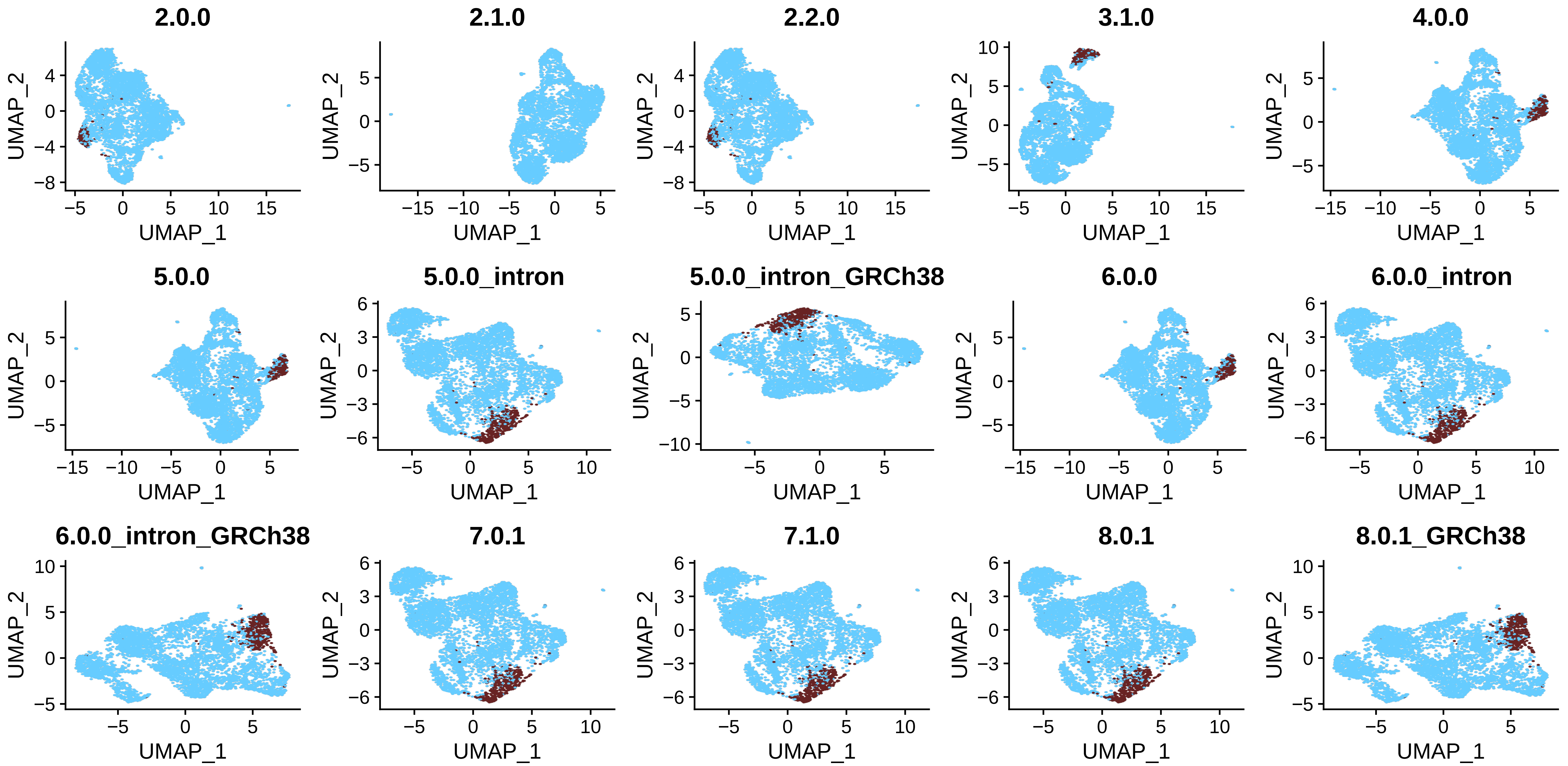

### Supplementary_Figure_5_fixed_neurons_6days_2000.pdf

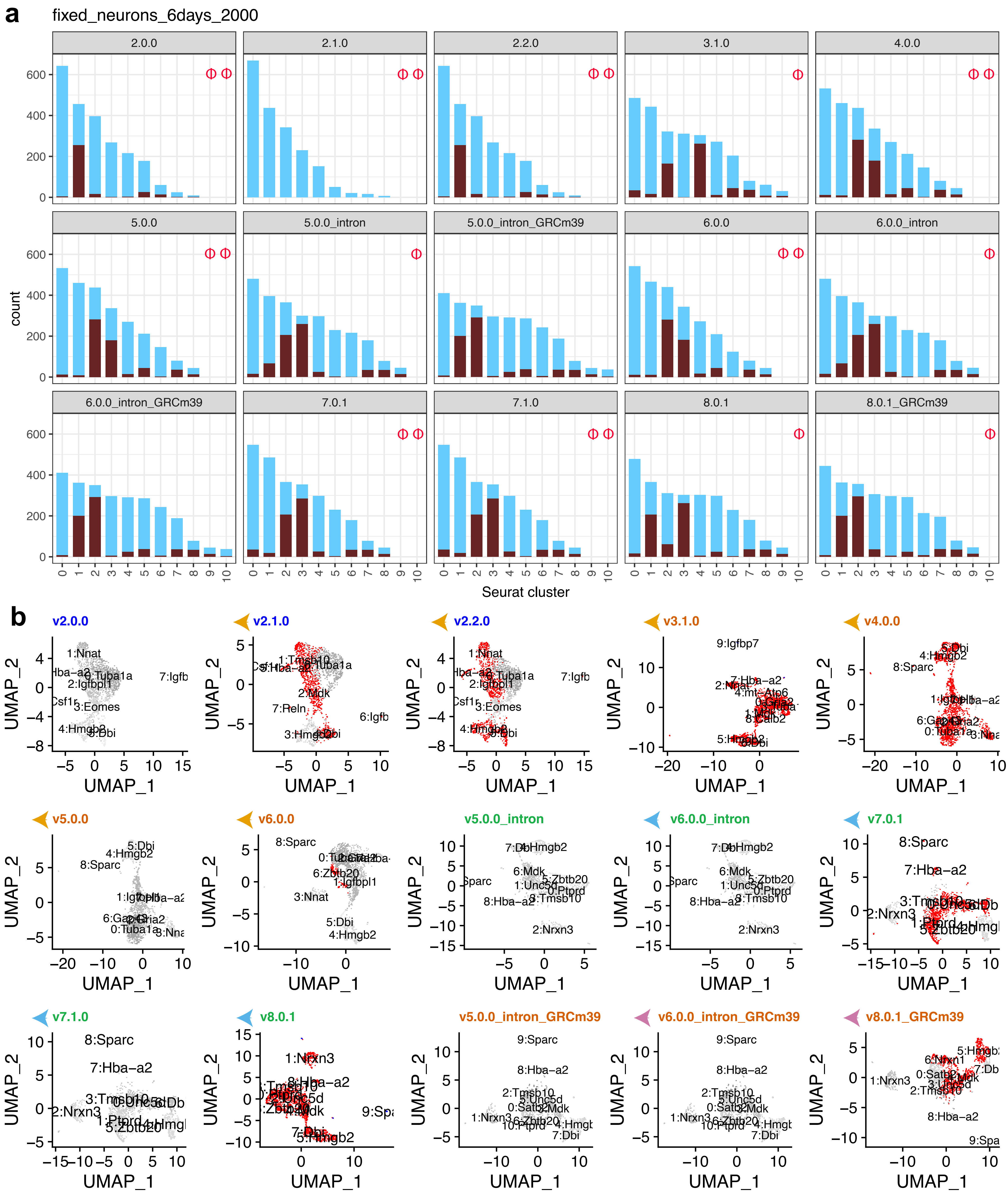

### Supplementary_Figure_5_fixed_neurons_2000.pdf

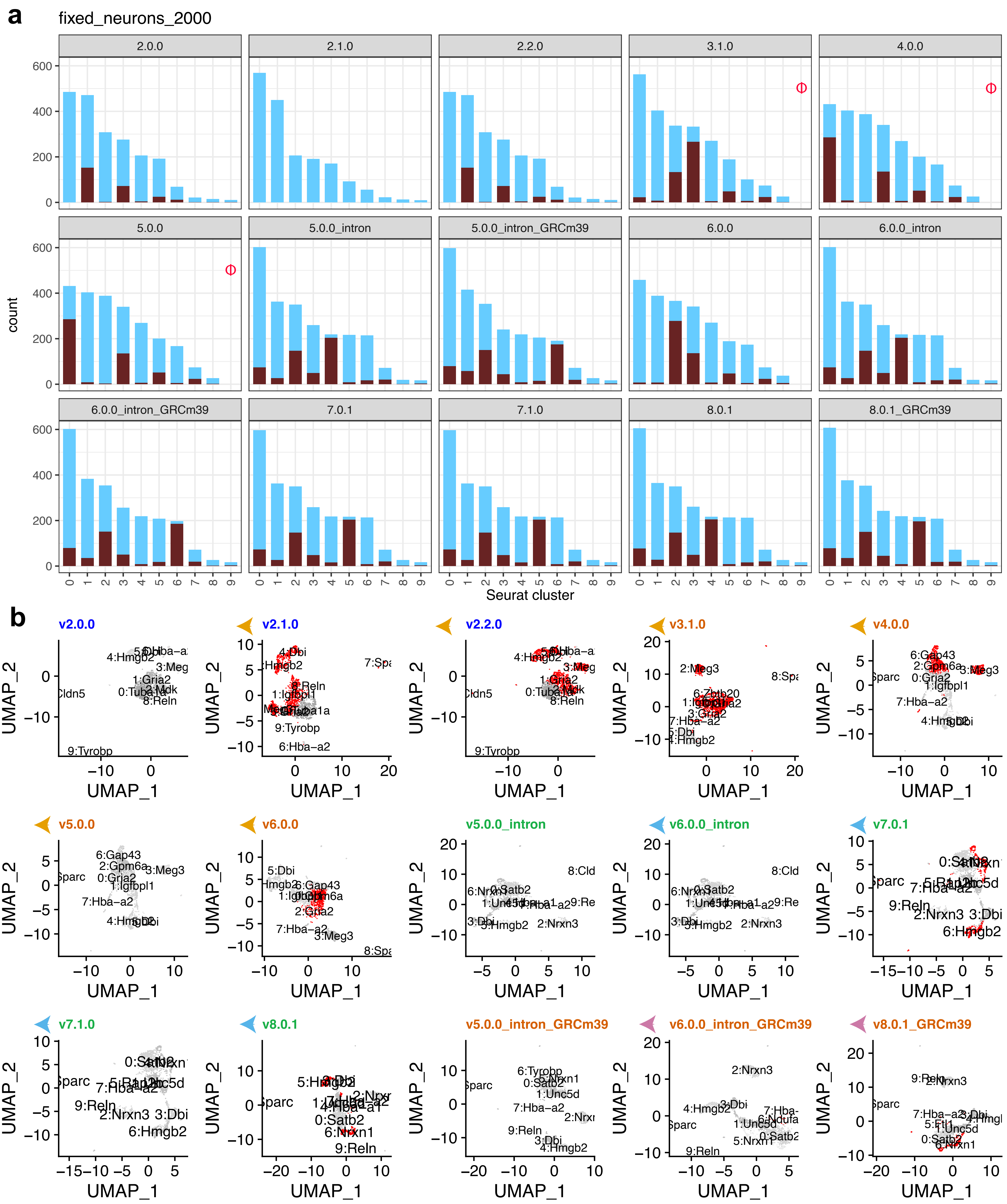

### Supplementary_Figure_5_GSE143607.pdf

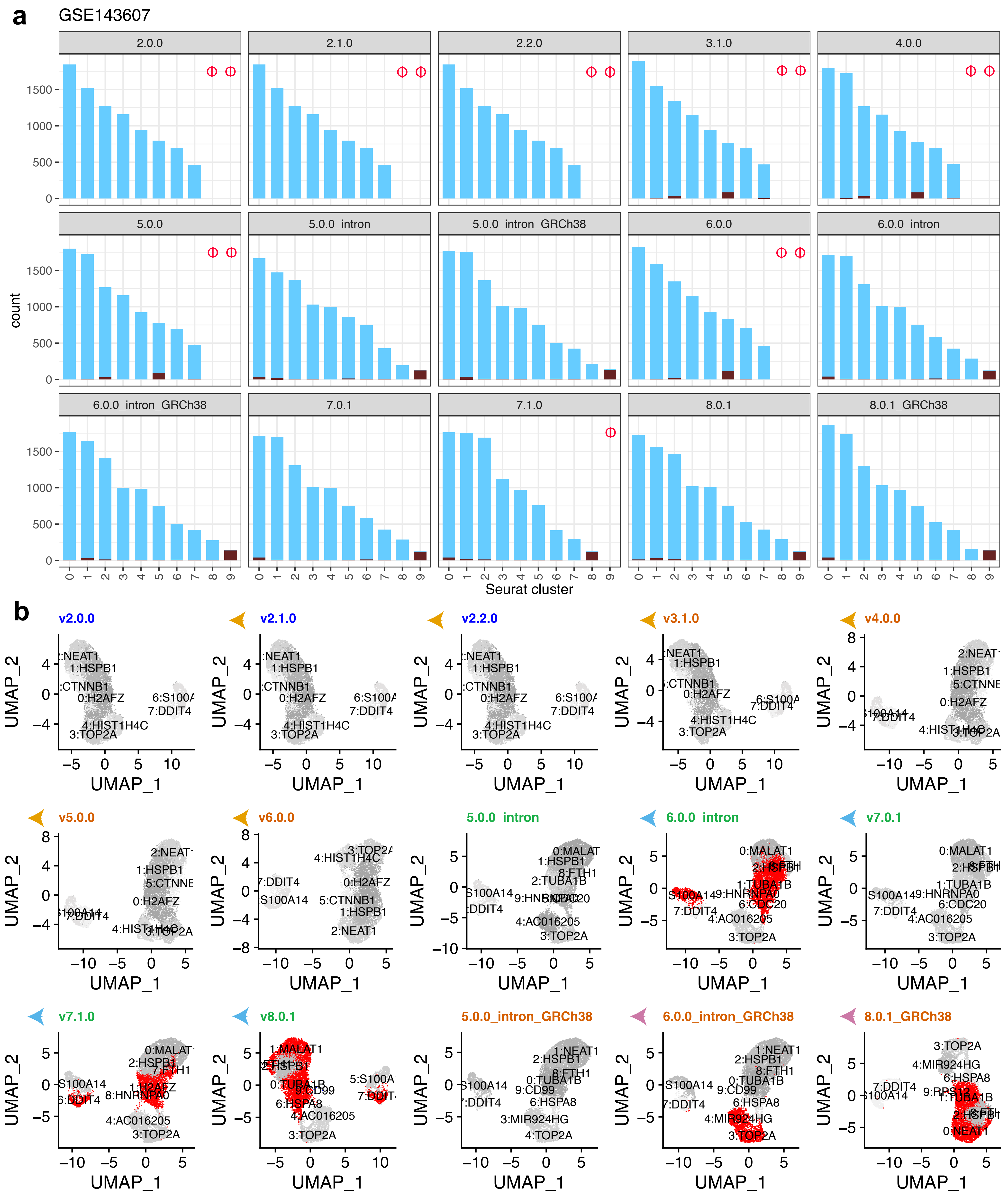

### Supplementary_Figure_5_neuron_9k.pdf

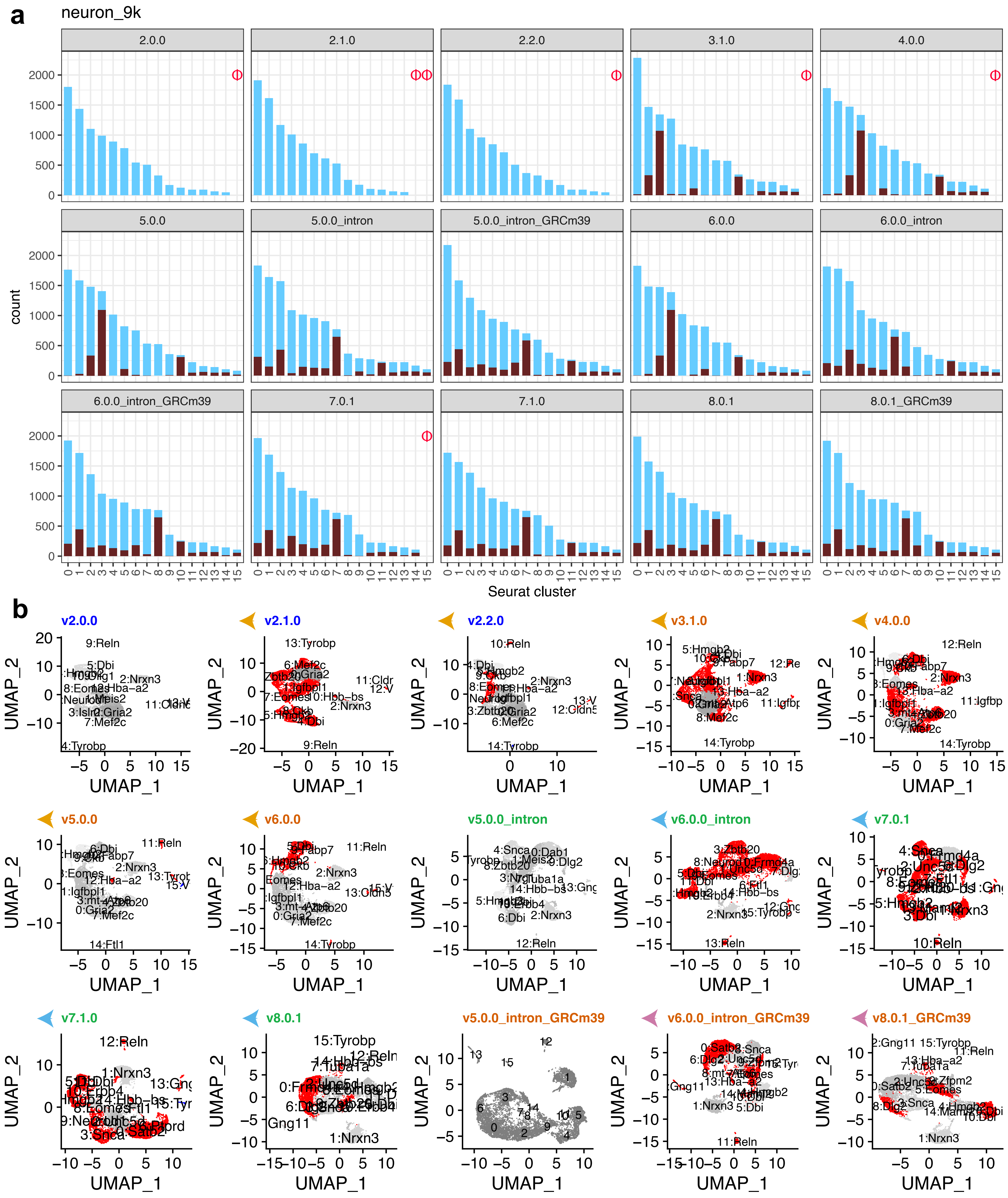

### Supplementary_Figure_5_neurons_900.pdf

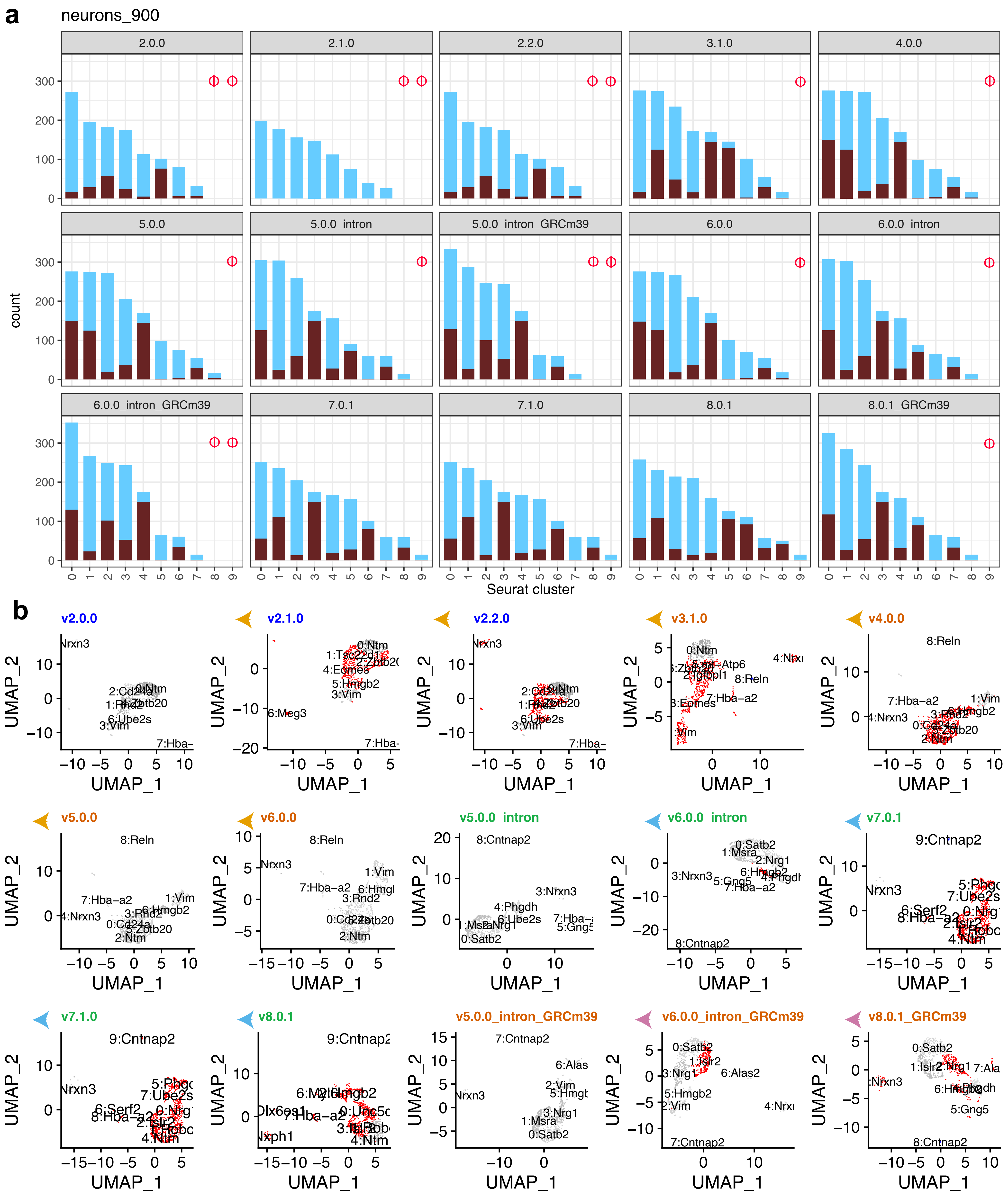

### Supplementary_Figure_5_neurons_2000.pdf

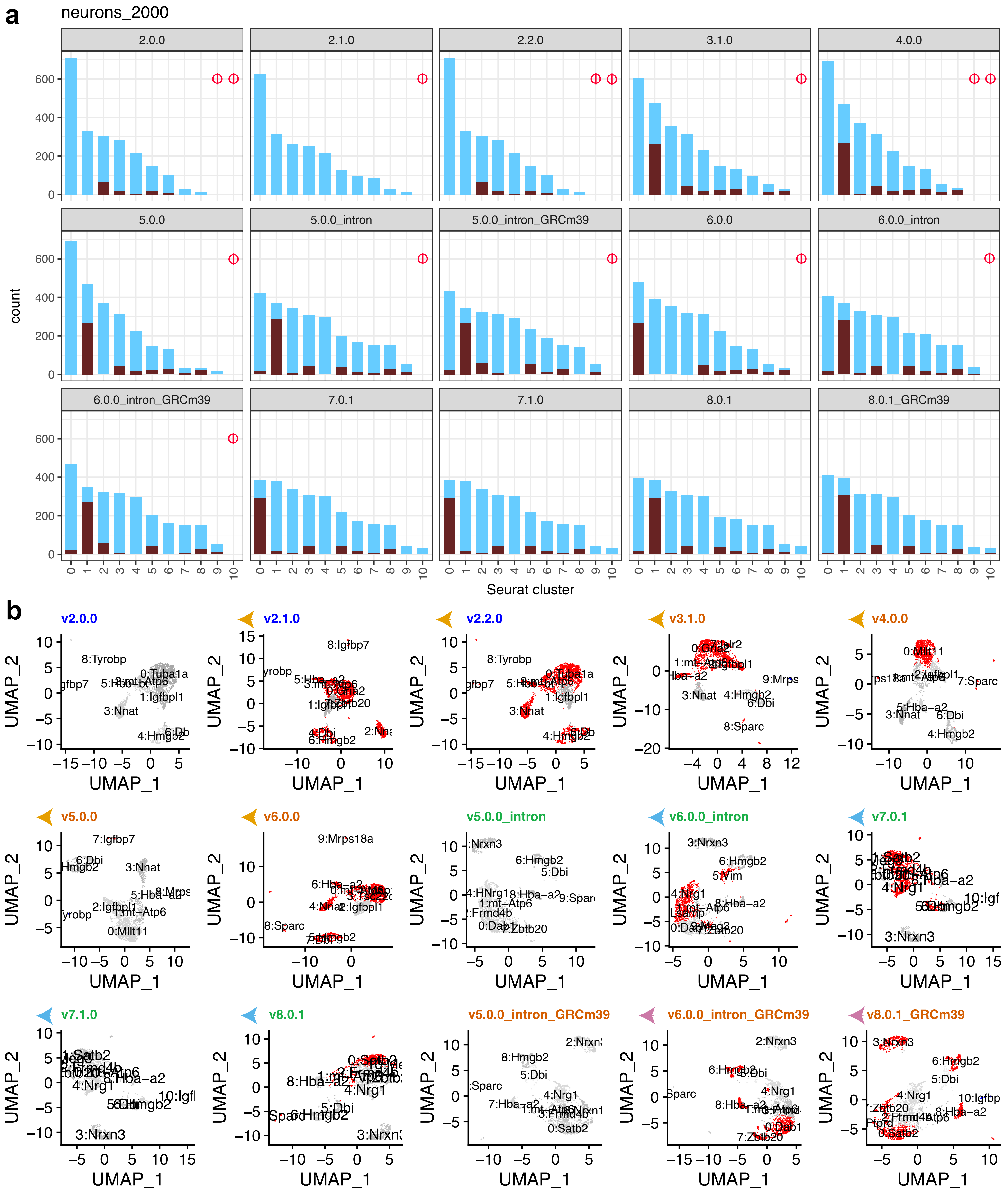

### Supplementary_Figure_5_nuclei_2k.pdf

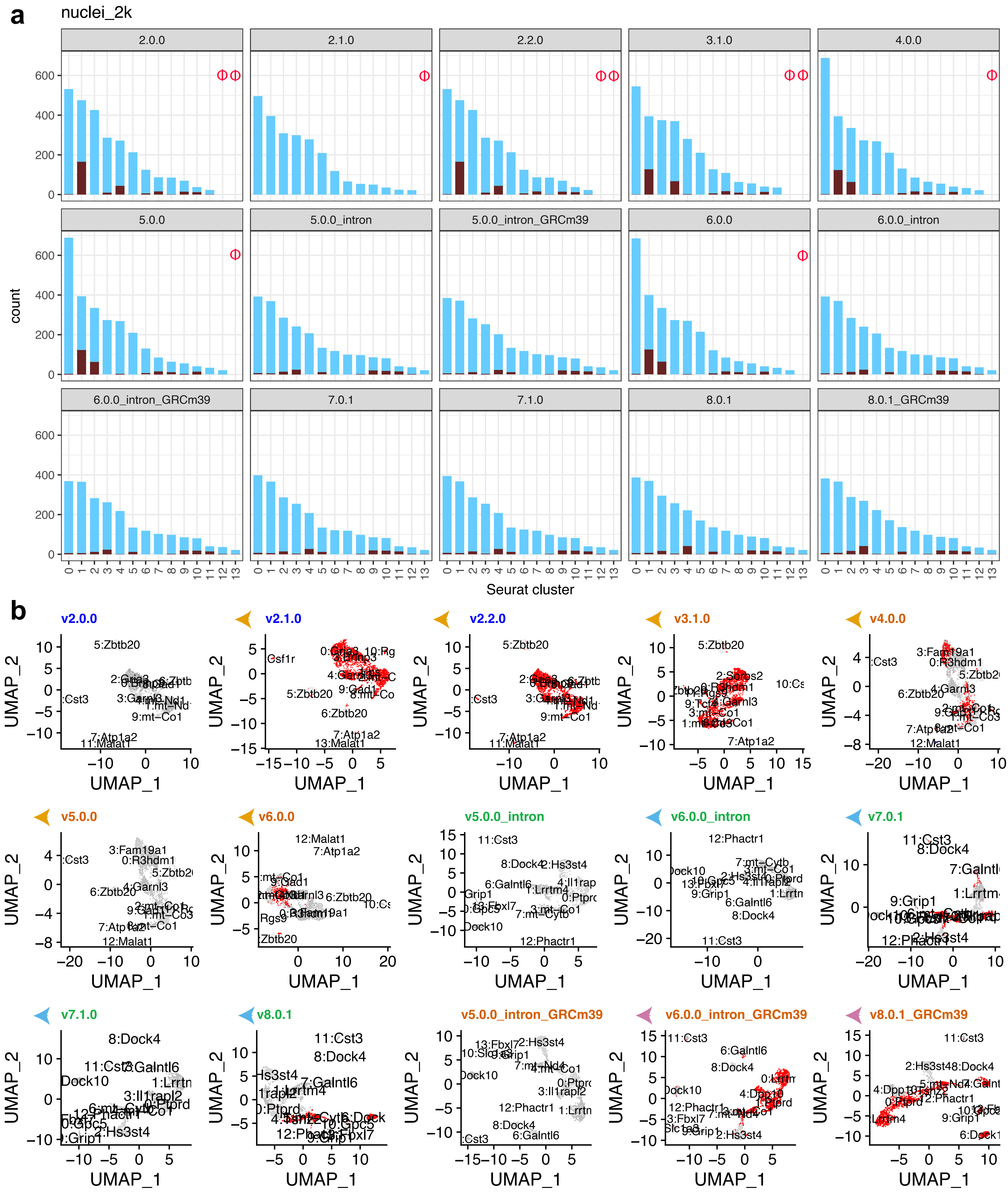

### Supplementary_Figure_5_nuclei_900.pdf

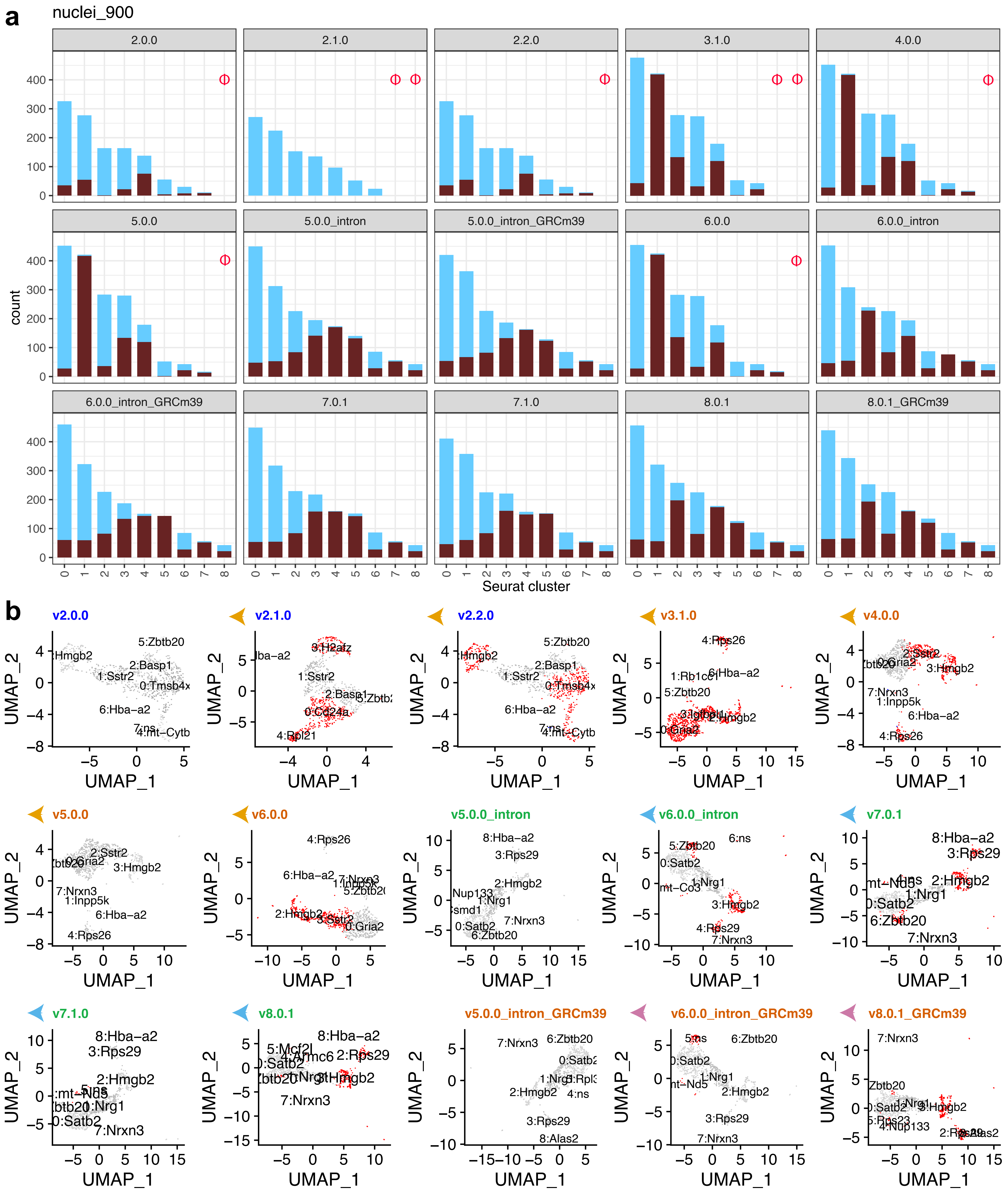

### Supplementary_Figure_5_pbmc4k.pdf

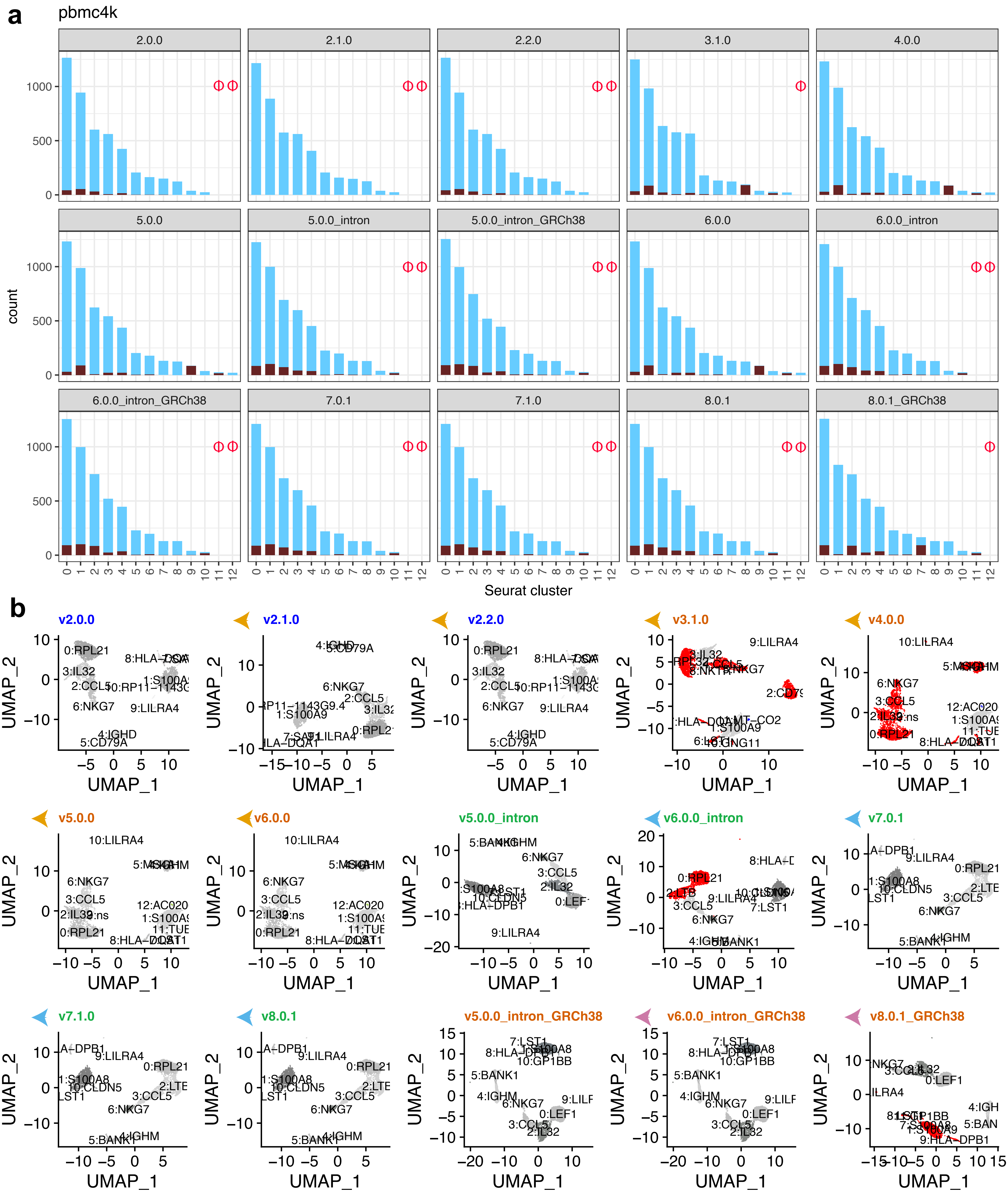

### Supplementary_Figure_5_t_3k.pdf

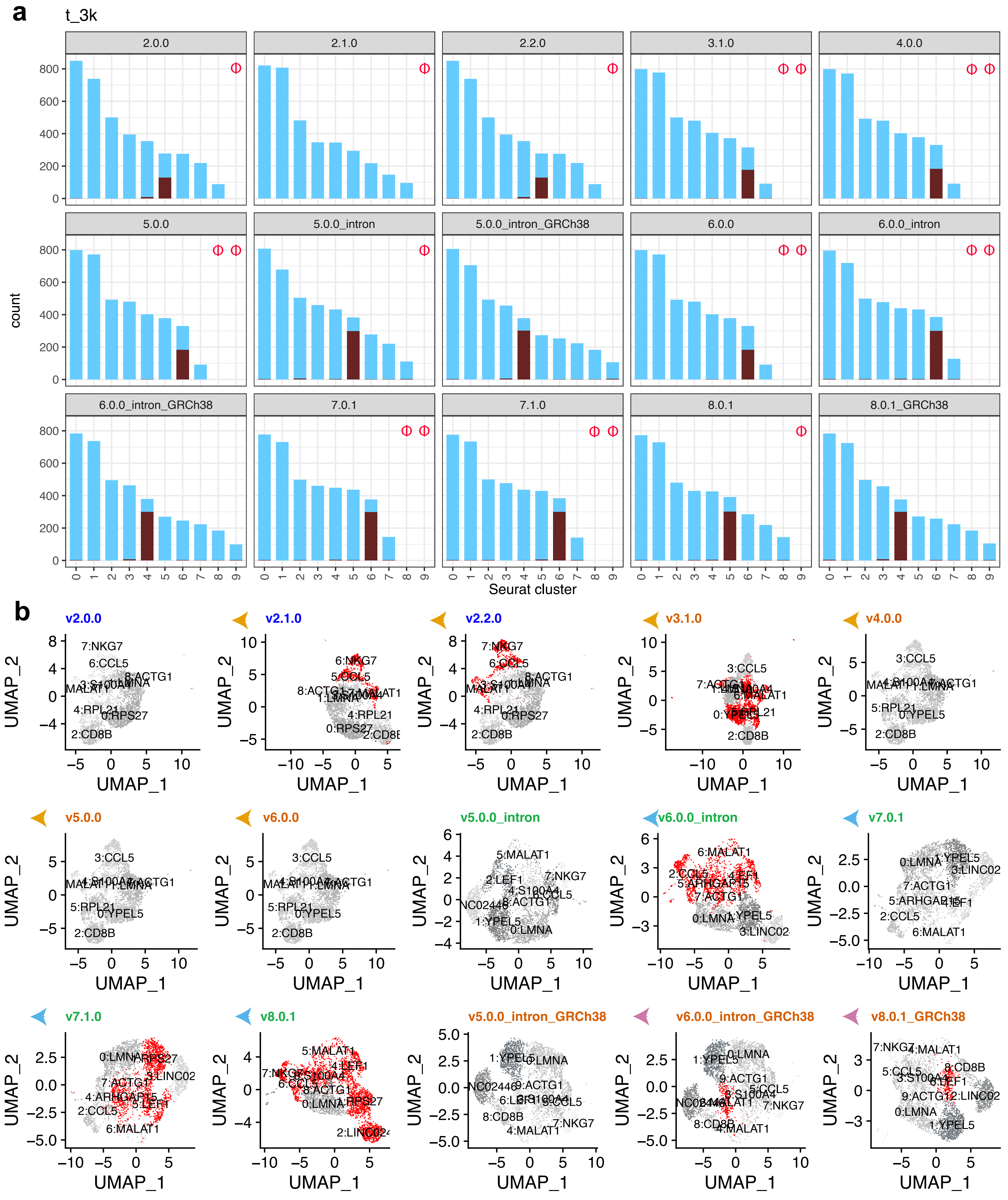

### Supplementary_Figure_5_t_4k.pdf

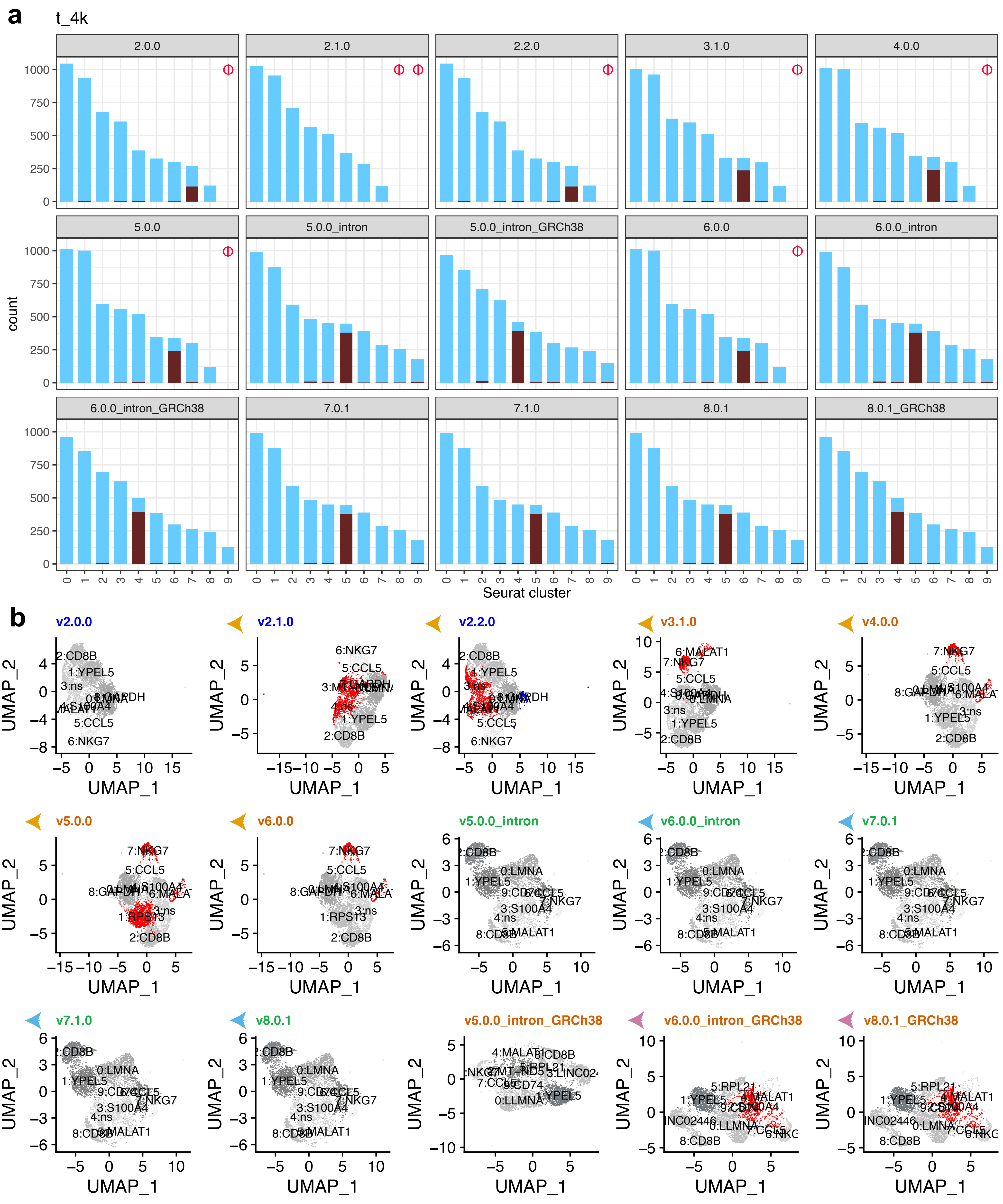
